# CodonBench: separating gene context, exon position, and variant-specific component in synonymous-variant classification

**DOI:** 10.64898/2026.08.12.744371

**Authors:** Yuanqing Liang, Weimin Zhu, Xiaoyong Pan, Huixian Li

## Abstract

**Background:** A synonymous variant leaves the protein unchanged, so a classifier that still separates pathogenic from benign examples must be reading something beyond the protein. Such variants test cleanly what a codon-level model adds to a protein-level one.

**Methods:** We assembled SynPath, a benchmark of ClinVar synonymous variants, and scored frozen representations under two splitting protocols. Under gene-held-out folds whole genes are withheld, and each coding-sequence model is compared against a position-matched control.

**Results:** The models were not trained for this task, so these numbers describe transfer, not designed performance. Random splitting flatters every representation. Once whole genes are held out, they fall into a narrow band near chance, the best at 0.58, and low-rank fine-tuning does not restore the random-split level. A feature that reads no sequence, only the distance from the variant to the nearest exon junction, reaches 0.84 under the same folds, above every neural representation tested. In a protein model the variant-specific component is zero by construction, because both alleles present the same input. In coding-sequence models only the response vector, the difference between the alternate and reference embeddings, keeps signal on unseen genes and beats its position-matched control.

**Conclusions:** Synonymous-variant pathogenicity is predictable mainly from gene identity and from position. Only a small part reflects the variant itself, and only in models that read the coding sequence. Benchmarks judged on random splits will overstate what these models have learned.

## Background

Synonymous variants change a codon without changing the amino acid it encodes. Because the protein is held fixed, a classifier performing above chance on this task must be reading a signal that lies outside the protein sequence. That property makes synonymous variants a natural setting in which to ask whether codon-level language models (cLMs) capture something that protein-level language models (pLMs) cannot (1,2). Both families span a range of architectures and training corpora (3–8), and several have been assessed on codon-level tasks directly (9,10).

The task is clean in principle but not in practice. Variants do not arrive uniformly from sequence space; they arrive attached to genes, and a gene carries its own label prevalence, its own nucleotide composition, and its own share of clinical attention. A classifier can therefore score well by recognising which gene a variant came from rather than what the variant does. Benchmarks in other domains have repeatedly shown that apparent performance can disappear once such structure is controlled (11–14), and benchmarks of genomic sequence models have begun to report split-dependent variation of this kind (15–17).

Two limits on what this design can establish should be stated at the outset. First, the protein context is held fixed within a variant, but is not matched across genes and positions, so a model that reads protein context retains room to score. Second, every model here is evaluated on a task it was not trained for, so a weak result is evidence about transfer to this task, and does not by itself refute a claim made about a different task.

An earlier version of this work was deposited as a preprint (10); it asked whether the apparent advantage of codon-level models on this task survives protocols that control for leakage, and it does not. The present version adds exon position as a source measured under the same folds as the model panel, and a control variant matched not only in window but at the same position of the same codon.

We decompose predictability into named sources. Three were anticipated — gene identity, sequence background (fixed by an identity rather than measured), and the variant-specific component. A fourth emerged from the data: a variant’s distance to the nearest exon junction, a quantity that needs neither a model nor the sequence; it proved to be the strongest of the four. We then tested whether any variant-specific component survives when gene identity and position are both controlled. synVep (18) scores synonymous variants specifically, and its features include splice-related distances, so the positional shortcut reported below is in principle available to tools built for this variant class.

## Results

The results are organised by conclusion rather than by protocol; Figure 1 summarises the design: how much apparent performance survives gene hold-out; the strongest single source, which reads no sequence; whether a variant-specific component survives with gene and position both controlled; and the bounds of the attribution. Supplementary tables are cited as Table S*n* and provided in Additional file 1.

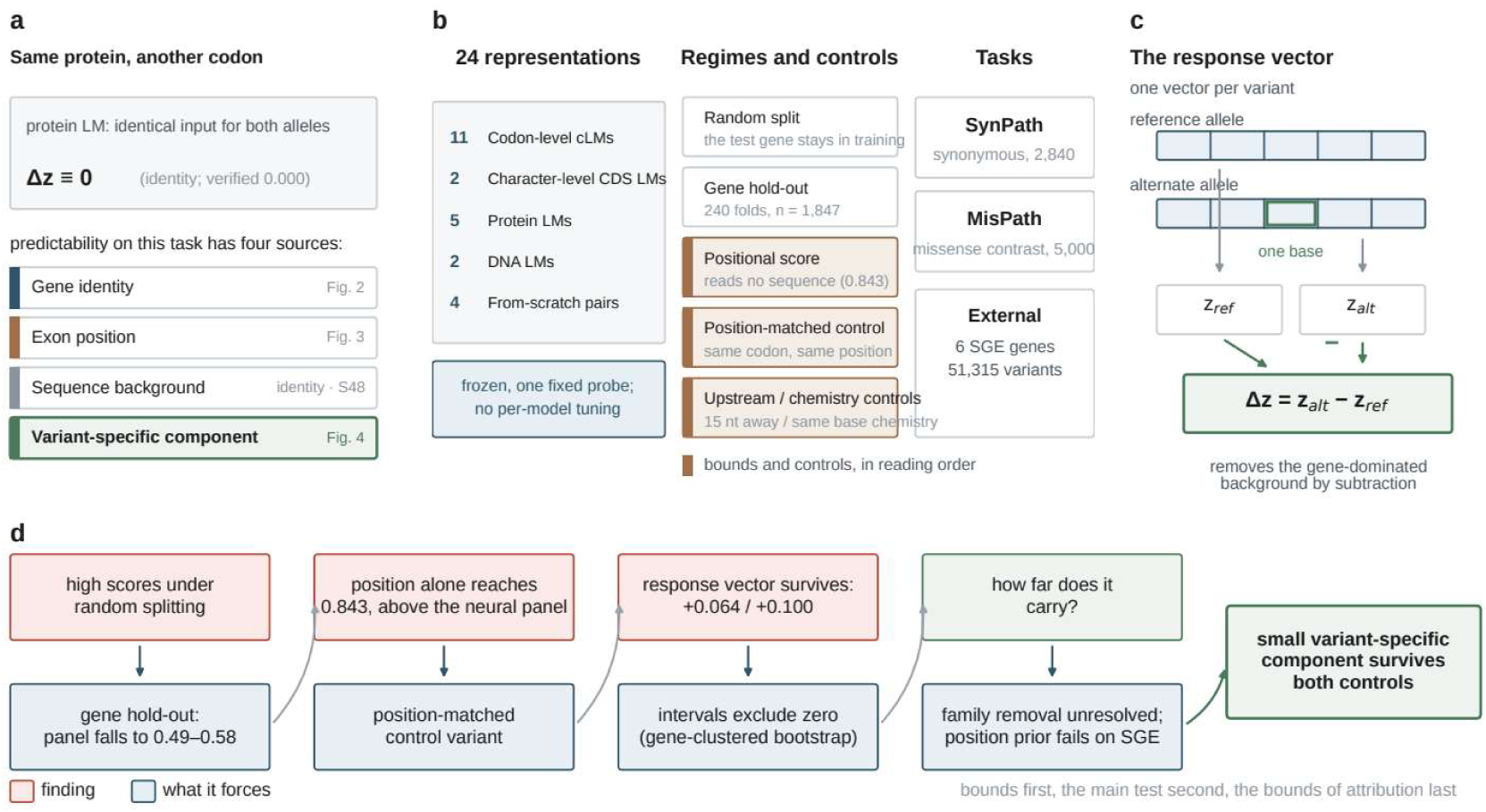
Four sources of predictability, the benchmark design, and the order of evidence. (a) A synonymous variant leaves the encoded protein unchanged, so a protein model receives an identical input string for both alleles and its response vector is exactly zero — an identity, verified numerically (0.000, zero-width interval; Table S48); predictability on this task decomposes into four sources, weighed in the figures indicated. (b) The benchmark: 24 frozen representations under one fixed probe; two splitting regimes (random split; gene hold-out over 240 folds, n = 1,847); a control ladder (a positional score reading no sequence, a position-matched control, and upstream and chemistry-matched controls); and the task set (SynPath, the MisPath contrast arm, and six external SGE genes). (c) The response vector, Δz = z_alt_ − z_ref_, one vector per variant: subtraction removes the gene-dominated background and leaves the model’s reaction to the substitution. (d) The order of evidence: bounds first — the panel collapses under gene hold-out, and position alone reaches 0.843, above every neural representation on held-out genes; the main test second — the variant-specific component survives the position-matched control (+0.064 and +0.100, intervals excluding zero); and the bounds of the attribution last — family removal is unresolved, and the position prior fails on measured function.

The benchmark contains 2,840 ClinVar synonymous variants drawn from 497 transcripts, with pathogenic and benign classes balanced by construction, so that the AUC (area under the ROC curve) is class-symmetric. Two properties of the stored records had to be resolved before any modelling (Methods, *Strand, frame, and exclusions*): the stored windows are already on the coding strand, and the reading frame is taken from the variant’s own position in the coding sequence, never chosen to make the two alleles agree, because a frame chosen that way would guarantee the agreement it is later used to verify. So fixed, 2,823 of the 2,840 variants resolve, and 2,821 of those translate to the same amino acid under both alleles. The leave-one-gene-out design covers 1,847 of the 2,840 variants, the remaining 993 sitting on transcripts too small to form a fold; the evaluated set is not a random sample of the whole (54.1% pathogenic against 42.3% in the remainder; Table S38).

### Gene hold-out collapses the whole panel to a band around chance

The panel’s collapse under gene hold-out is real, and not an artefact of how inputs are encoded, of how models are read out, or of how they are probed. We evaluated 24 frozen representations under two splitting regimes, using an identical probe in both: a logistic regression on standardised features, with standardisation fitted on the training folds only (Methods, *Splitting and probe*). Under random splitting most representations clear chance by a wide margin; under gene hold-out the entire panel contracts to a band around chance (Table S47).

Two features stand out (Figure 2a, b): the gap between the two regimes runs from 0.10 to 0.26 in AUC across the panel; and once the gap is closed the codon models occupy the same band as the protein models, with no representation exceeding 0.58 (Table S39).

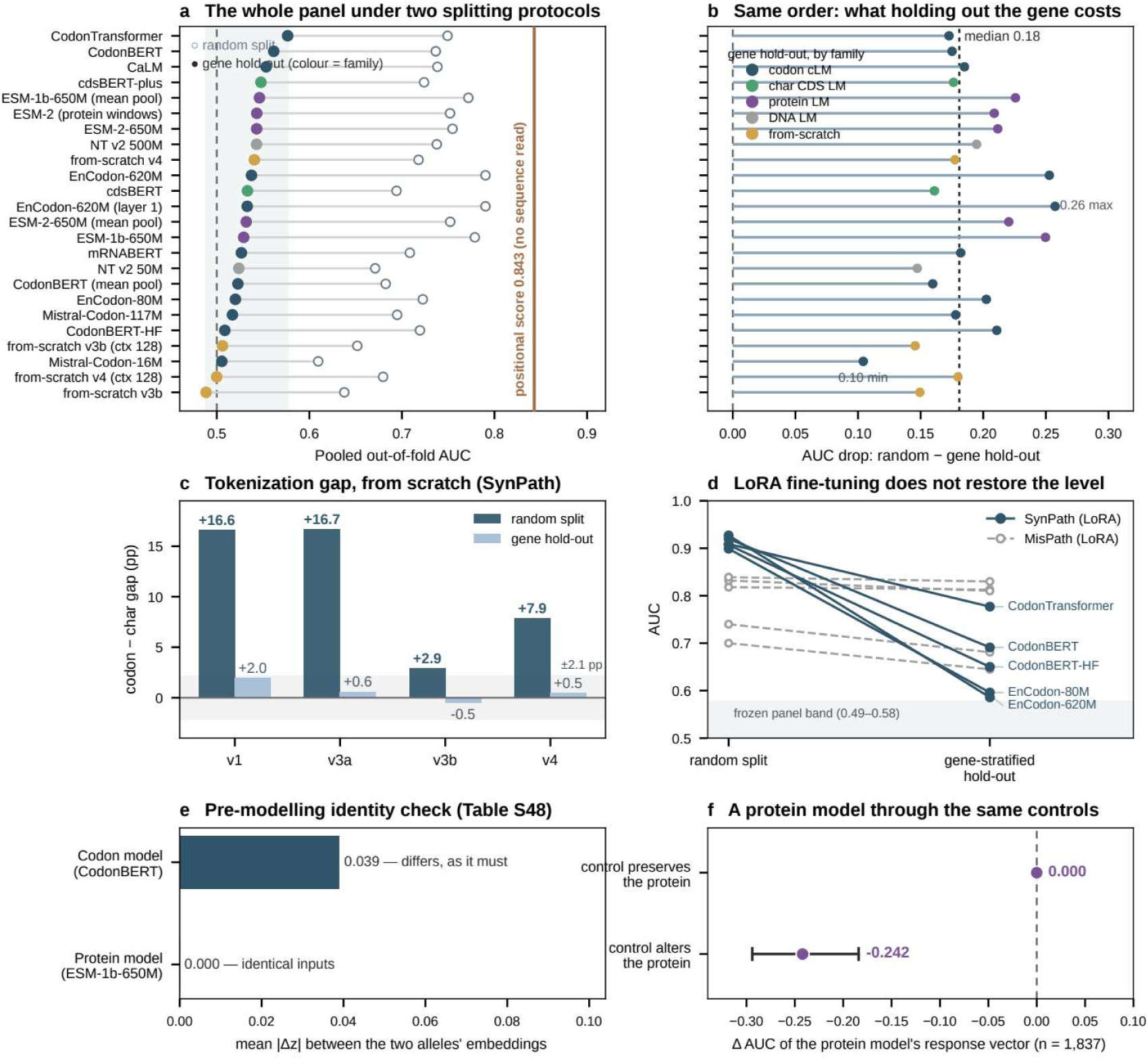
Gene hold-out collapses the whole representation panel. (a) Pooled out-of-fold AUC for 24 frozen representations under random splitting (open markers) and gene hold-out (filled markers), ordered by the gene-held-out value; the dashed line marks chance and the solid line the positional score, which reads no sequence. (b) The drop in AUC between the two protocols for each representation; the dotted line marks the median. (c) The gap between codon and character tokenization, trained from scratch, under the two protocols, for four training configurations that differ in corpus size, number of epochs and parameter count (Tables S6 and S15). (d) Five codon language models fine-tuned with low-rank adaptation, on the synonymous (solid) and missense (dashed) arms, under random splitting and gene-stratified hold-out (Table S9); the shaded band marks the frozen panel’s hold-out range. (e) Mean absolute difference between the reference and alternate embeddings: a codon model must differ and a protein model must not, because its two alleles are the same input (Table S48). (f) The same control comparison as in Figure 4, applied to that protein model: against a control that preserves the protein the difference is exactly zero with an interval of zero width, and against a control that alters the protein it is −0.242 (95% CI −0.294 to −0.184), n = 1,837 (Table S48).

#### Can the collapse be blamed on the encoding?

We first tested tokenization, codon by codon against character by character, trained from scratch under identical protocols (Methods, *Sequence inputs*). Under random splitting the codon tokenization carries a clear advantage, of up to nearly seventeen percentage points; under gene hold-out that advantage disappears, and the residual differences fall within about two percentage points and do not agree in sign (Figure 2c; Table S6, against Table S15).

#### Can it be blamed on the readout?

The second alternative is to let the model fit end to end. Low-rank adaptation gives the largest fitting freedom we tested, and under gene-stratified splits it does not restore the advantage: the drops are on average larger than the frozen panel’s, and although the fine-tuned models land above every frozen representation in the same regime — so a frozen read-out overstates how little these models carry — they remain far below what the same models reach under random splitting. The comparison rests on gene-stratified splits over three seeds, so it is not same-protocol (Table S9 and Figure 2d).

#### Can it be blamed on the probe?

The difference between a linear and a nonlinear probe on this task is not a property of the model but of the probe’s implementation. Decomposing one such gain step by step, no single configuration choice produces it (Figure 5c; Table S22), and across models the size of that step is model-conditional, in one case negative (Figure 5d; Table S23).

#### What follows from the input, and what does not

A synonymous variant leaves the protein unchanged, so a protein language model receives the same character string for the reference and the alternate allele; the two embeddings must therefore be identical, and their difference must carry no information about the variant. We verified this numerically (Figure 2e; Table S48) and carried the protein model through the same control comparison applied to the codon models below: against a control that leaves the protein untouched, the protein model’s response vector differs by exactly zero, with a bootstrap interval of zero width; against a control that does change the protein, the same arm differs by −*0.242* (95% CI −*0.294* to −*0.184*; Figure 2f). The consequence is that every point of apparent performance a protein model shows on this task originates in sequence background. That these models also show the largest change between regimes is an observation rather than a consequence of that identity: having the highest apparent performance under random splitting, they have furthest to move.

#### Does either family collapse less than the other?

Eight paired comparisons under gene hold-out on a shared test set separate none from zero (Table S49), and the intervals are wide, with a half-width near 0.046, because the effective unit of replication is the gene rather than the variant (Methods, *Statistical inference*): this dataset does not resolve a family-level difference, which is not the same as establishing that no difference exists.

### The strongest single source is position, and it reads no sequence

In SynPath, more than half of the pathogenic synonymous variants sit on the first or the last base of an exon, and almost none of the benign ones do (Table S50; Figure 3b). The excess appears at both ends of exons rather than at one end preferentially, the exons involved are of ordinary length, and the effect is spread over 248 of 495 genes rather than concentrated in a few. The observation is strong enough to be used directly as a classifier: taking the distance from each variant to the nearest exon junction, negated, as the score (Methods, *Positional baselines*) — no model fitted, no sequence read — reaches an AUC of 0.843 under the same folds and the same evaluation as the panel above, above every neural representation in Table S47 (Figure 3d). The enrichment is biologically plausible: exon boundaries are where the sequence elements that direct splicing are read (19,20), and synonymous changes there can alter splicing rather than protein sequence (21–23).

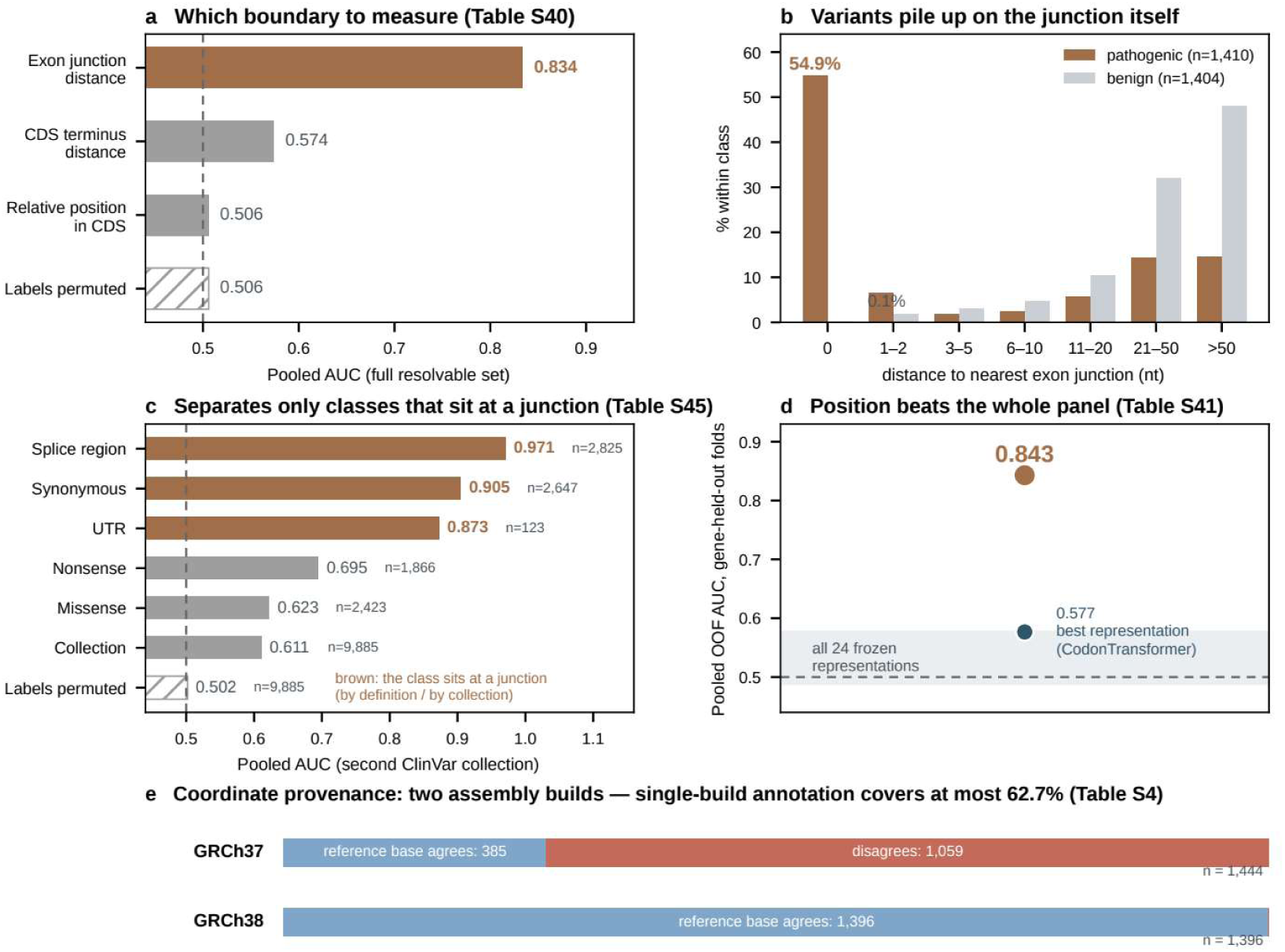
Position alone outperforms every neural representation under gene hold-out. (a) Pooled AUC for three positional quantities on the full resolvable set, with labels permuted as a control; the dashed line marks chance. The distance to the nearest exon junction needs neither a model nor a sequence, while the distance to the coding-sequence termini, which an earlier audit used to set splicing aside, barely separates the classes. (b) Distance of each variant to the nearest exon junction, as a percentage within each class: pathogenic variants accumulate on the junction itself, benign variants lie far from it. (c) The same score applied across consequence classes: it orders the classes that sit at a junction, splice-region ones by definition and synonymous ones through the reporting pattern above, and it orders missense substitutions only weakly, though above chance (Table S45). (d) The junction distance alone under the LOGO-240 evaluation folds, against the best representation under the same folds (Table S41). (e) Coordinate provenance: the 2,840 records split by the assembly build they are annotated on, and within each build by whether the recorded reference base matches the GRCh38 reference at the recorded position; every mismatch falls in the GRCh37 half, which bounds any coordinate-based annotation of this set (Methods, Coordinate provenance; Table S4).

Table S40 also settles which boundary should be measured. The distance to the nearest CDS terminus, which is what an earlier analysis used when arguing that splicing confounding could be set aside, reaches only 0.534. A coding sequence spans several coding exons, so its two termini are not exon junctions and the two quantities are not interchangeable.

#### How specific is the shortcut?

We applied the same quantity, with no model fitted and no sequence read, to a second ClinVar collection that mixes several consequence classes (Methods, *Positional baselines*). Position separates a class where that class sits at a junction: splice-region variants do so by definition; synonymous variants do so through how they were collected; missense substitutions carry no such constraint and are ordered only weakly, though above chance, against 0.50 for permuted labels (Table S45). The score therefore tracks proximity to a junction, not some generic property of ClinVar. One caution: the junction distance is computed from transcript-structure annotation, which lies outside the window the representations are given, so it is used here to locate where predictability comes from, not to compete with the models on equal terms.

Whether what the positional arm captures is independent of predicted splice disruption cannot be established here: the variant set mixes genome builds (1,444 records on GRCh37, 1,396 on GRCh38; at 1,059 recorded positions the recorded reference base does not match the GRCh38 reference), so most records cannot be located on the genome the predictor was run against (Methods, *Coordinate provenance*; Table S4). An earlier estimate that 0.007% of records score above the conventional DS ≥ 0.2 threshold came from this broken join and is withdrawn. Whether the enrichment reflects splice biology, the way these records came to be reported, or both, therefore stays open; the separation above never depended on it.

### A directional variant-specific component survives both controls

Absolute embeddings are dominated by gene identity, so to ask whether a model reads anything specific to the substitution, that background has to be removed first. The removal is also the standard scoring move in variant effect prediction (24–27): both alleles go through the frozen model, and we subtract one embedding from the other. We call this difference the response vector, one vector per variant. The control has to match the real substitution in position (Methods, *Controls*): a different synonymous codon at the variant’s own position, the substituted base at the same within-codon position as the real substitution.

The first thing to establish is how many of the controls were actually built: where the amino acid has only two codons and the reference and alternate codon are those two, no third synonymous codon exists and the control response vector is the zero vector, which carries no signal and pulls the control arm towards chance. This group is not small: it is 630 of the 1,839 evaluated records (34.3%; Table S43). On the 1,209 records whose control was actually built, the response vector exceeds the position-matched control by +*0.064* in one model and +*0.100* in the other, both intervals excluding zero (Table S55). Over the full evaluated set the same difference reads +*0.138* and +*0.145*, inflated by the never-built controls (Table S42). This comparison, the response vector against the position-matched control, is the single pre-specified test of the study; the strata and decompositions that follow are exploratory.

#### What the component is, and what it is not

Normalising the response vector to unit length leaves the difference essentially where it was, +*0.064* (0.019 to 0.110) and +*0.103* (0.058 to 0.147), so what discriminates is the direction of the model’s reaction and not its size; and adding the response vector on top of position, of simple sequence features, and of the magnitude leaves the difference between +*0.063* and +*0.091* across the two models, every interval excluding zero (Table S58), so it is no re-encoding of something a simple feature already captures. A harsher variant projects the response vector into the space orthogonal to those features and does remove the difference; a projection that drives both arms below chance cannot discriminate between them, so it is reported and not interpreted (Table S55). The component is also not spread evenly over substitutions: where the real substitution moves the codon to a rarer synonym, the difference is resolved in both models, +*0.123* (0.057 to 0.191) and +*0.101* (0.040 to 0.161); where the codon becomes more common it is resolved in one model only; and where the frequency barely changes it is unresolved in both (Table S56). Pathogenic synonymous variants in SynPath do preferentially replace common codons with rare ones, the mean change in synonymous-codon frequency being −*0.053* for the real substitutions against −*0.010* for the control variants, so part of what the response vector reads is a codon-usage signal that the substitution itself carries.

The two arms are not comparable on every simple feature: 76% of the real substitutions are transitions against 15% of the control variants (Table S57), which bounds what the comparison can claim. Two further controls behave as expected: a label-permuted response vector returns to chance, and the protein model gives a difference of exactly zero with an interval of zero width. The component is small, it transfers to genes the model has never seen, and it is invisible in the absolute embedding, where gene identity dominates (Figure 4).

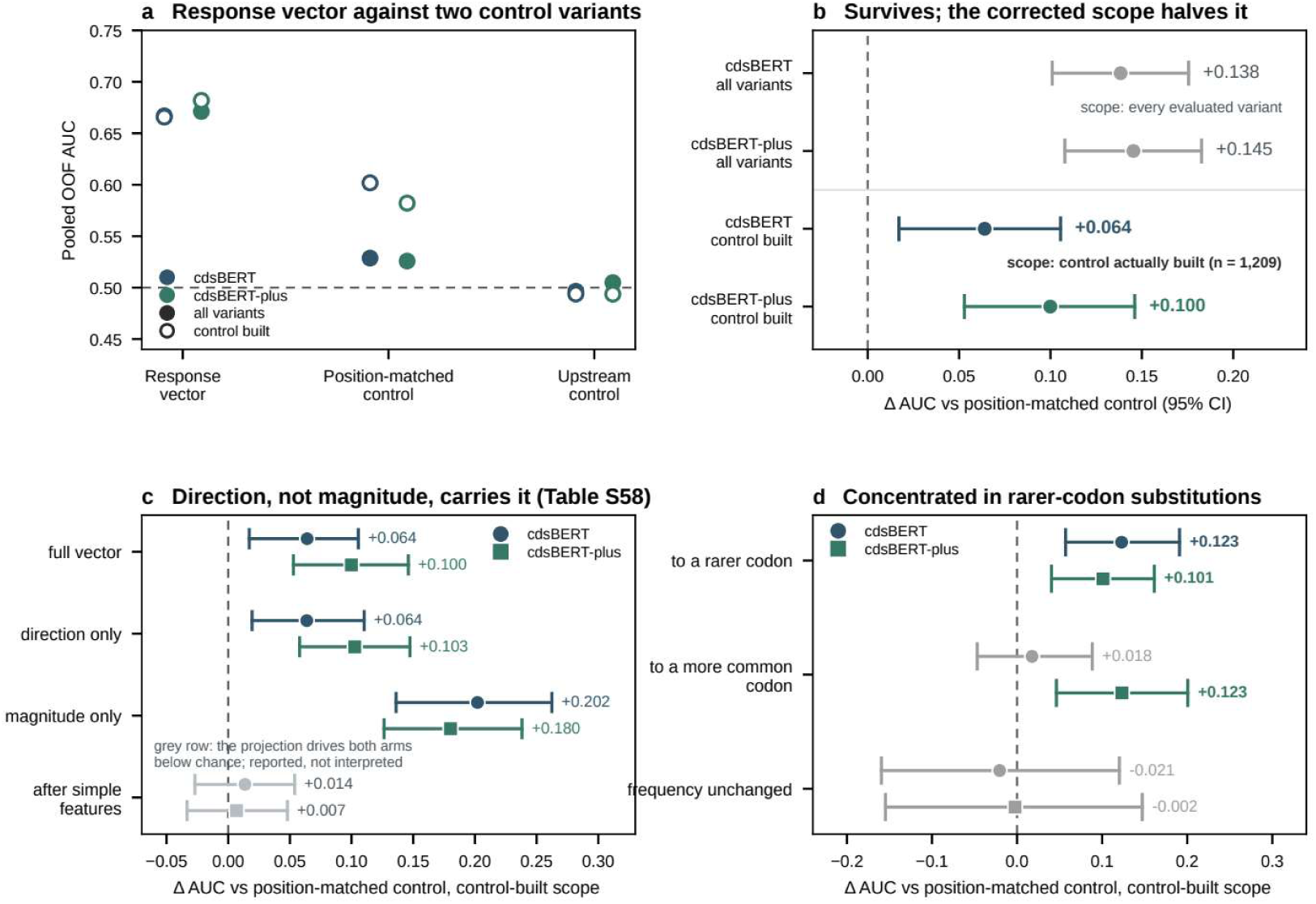
The variant-specific component and what it is made of. (a) Pooled out-of-fold AUC for the response vector, the position-matched control and the upstream control in the two models that support this analysis, both of which read the coding sequence character by character; filled markers give every evaluated variant and open markers the subset whose position-matched control was actually built; the dashed line marks chance. (b) The difference against the position-matched control under both scopes in both models, with intervals from a bootstrap that resamples genes. Excluding the controls that were never built roughly halves the difference and leaves it significant. (c) The same comparison restricted to that subset, decomposed into direction, magnitude and the residual after simple sequence features; the direction-only vector reproduces the full difference, while the residual after simple sequence features does not exclude zero and drives both arms below chance; that row is shown de-emphasised and is not interpreted. (d) The difference by whether the real substitution moves the codon to a rarer or to a more common synonym; intervals that include zero are de-emphasised.

### What the dependence is on, and how far comparisons carry

The dependence is on the specific gene, not on its homologous relatives. Training that excluded not only the test genes but every gene sharing an HGNC family with them (Methods, *Family and support analyses*) removes about 44% of the remaining training genes, so we added a control that removes the same number of training genes chosen at random. The step from known genes to unseen genes costs every model between 0.11 and 0.16 in AUC. The further step to removing whole families costs nothing measurable: the unseen-to-family differences stay within ±*0.02* and do not agree in sign across models, and the comparison against the diversity control is likewise flat (Table S53; Figure 5a).

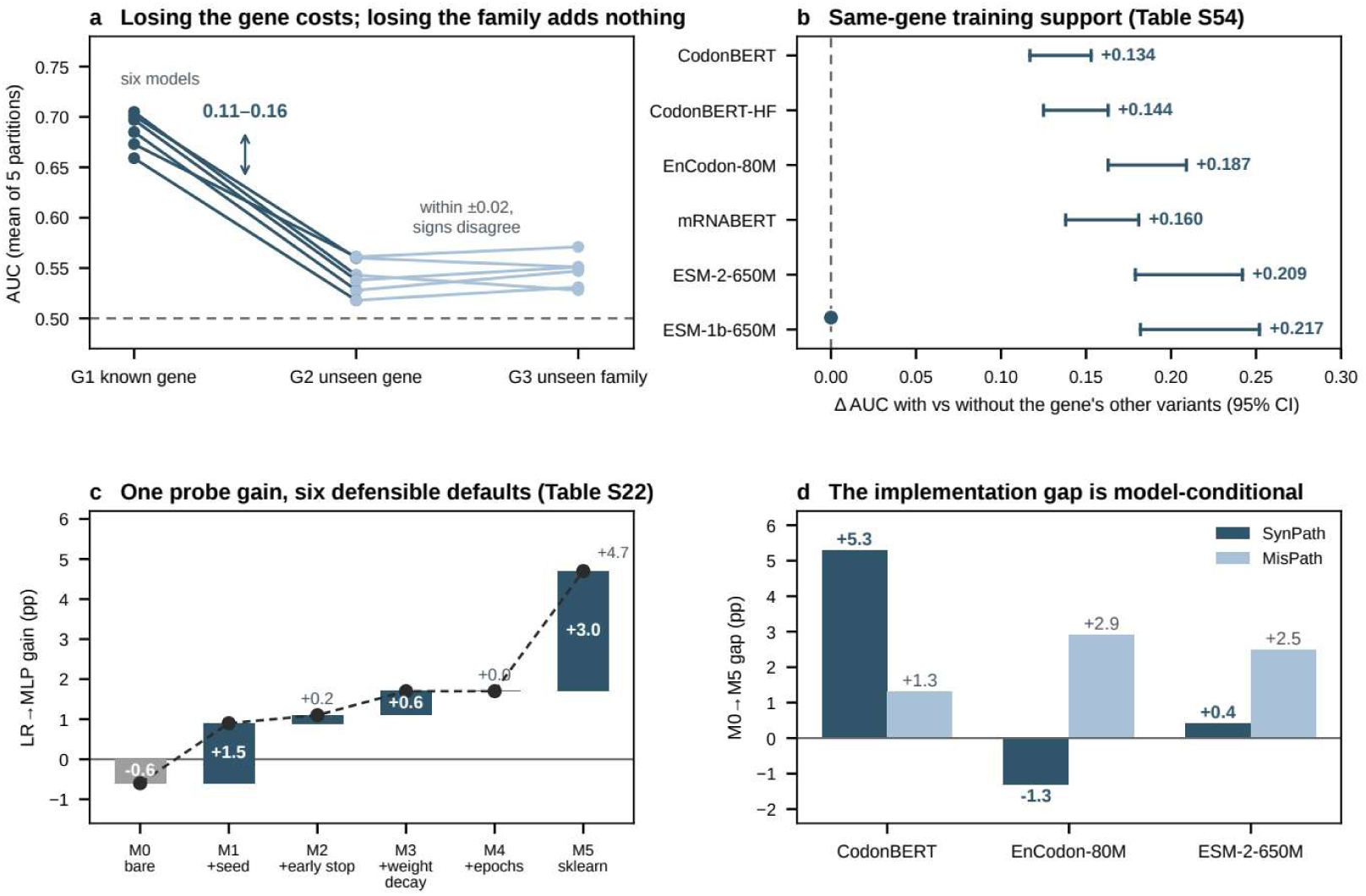
Attribution to the specific gene, and what belongs to the protocol. (a) Removing a test gene, then its whole homologous family. G1 uses a test gene whose other variants remain in training; G2 removes the test gene entirely; G3 additionally removes every training gene sharing an HGNC family with it, with training-set size matched across arms; points are means over five independent gene partitions (Table S53). (b) Training support from the other variants of a held-out gene: each marker is the change in AUC when the gene’s remaining variants are available against their removal, with a 95% interval from a gene-clustered bootstrap (Table S54). (c) Stepwise build-up of one model’s nonlinear-probe gain from a bare implementation to the reference library implementation (fixed seed, early stopping, weight decay, doubled epochs), in percentage points; bars give each step’s increment and the dashed trace the running total; no single step produces the apparent gain (Table S22). (d) The same step-wise gap across three models and the two task arms; it is model-conditional, and in one case negative (Table S23).

#### Is the same-gene dependence peculiar to codon models?

It is not. For each gene carrying more than one variant we held out one variant and trained twice, once with the gene’s other variants available and once without them, with training-set size matched in both arms (Methods, *Family and support analyses*). Both families lose accuracy when the gene’s other variants are removed, and the loss is significant for every model (Table S54). Removing a gene’s other variants also removes their gene-specific biology, so we do not read this as a clean measure of leakage; what the comparison supports is that both families draw on gene-linked information to a similar degree (Figure 5b).

#### How much of the difference is the analysis rather than the model?

Three components of the protocol move the answers (Methods, *Protocol sensitivity and software*): the pooling rule, whose apparent effect vanishes under a second implementation of the same classifier (Table S20); the epoch-selection rule, the larger of the two — chosen on the test split, the apparent gain of a nonlinear probe ranges up to twenty-three percentage points, and chosen on a held-out validation split, the gain turns negative on one of the two tasks for all three models compared (Tables S11 and S12); and feature scaling, which we treat as a requirement rather than a choice.

#### Does the pattern depend on curated labels?

On independently measured function, the positional shortcut does not survive gene hold-out. We repeated the comparison on saturation genome editing data, where function is measured rather than annotated: six genes, 51,315 variants in total, 26,783 of them coding single-nucleotide variants, of which 5,966 are synonymous (Table S32; Methods, *External data*) (28–33). Both arms use a positional feature, so what is tested is whether a position prior generalises to unseen genes when the outcome is measured rather than curated. It does not: on the synonymous arm, random splitting separates the classes while gene hold-out does not, and on the coding arm the held-out value falls slightly below chance (Figure 6). No frozen representation was evaluated on these data, and the check covers six genes, all of them DNA-damage or cancer genes; the below-chance value on the coding arm is reported without an explanation we can defend.

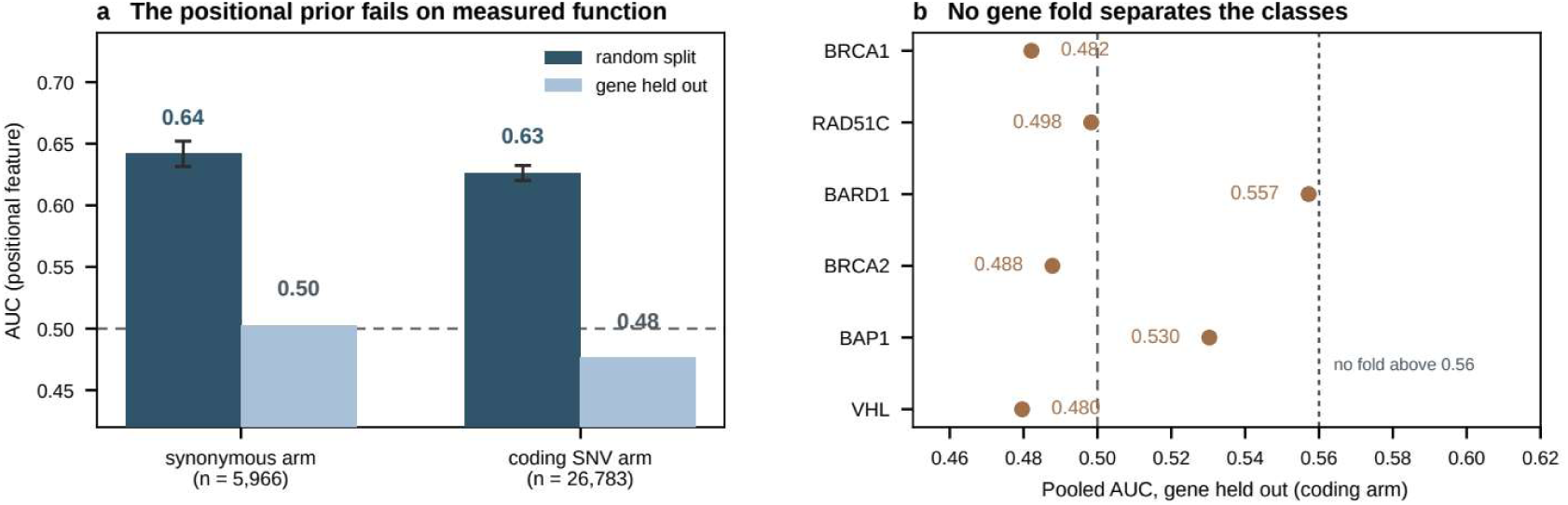
The same contraction on measured function. Saturation genome editing measurements from six genes (Table S32). (a) Pooled AUC on the synonymous and coding arms under random splitting and gene hold-out; error bars are the standard deviation over five random splits. (b) The held-out value for each gene of the coding arm; the dotted line marks the highest value reached by any gene fold, and the dashed line marks chance.

## Discussion

The results separate into a negative part and a positive part, and the positive part is the smaller of the two. Most of the predictability in this task is not about the variant: every representation we tested loses between 0.10 and 0.26 in AUC when whole genes are held out, and afterwards the entire panel sits in a narrow band around chance. For protein models this is not an empirical finding but a consequence of the input — the two alleles cannot differ, so whatever the model scores must come from background. Independent work on a different variant class reaches the same shape: under random splitting, a null model reading only per-gene pathogenicity rates approaches the performance of current missense predictors (34). The same failure to carry over appears for the positional shortcut where the outcome is measured rather than curated.

The dependence is also narrower than one might expect: it is the identity of the specific gene that carries the model, not its homology group — removing a test gene’s entire family costs nothing beyond removing the gene itself. Protein language models likewise carry a strong signal of the organism a sequence belongs to (35).

A fourth source emerged that we had not anticipated, and it is the strongest one in the study: exon position, which requires neither a model nor a sequence. This is a property of how the variants were collected rather than of the models — pathogenic synonymous variants are reported where clinicians look, and the ones that disrupt splicing lie at exon ends — so any benchmark assembled from ClinVar synonymous variants will inherit it. The remedy is not a stronger model but a control matched on position.

The positive part is that a component does survive both controls. Once the background is subtracted, what remains still predicts pathogenicity in genes the model has never seen, and it does so significantly better than a control variant at the same position of the same codon. It is smaller than the uncorrected comparison suggests, because a third of the controls were never built and contribute nothing; it is directional; and it is concentrated in substitutions that move a codon towards a rarer synonym. It is invisible in the absolute embeddings, which are dominated by gene identity.

Several limits bound these claims. The family-level comparison is unresolved rather than null, because the gene is the unit of replication and the intervals are wide. The real variants and their controls are not comparable on every simple feature, the largest gap being the proportion of transitions, 76% against 15% (Table S57), and a stricter, chemistry-matched control can be built for only 34.3% of the records; what that row establishes is that the difference is sensitive to how the control is built (Table S59). Several quantities reported here are small enough that the protocol matters, the pooling rule, the epoch-selection rule and the probe implementation each moving the answers by amounts comparable to the effects being measured. Fine-tuning leaves the models above the frozen band but is neither same-protocol nor full-parameter (36), so a frozen read-out is the conservative half of the story. The evaluated subset is not a random sample of SynPath: the leave-one-gene-out design covers only the 1,847 variants on transcripts large enough to form an informative fold.

## Conclusions

On this benchmark, performance that looks substantial under random splitting largely reflects gene identity and variant position rather than the substitution. Under gene hold-out the entire panel falls to a narrow band around chance, while a positional baseline that reads no sequence reaches a substantially higher level. A small variant-specific component does survive gene hold-out in the response vector of a coding-sequence model — smaller than a naive comparison reports, directional, and concentrated in substitutions that move a codon towards a rarer synonym. On independently measured function, only the positional arm was tested, and it fails to generalize. Benchmarking synonymous variants without holding genes out, and without a control matched on position, will overstate what these models have learned.

## List of abbreviations

AUC: area under the receiver operating characteristic curve
CDS: coding sequence
CI: confidence interval
cLM: codon language model
HGNC: HUGO Gene Nomenclature Committee
LLR: log-likelihood ratio
LoRA: low-rank adaptation
LR: logistic regression
LOGO: leave-one-gene-out
MDES: minimum detectable effect size
MLP: multilayer perceptron
OOF: out-of-fold
pLM: protein language model
SD: standard deviation
SGE: saturation genome editing
SNV: single-nucleotide variant.

## Methods

### Data

SynPath comprises 2,840 ClinVar synonymous variants across 497 transcripts, with labels as recorded by ClinVar; MisPath applies the same protocol to 5,000 ClinVar missense variants balanced 1:1, carried only as a contrast arm. The frozen input inventory and every exclusion are listed in Additional file 1 (Table S38).

### Sequence inputs

Each record is presented in two forms cut from the same stored transcript coding sequence: the codon input, the variant’s codon with the 15 codons on either side as separate tokens (31 tokens); and the character input, the raw substring spanning 45 nucleotides on either side, truncated at transcript ends (46–91 nucleotides). For a variant on the minus strand only the two alleles are complemented before the window is cut.

### Strand, frame, and exclusions

Stored windows lie on the coding strand and are used as given; the reading frame is taken from the variant’s own CDS position. The control comparisons cover 1,839 (position-matched) and 1,837 (protein-model) records.

### Splitting and probe

Two regimes are used throughout: a stratified five-fold random split with a fixed seed, and 240 fixed leave-one-gene-out folds, each holding out all variants of one transcript. A logistic regression (*C* = *1*, L-BFGS) is fitted to each training fold, with features standardised on the training folds only (12,37,38). One probe configuration is applied to all 24 frozen representations, without fine-tuning or per-model search, so differences between rows are attributable to the representations (Table S39). Throughout, the drop of a representation is its random-split AUC minus its gene-held-out AUC.

### Positional baselines

Distances to the nearest exon junction and to the nearest CDS terminus are taken from the frozen variant table; the score is the negated distance, and no model is fitted.

### Controls

The position-matched control substitutes a different synonymous codon at the variant’s own position, with the substituted base at the same within-codon position, constructed only where a third synonymous codon exists and drawn uniformly under a fixed seed. The upstream control substitutes an unrelated base 15 nt away; the permuted-label arm shuffles labels as a null reference. Records whose control vector is exactly zero keep it as zero (34.3% of the set), and every comparison reports which scope it used.

### Family and support analyses

Genes were annotated with curated HGNC family membership (39) (457 genes, 396 families). The family-removal and same-gene support analyses use gene-level partitions with training-set size matched across arms, and both were repeated under a second, independently drawn partition, retaining direction and interval (Table S46).

### Coordinate provenance

The set mixes two assembly builds (1,444 records on GRCh37, 1,396 on GRCh38); at 1,059 recorded positions the recorded reference base disagrees with the GRCh38 reference, all in the GRCh37 half. Model inputs are cut from stored coding sequences, so no representation-based result depends on the build (Table S4).

### External data

The independent check uses saturation genome editing measurements from six genes (RAD51C, BAP1, BARD1, BRCA2, BRCA1, VHL): 51,315 variants, 26,783 coding single-nucleotide variants, of which 5,966 are synonymous; the outcome is the assayed functional score rather than a curated classification.

### Statistical inference

The unit of replication is the gene, not the variant. Paired comparisons use a gene-resampling bootstrap (1,000 replicates), and a comparison is reported as resolved only when its interval excludes zero. A fold-level sensitivity analysis puts the resolution floor at about 9 percentage points of AUC, above the effects reported here; the surviving component is below that floor, so our reading of it rests on the gene-level bootstrap. The response-vector comparison against the position-matched control is the single pre-specified test; the subgroup analyses (Tables S56 to S58) are exploratory and are not corrected for multiplicity.

### Protocol sensitivity and software

Results are sensitive to the probe, to its implementation, and to epoch selection; Section S6 of Additional file 1 states what each table varies and which comparisons across them are licit. Models were run with PyTorch (40) and Hugging Face Transformers (41).

## Declarations

### Ethics approval and consent to participate

Not applicable. This study analyses publicly available ClinVar variant annotations and involves neither human participants nor identifiable data.

### Consent for publication

Not applicable.

### Availability of data and materials

The benchmark datasets, the evaluation pipeline and the analysis code that produces every number reported here are available at github.com/missdu/CodonBench. The release analysed here is archived as version 1.0.1, doi:10.5281/zenodo.23176516. Project name: CodonBench evaluation and audit pipeline. Project home page: github.com/missdu/CodonBench. Archived version: 1.0.1 (Zenodo), doi:10.5281/zenodo.23176516. Operating system: Linux (x86-64). Programming language: Python 3.11. Other requirements: NumPy, pandas, scikit-learn, PyTorch. License: MIT. Restrictions for non-academics: none.

Input variant annotations are derived from public ClinVar releases, whose accessions are recorded in the frozen variant table; derived tables, fold definitions and per-fold result files are provided as additional files. The saturation genome editing measurements used for the external validation are publicly available from their original publications and from MaveDB (33). Pretrained model weights are available from the Hugging Face repositories cited in the model panel. Every value reported in the main text is stated together with the result file and fold definition it was computed from.

An earlier version of this manuscript has been deposited as a preprint on bioRxiv. DOI: 10.64898/2026.08.12.744371 (10). The present manuscript supersedes it: here the positional source is measured as a separate quantity under a single fold definition, and a position-matched control is added to the response-vector comparison.

### Author contributions (CRediT)

*Yuanqing Liang:* Conceptualization, Methodology, Software, Formal analysis, Investigation, Visualization, Writing – original draft. *Weimin Zhu:* Conceptualization, Methodology, Software, Resources, Validation. *Xiaoyong Pan:* Supervision, Funding acquisition, Resources, Formal analysis, Writing – review & editing. *Huixian Li:* Supervision, Investigation, Formal analysis, Resources, Writing – review & editing. All authors read and approved the final manuscript.. All authors read and approved the final manuscript.

## Acknowledgements.

We thank the Pan Lab at Shanghai Jiao Tong University for computational resources and support.

## Use of artificial intelligence

During software development for this study the authors used CodeArts, an AI-assisted coding tool, to draft and debug parts of the evaluation code and to suggest revisions to the wording of the manuscript. The tool generated no data, ran no analysis and derived no result: every experiment was executed by the authors against the frozen input release, and every number, table and figure reported here was produced by the authors’ own pipeline and checked by them independently. All AI-suggested code and text were read line by line, revised where necessary, and approved by the authors, who take full responsibility for the accuracy and integrity of the work. The tool meets none of the criteria for authorship and is not listed as an author.

## Additional files

### Additional file 1

*File name:* Supplementary Information. *File format:* PDF. *Title of data:* Supplementary tables S1–S51 and S53–S59 (with S57b and S59b; 60 tables in total) and Supplementary Figures S1–S3 for “CodonBench: separating gene context, exon position, and variant-specific component in synonymous-variant classification”. *Description of data:* the frozen input inventory and audit summary; the splice-adjacency and coordinate-provenance tables; the full numeric results of every experiment reported in the main text, given per fold and per model; the control-construction counts behind the response-vector comparison; and three supporting figures (review status by ClinVar review level, training curves, and layer-wise probing).

## References

1. Carlos Outeiral, Charlotte M. Deane. Codon language embeddings provide strong signals for use in protein engineering. Nature Machine Intelligence. 2024. doi:10.1038/s42256-024-00791-0

2. Mohsen Naghipourfar, Siyu Chen, Mary K. Howard, Christian B. Macdonald, Ali Saberi, Timothy Hagen, Mohammad R. K. Mofrad, Willow Coyote-Maestas, Hani Goodarzi. A suite of foundation models captures the contextual interplay between codons. bioRxiv. 2024. doi:10.1101/2024.10.10.617568

3. Alexander Rives, Joshua Meier, Tom Sercu, Siddharth Goyal, Zeming Lin, others. Biological structure and function emerge from scaling unsupervised learning to 250 million protein sequences. Proceedings of the National Academy of Sciences. 2021;118(15):e2016239118. doi:10.1073/pnas.2016239118

4. Hugo Dalla-Torre, others. Nucleotide transformer: Building and evaluating robust foundation models for human genomics. Nature Methods. 2025. doi:10.1038/s41592-024-02523-z

5. Sizhen Li, Saeed Moayedpour, Ruben Li, Michael Bailey, Saleh Riahi, others. CodonBERT large language model for mRNA vaccines. Genome Research. 2024;34(7):1027–35. doi:10.1101/gr.278870.123

6. Yi Xiong, An Wang, Yu Kang, Cheng Shen, Cheng-Yu Hsieh, Tingjun Hou. mRNABERT: Advancing mRNA sequence design with a universal language model and comprehensive dataset. Nature Communications. 2025. doi:10.1038/s41467-025-65340-8

7. Adibvafa Fallahpour, others. CodonTransformer: A multispecies codon optimizer using context-aware neural networks. Nature Communications. 2025. doi:10.1038/s41467-025-58588-7

8. Logan Hallee, Nikolaos Rafailidis, Jason P. Gleghorn. cdsBERT — extending protein language models with codon awareness. bioRxiv. 2023. doi:10.1101/2023.09.15.558027

9. Shushan Toneyan, Katharina Scholz, Carlo De Donno, Felix Noack, Sven Auslaender, Tony Cijsouw, Joshua L. Payne. Assessing codon language models for context-aware codon optimization in nucleic acid-based medicines. bioRxiv. 2026. doi:10.64898/2026.08.11.744178

10. Yuanqing Liang, Weimin Zhu, Huiying Liang, Xiaoyong Pan. A leakage-controlled benchmark shows apparent codon-language-model advantages in synonymous-variant prediction are evaluation artifacts. bioRxiv. 2026. doi:10.64898/2026.08.12.744371

11. Kevin Musgrave, Serge Belongie, Ser-Nam Lim. A metric learning reality check. In: Proceedings of ECCV 2020. 2020. doi:10.1007/978-3-030-58595-2_41

12. Tiago Pimentel, Naomi Saphra, Adina Williams, Ryan Cotterell. Pareto probing: Trading off accuracy for complexity. In: Proceedings of EMNLP 2020. 2020. doi:10.18653/v1/2020.emnlp-main.254

13. Alon Jacovi, Avi Caciularu, Omer Goldman, Yoav Goldberg. Stop uploading test data in plain text: Practical strategies for mitigating data contamination by evaluation benchmarks. In: Proceedings of EMNLP 2023. 2023. doi:10.18653/v1/2023.emnlp-main.308

14. Sayash Kapoor, Arvind Narayanan. Leakage and the reproducibility crisis in machine-learning-based science. Patterns. 2023;4(9):100804. doi:10.1016/j.patter.2023.100804

15. Junlan Feng, others. Benchmarking DNA foundation models for genomic and genetic tasks. Nature Communications. 2025. doi:10.1038/s41467-025-65823-8

16. Na You, Chao Liu, Hui Lin, Shiwei Wu, Guanjun Chen, Ning Shen. Benchmarking pre-trained genomic language models for RNA sequence-related predictive applications. Nature Communications. 2025. doi:10.1038/s41467-025-66899-y

17. Wenjun Cheng, Zhihang Song, Yang Zhang, Shenglong Wang, Di Wang, Meng Yang, Lei Li, Jianzhu Ma. DNALONGBENCH: A benchmark suite for long-range DNA prediction tasks. Nature Communications. 2025. doi:10.1038/s41467-025-65077-4

18. Zishuo Zeng, Ariel A. Aptekmann, Yana Bromberg. Decoding the effects of synonymous variants. Nucleic Acids Research. 2021;49(22):12673–91. doi:10.1093/nar/gkab1159

19. Luca Cartegni, Shern L. Chew, Adrian R. Krainer. Listening to silence and understanding nonsense: Exonic mutations that affect splicing. Nature Reviews Genetics. 2002;3(4):285–98. doi:10.1038/nrg775

20. Minh-Tam Pham, Michael J. G. Milevskiy, Jane E. Visvader, Yunshun Chen. Incorporating exon-exon junction reads enhances differential splicing detection. BMC Bioinformatics. 2025. doi:10.1186/s12859-025-06210-4

21. Franco Pagani, Marina Raponi, Francisco E. Baralle. Synonymous mutations in CFTR exon 12 affect splicing and are not neutral in evolution. Proceedings of the National Academy of Sciences. 2005. doi:10.1073/pnas.0502288102

22. J. V. Chamary, Joanna L. Parmley, Laurence D. Hurst. Evidence for purifying selection against synonymous mutations in mammalian exonic splicing enhancers. Molecular Biology and Evolution. 2006. doi:10.1093/molbev/msj035

23. Karin B. Nielsen, Sofie Sørensen, Luca Cartegni, Thomas J. Corydon, others. Seemingly neutral polymorphic variants may confer immunity to splicing-inactivating mutations: A synonymous SNP in exon 5 of MCAD protects from deleterious mutations in a flanking exonic splicing enhancer. American Journal of Human Genetics. 2007. doi:10.1086/511992

24. Jonathan Frazer, Pascal Notin, Mafalda Dias, others. Disease variant prediction with deep generative models of evolutionary data. Nature. 2021;599:91–5. doi:10.1038/s41586-021-04043-8

25. Gonzalo Benegas, Sanjit Singh Batra, Yun S. Song. DNA language models are powerful predictors of genome-wide variant effects. Proceedings of the National Academy of Sciences. 2023;120. doi:10.1073/pnas.2311219120

26. Brian M. Schilder, Zhuoyi Liu, Jenna Desmarais, others. Genetic background shapes AI-predicted variant effects. bioRxiv. 2026. doi:10.64898/2026.04.04.715328

27. Žiga Avsec, Natasha Latysheva, Jun Cheng, others. Advancing regulatory variant effect prediction with AlphaGenome. Nature. 2026;649:1206–18. doi:10.1038/s41586-025-10014-0

28. Gregory M. Findlay, Riza M. Daza, Brotman Martin, Melissa D. Zhang, Anh P. Leith, Jay Shendure. Accurate classification of BRCA1 variants with saturation genome editing. Nature. 2018. doi:10.1038/s41586-018-0461-z

29. Rebeca Olvera-León, Fang Zhang, Victoria Offord, Yuhan Zhao, Huei San Tan, others. High-resolution functional mapping of RAD51C by saturation genome editing. Cell. 2024. doi:10.1016/j.cell.2024.08.039

30. Andrew J. Waters, Tobias Brendler-Spaeth, Danielle Smith, Victoria Offord, Huei San Tan, others. Saturation genome editing of BAP1 functionally classifies somatic and germline variants. Nature Genetics. 2024. doi:10.1038/s41588-024-01799-3

31. Max Buckley, Céline Terwagne, Andrea Ganner, Laura Cubitt, Rebecca Brewer, others. Saturation genome editing maps the functional spectrum of pathogenic VHL alleles. Nature Genetics. 2024. doi:10.1038/s41588-024-01800-z

32. Hongyuan Huang, Chunling Hu, Jie Na, Steven N. Hart, Rohan D. Gnanaolivu, others. Functional evaluation and clinical classification of BRCA2 variants. Nature. 2025. doi:10.1038/s41586-024-08388-8

33. Alan F. Rubin, Jeremy Stone, Alice H. Bianchi, Benjamin J. Capodanno, others. MaveDB 2024: A curated community database with over seven million variant effects from multiplexed functional assays. Genome Biology. 2025. doi:10.1186/s13059-025-03476-y

34. S. Harrizi, I. Nait Irahal, K. Mostafa, D. Arnoult. Gene identity, not variant effect, dominates ClinVar benchmarks of missense pathogenicity predictors. Preprint; 2026. doi:10.64898/2026.08.21.746180

35. Logan Hallee, Tomer Peleg, Nikolaos Rafailidis, Jason P. Gleghorn. Protein language models are accidental taxonomists. BMC Bioinformatics. 2026. doi:10.1186/s12859-026-06491-3

36. Edward J. Hu, Yelong Shen, Phillip Wallis, Zeyuan Allen-Zhu, Yuanzhi Li, Shean Wang, Lu Wang, Weizhu Chen. LoRA: Low-rank adaptation of large language models. In: International conference on learning representations (ICLR 2022). OpenReview.net; 2022.

37. Guillaume Alain, Yoshua Bengio. Understanding intermediate layers using linear classifier probes. In: International conference on learning representations (ICLR 2017). OpenReview.net; 2017.

38. John Hewitt, Percy Liang. Designing and interpreting probes with control tasks. In: Proceedings of EMNLP-IJCNLP 2019. 2019. doi:10.18653/v1/d19-1275

39. Ruth L. Seal, Bryony Braschi, Kristian Gray, Jonathan McClay, Susan Tweedie, Elspeth A. Bruford. Genenames.org: The HGNC and PGNC resources in 2026. Nucleic Acids Research. 2026. doi:10.1093/nar/gkaf1229

40. Adam Paszke, Sam Gross, Francisco Massa, Adam Lerer, others. PyTorch: An imperative style, high-performance deep learning library. In: Advances in neural information processing systems 32. 2019.

41. Thomas Wolf, Lysandre Debut, Victor Sanh, Julieni Chaumond, others. Transformers: State-of-the-art natural language processing. In: Proceedings of EMNLP 2020: System demonstrations. 2020. doi:10.18653/v1/2020.emnlp-demos.6

