## Supporting Information: supplementary tables S1-S59 and figures S1-S3 for "CodonBench: separating gene context, exon position, and variant-specific component in synonymous-variant classification"

File format: PDF

#### Contents

Supplementary Tables S1-S51 and S53-S59, including S57b and S59b

(60 tables in total), and Supplementary Figures S1-S3.

for:

**CodonBench: separating gene context, exon position, and variant-specific component in synonymous-variant classification**

Table S52 was retired while the working documents were merged;

its content now appears in Table S42.

Supplementary Information for
"CodonBench: separating gene context, exon position, and variant-specific component in synonymous-variant classification"

#### Note on numbering and how to find things

This supplement was assembled by merging several working documents, so the tables and figures below do not appear in numerical order. The numbers are kept as stable identifiers so that every cross-reference in the main text stays valid; those references were checked to resolve, not merely assumed to. Table S52 was retired while the working documents were merged; its content lives in Table S42. We ask that these numbers be kept unchanged in production rather than renumbered. The index below gives the order in which each item appears.


- Supplementary Table S38: Frozen input inventory and audit summary. (p. 4)
- Supplementary Table S4: Splice-adjacent variants, and which boundary is measured (p. 5)
- Supplementary Table S30: Experiment-to-model-subset mapping (p. 6)
- Supplementary Table S1: Model details (p. 8)
- Supplementary Table S35: Source provenance—original tasks, evaluation protocols, and claims (p. 9)
- Supplementary Table S13: Five-dimensional evaluation framework (p. 11)
- Supplementary Table S39: Representation panel, width, family, and offline loadability. (p. 12)
- Supplementary Table S2: P1/P2 validation (LR, random splits) (p. 14)
- Supplementary Table S7: Depth signature (6 pretrained, 5-fold CV) (p. 15)
- Supplementary Table S8: LOGO-CV results (6 pretrained models) (p. 16)
- Supplementary Table S17: Channel specificity (13 neural, single 80/20 split) (p. 18)
- Supplementary Table S18: Channel specificity per-model LR→MLP gains (p. 19)
- Supplementary Table S34: Full 21-model panel (single 80/20 split, unified protocol) (p. 20)
- Supplementary Table S37: Reference numbers at a glance—random-split versus gene-held-out evaluation (p. 22)
- Supplementary Table S3: Zero-shot LLR AUC (Level 0) (p. 23)
- Supplementary Table S5: LoRA fine-tuning (random splits) (p. 24)
- Supplementary Table S6: Tokenization ablation (LOGO-CV, SynPath) (p. 25)

- Supplementary Table S9: LoRA gene-stratified collapse (p. 26)
- Supplementary Table S10: Synonym randomization (p. 27)
- Supplementary Table S14: LoRA rank ablation (p. 28)
- Supplementary Table S15: Tokenization ablation (random splits, SynPath) (p. 29)
- Supplementary Table S16: Tokenization LOGO-CV paired $t$ tests (p. 30)
- Supplementary Table S20: Pooling ablation (CodonBERT, SynPath LOGO-CV) (p. 31)
- Supplementary Table S22: M0→M5 waterfall (CodonBERT, SynPath LOGO-CV) (p. 32)
- Supplementary Table S23: Cross-model M0→M5 decomposition (p. 33)
- Supplementary Table S24: Tokenization ablation under a single 80/20 split (p. 34)
- Supplementary Table S25: Paired sign test (tokenization LOGO-CV) (p. 35)
- Supplementary Table S26: 20-seed PyTorch MLP sweep (p. 36)
- Supplementary Table S27: 20-seed sklearn MLP sweep (p. 37)
- Supplementary Table S40: Three positional quantities on the same variants. (p. 38)
- Supplementary Table S41: Positional baselines under the leave-one-gene-out folds ($n$ = 1,847). (p. 38)
- Supplementary Table S45: The positional score applied to the consequence classes of a second ClinVar collection. (p. 38)
- Supplementary Table S42: Response vector against two controls, in two character-level coding-sequence models (leave-one-gene-out). (p. 40)
- Supplementary Table S43: Control construction coverage and exclusions. (p. 41)
- Supplementary Table S11: Regression best-epoch selection (p. 42)
- Supplementary Table S12: Regression validation-based early stopping (p. 43)
- Supplementary Table S19: Layer-wise probing (CodonBERT-HF, 12 layers) (p. 44)
- Supplementary Table S28: Sensitivity at 90% specificity (p. 45)
- Supplementary Table S36: LoRA-level regression protocol-matched comparison. (p. 46)
- Supplementary Table S31: MDES analysis (p. 47)
- Supplementary Table S29: Attribute regression ($R^2$) (p. 48)
- Supplementary Table S21: Codon-optimality independence (p. 49)
- Supplementary Table S44: Uncertainty types, what addresses each, and what is not addressed. (p. 50)
- Supplementary Table S46: Partition robustness for the family and support analyses. (p. 51)
- Supplementary Table S32: Position-only baseline two-arm experiment (p. 52)
- Supplementary Figure S1: Review-status stratification (p. 8)

- Supplementary Note S1: Pooling and standardization corrections (p. 54)
- Supplementary Table S33: Reproduction of the original protocol, and collapse under gene-held-out evaluation (p. 55)
- Supplementary Figure S2: Training curves (p. 14)
- Supplementary Figure S3: Layer-wise probing (p. 23)
- Supplementary Table S47: Frozen representations under the two splitting regimes (p. 65)
- Supplementary Table S48: Pre-modelling checks on the two model families (p. 65)
- Supplementary Table S49: Paired codon-model minus protein-model differences under gene hold-out (p. 66)
- Supplementary Table S50: Position of each variant relative to an exon junction (p. 66)
- Supplementary Table S51: Positional baselines under the leave-one-gene-out folds (p. 66)
- Supplementary Table S53: Removing whole homologous families, with a size-matched control (p. 67)
- Supplementary Table S54: Training support from other variants of the held-out gene (p. 67)
- Supplementary Table S55: Decomposition of the variant-specific component (p. 68)
- Supplementary Table S58: Increment of the response vector over stated baselines (p. 68)
- Supplementary Table S56: The component by stratum (p. 69)
- Supplementary Table S57: Coverage, comparability and checks (p. 70)
- Supplementary Table S57b: Rank decomposition of the response vector (cdsBERT, scope A) (p. 70)
- Supplementary Table S59: The response vector against a control matched on nucleotide chemistry (p. 70)
- Supplementary Table S59b: Whether each control matches the real arm on substitution chemistry (p. 72)

### S1. Data and audit

*Contents.* Data inventory, frozen-package fields, sequence-input construction, strand and phase verification, missing and mismatched items, splice-site confounds, experiment-to-model-subset mapping. Sequence hashes are not tabulated here: the input arrays are distributed with the analysis archive, and their identifiers are the file names recorded in Table S38 rather than digests reproduced in the supplement.

**[S1] Data inventory and audit.**

*Purpose.* Record what the frozen input package contains, what was verified before modelling, and which records fall outside the evaluated set, so that every downstream count can be traced to a field.

*Key observations.* The frozen package holds 2,840 ClinVar synonymous variants from 497 transcripts with a balanced label split. Strand and frame handling resolve 2,823 records (99.4%), and of those 2,821 translate to the same amino acid under both alleles, an agreement that is checked rather than assumed. The evaluated subset is smaller than the resolvable set and is not a random draw from it: the leave-one-gene-out design covers 1,847 records with a pathogenic fraction of 54.1%, against 42.3% in the 993 records that sit on transcripts too small to form a fold. Exon-junction and CDS-terminus distances are missing for 26 records each, and those records are excluded from the positional analyses alone. Genomic coordinates span two assembly builds.

*Relation to main text.* Results, the opening paragraphs of Results; Methods (data, strand and frame, exclusions).

**Table S38: Frozen input inventory and audit summary.**

| Item | Value | Note |
|---|---|---|
| Variants in frozen package | 2,840 | ClinVar synonymous, balanced 1:1 |
| Transcripts | 497 | as recorded in `transcript_id` |
| Pathogenic / benign | 1,420 / 1,420 | exact balance |
| Resolved after strand and frame | 2,823 (99.4%) | codon parse succeeds |
| of which translate identically | 2,821 | check passed, not a construction step |
| Leave-one-gene-out folds | 240 | one transcript held out per fold |
| Variants inside those folds | 1,847 | evaluated set |
| pathogenic fraction | 54.1% | 1,000 pathogenic, 847 benign |
| Variants outside the folds | 993 | transcripts too small for a fold |
| pathogenic fraction | 42.3% | differs from the evaluated set |
| Median fold size | 5 variants | fold-level statistics are unstable |
| Missing exon-junction distance | 26 | excluded from positional analyses only |
| Missing CDS-terminus distance | 26 | same records |
| Assembly GRCh37 / GRCh38 | 1,444 / 1,396 | matched build-wise, never across |
| Frozen-package fields | 25 columns | see list below |

**Frozen-package fields.** `allele_id`, `variation_id`, `genome_build`, `chromosome`, `position_vcf`, `ref`, `alt`, `gene_id`, `gene_id_raw`, `transcript_id_with_version`, `consequence`, `reference_codon`, `alternate_codon`, `reference_aa`, `alternate_aa`, `cds_position`, `exon_boundary_distance`, `cds_boundary_distance`, `n_exon_in_tx`, `clinvar_label`, `review_status`, `clinvar_release`, `strand`, `codon_parse_ok`, `seq_len`.

*Two corrections applied to the recorded recipe.* The stored windows are already on the coding strand, so reverse-complementing the window is an error; for minus-strand records only the two alleles are complemented. The reading frame is taken from the CDS position as $(p-1) \bmod 3$ and is never selected by requiring the two alleles to translate identically, since a frame chosen that way would guarantee the agreement it is later used to check.

*How the two sequence inputs are produced.* Each record is presented to a model in two forms, both cut from the same stored transcript CDS at the variant's own position, and both produced by the same script from the same per-transcript CDS record. The **codon input** takes the variant's codon plus the 15 codons on either side and joins them as separate codon tokens; because the window is defined in codons it is always 31 tokens long. The **character input** takes the raw CDS substring spanning 45 nt on either side of the variant nucleotide and is truncated at the transcript ends, so its length runs from 46 to 91 nt and is not fixed. The two forms therefore share a centre but not an extent, 93 nt against 46–91 nt. Neither is reverse-complemented, for the reason given above: the CDS is stored in coding orientation, so a substring of it is already on the coding strand and already in frame. The two arms of the tokenization comparison consequently differ only in how one sequence is segmented, never in which sequence is read. A table of per-record window hashes is not reproduced here, for the reason given under *Items not verified here.*

*Items not verified here.* Sequence hashes for the stored windows are recorded in the run manifest but are not reproduced in this table; they should be added from the manifest before submission if the journal requires per-record checksums.

#### Supplementary Table S4: Splice-adjacent variants, and which boundary is measured

*Purpose.* Records what is known about each variant's proximity to a splice junction, and separates that quantity from proximity to a CDS terminus, which is a different boundary.
*Key observations.* This table previously reported that only 0.007% of synonymous variants here exceed the SpliceAI DS $\geq 0.2$ threshold, and the main text concluded from it that the positional shortcut is not a restatement of predicted splice disruption. Both are withdrawn. That figure combined a numerator drawn from the records that the annotator managed to match, 211 in total, with the denominator of a different population, every synonymous variant in ClinVar, so it is not the quantity it was presented as. A second run built the same way matched 162 records and found every one of them benign, so even the coverage is not reproducible. What stands independently of all this is that the distance to the nearest *exon junction* is strongly associated with the label and works as a classifier on its own (Table S41), because pathogenic synonymous variants accumulate where splice signals are.

An earlier version of this table reported that only 0.2% of variants fall within 3 nt of a *CDS boundary*, and read that as evidence that splice-site confounding could be set aside. That count concerns the two termini of the coding sequence, not exon junctions, and a coding sequence spans several coding exons. On the same variants the CDS-terminus distance reaches an AUC of 0.534 whereas the exon-junction distance reaches 0.843 (Table S41). The two quantities are therefore not interchangeable, and the earlier count does not bear on the splicing question.
*Coordinate provenance.* Each record carries the assembly build it was annotated on. Of the 2,840 variants, 1,444 are recorded against GRCh37 and 1,396 against GRCh38, and taking each record's position and reading the GRCh38 reference base there gives a disagreement with the recorded reference allele for 1,059 records, 37.3% of the set. The predictor was run against GRCh38 alone, so those records were unreachable, and every GRCh38-annotated record agrees, which is what identifies the build mismatch as the cause. Coordinates play no part in constructing model inputs, which are cut from stored coding sequences, so nothing else in this paper depends on which build a record carries. Lifting the GRCh37 subset over and repeating the annotation is the obvious repair and is left undone: the evaluation server has no network access and no runtime able to run the predictor, the two being confirmed by direct test.
*Relation to main text.* Results, *The strongest single source is position, and it reads no sequence*; Methods, *Coordinate provenance*; *Positional baselines.*

| **Metric** | **Value** |
|---|---|
| Records whose recorded reference base matches the GRCh38 reference | 1,781 of 2,840 (62.7%) |
| Records where the two disagree, so the record cannot be located | 1,059 of 2,840 (37.3%) |
| Records annotated on GRCh37 coordinates | 1,444 |
| Records annotated on GRCh38 coordinates | 1,396 |
| GRCh37 records whose base matches GRCh38 | 385 of 1,444 (26.7%) |
| GRCh37 records that disagree | 1,059 of 1,444 (73.3%) |
| AUC, distance to nearest exon junction | 0.843 |
| AUC, distance to nearest CDS terminus | 0.534 |

#### Supplementary Table S30: Experiment-to-model-subset mapping

*Purpose.* Maps every experiment in the main text to the specific subset of the 21-model panel used, so that reviewers can trace which models contribute to each figure panel and each numerical claim. Section numbers refer to the main text (Results Sec. 2.1–2.4; M = Methods). The 21-model panel comprises 15 neural models (9 codon cLMs, 2 character CDS LMs, 2 protein LMs, 2 DNA LMs) and 6 non-neural baselines. "Evaluable neural" = 13 (9 cLM + 2 char + 2 pLM); the 2 DNA LMs are probed in the full panel (Table S34) but excluded from the channel-specificity contrast (Table S17). Counts are nested, not additive: 15 = 13 + 2 DNA LMs, 13 = 11 cLM + 2 protein LMs, 11 = 9 codon-level + 2 character-level. From-scratch models: v1 (4.8K CDS, 10 ep, 20M), v3a (4.8K CDS, 100 ep, 20M), v3b (114K CDS, 30 ep, 20M), v4 (114K CDS, 30 ep, 110M), v2 (synthetic-CDS negative control, 20M). The 6 pretrained models referenced throughout are: CodonBERT, CodonBERT-HF, EnCodon-80M, mRNABERT (4 cLMs) + ESM-2-650M, ESM-1b-650M (2 pLMs).
*Key observations.* Different experiments use different model subsets, determined by experimental design: the tokenization ablation requires matched from-scratch pairs; the depth signature uses 6 pretrained models; channel specificity extends to all 13 evaluable neural models; LoRA uses 5 cLMs with compatible architectures; the M0→M5 decomposition uses 3 models representing distinct behaviors.
*Relation to main text.* Methods, *Representation panel* (second paragraph); provides the complete mapping for all figure panels and numerical claims.

| Sec. | Fig. | Experiment | Model subset | $n$ |
|---|---|---|---|---|
| *Results Sec. 2.1 — Random-split protocol* | | | | |
| 2.1 | 2a | P1/P2 direction reversal | full 21-model panel: 15 neural + 6 non-neural baselines | 21 |
| 2.1 | 2b | Tokenization ablation | 5 from-scratch pairs ×{codon, char}: v1, v3a, v3b, v4, v2 | 10 |
| 2.1 | 2c | Depth signature | 6 pretrained: 4 cLMs (CodonBERT, CB-HF, EnCodon-80M, mRNABERT) + 2 pLMs | 6 |
| 2.1 | 2d | Channel specificity | 13 evaluable neural (9 cLM + 2 char + 2 pLM) | 13 |
| 2.1 | 2e | LoRA ceiling | 5 codon cLMs with LoRA-compatible transformers (Supp. Table S9) | 5 |
| *Results Sec. 2.2 — Gene-held-out collapse* | | | | |
| 2.2 | 3a | Tokenization collapse | Same 5 from-scratch pairs (LOGO-CV) | 10 |
| 2.2 | 3b | Probe-impl. shift | Same 6 pretrained (LOGO-CV; sklearn vs PyTorch MLP) | 6 |
| 2.2 | 3c | LoRA collapse | Same 5 cLMs (gene-stratified split) | 5 |
| 2.2 | 3d | Gene-overlap quantification | Dataset-level (no models) | — |
| 2.2 | 3e | Channel-specific collapse | 6 pretrained + onehot_pos (LOGO-CV) | 7 |
| *Results Sec. 2.3 — Probe-implementation artifact* | | | | |
| 2.3 | 4a | M0→M5 waterfall | CodonBERT (SynPath, LOGO-CV) | 1 |
| 2.3 | 4b | Seed sweep (20 seeds) | CodonBERT (SynPath, LOGO-CV) | 1 |
| 2.3 | 4c | Pooling gate | CodonBERT: CLS vs mean (SynPath, LOGO-CV) | 1 |
| 2.3 | 4d | Cross-model M0→M5 | 3 models: CodonBERT, EnCodon-80M, ESM-2-650M ×{SynPath, MisPath} | 3 |

| Sec. | Fig. | Experiment | Model subset | $n$ |
|---|---|---|---|---|
| 2.3 | 4e | Regression best-epoch bias | 3 cLMs: CodonBERT, CB-HF, EnCodon-80M $\times$ {mRFPExpr, FungalExpr} | 3 |
| 2.3 | 4f | $R^2 \approx 0$ tasks | Same 3 cLMs $\times$ {EcoliExpr, mRNAStab} | 3 |
| *Results Sec. 2.4 — Synonym randomization* | | | | |
| 2.4 | 5a | Syn-rand: face value | 3 cLMs (CodonBERT, CB-HF, EnCodon-80M) + ESM-2 + char control | 5 |
| 2.4 | 5b | Syn-rand: collapse | Same 5 models (5-fold pooled + LOGO-CV pooled + PyTorch MLP) | 5 |
| 2.4 | 5c | Numerical paradox | CodonBERT: LR vs MLP, per-fold vs pooled | 1 |
| 2.4 | 5d | Scale comparison | v3b (20M), v4 (110M), CodonBERT (ref; single-split) | 3 |
| *Methods — Additional experiments* | | | | |
| M | — | Attribute regression | 3 cLMs: CodonBERT, CB-HF, EnCodon-80M (GC3, CAI, position, pathogenicity) | 3 |
| M | — | Layer-wise probing | CodonBERT-HF (12 transformer layers; SynPath + MisPath) | 1 |
| M | — | Zero-shot LLR | 3 cLMs with MLM heads: CodonBERT, CB-HF, EnCodon-80M | 3 |

#### S2. Model registry

*Contents.* Per-model checkpoint identifier, parameter count, config, source provenance, evaluation framework, and offline loadability. Tokenizer identities are reported as the tokenization strategy actually used at extraction together with the unknown-token count for the models that could be loaded; for the representations taken from the cache, tokenizer digests are not reported, and the registry says so rather than leaving the field blank.

#### Supplementary Table S1: Model details

*Purpose.* Lists all 21 evaluation models (15 neural + 6 non-neural baselines) with their type, parameter count, vocabulary size, pretraining corpus, and reference.
*Key observations.* The panel spans 9 codon-level cLMs, 2 character-level CDS LMs, 2 protein LMs, 2 DNA LMs, and 6 baselines (including the onehot_pos memorization probe). Of the 24 surveyed codon/CDS-level models, 12 were loadable through standard interfaces and 11 were fully evaluable at all probe levels; Mistral-Codon-1M was excluded (CUDA OOM at MLP). The remaining 7 of the 31 surveyed models are non-CDS (2 protein-level, 2 DNA-level, 3 other) and do not enter this evaluable count.
*Relation to main text.* Methods, *Representation panel*; the model-selection pipeline (Surveyed → Evaluable → Panel) is described there.

| **Model** | **Type** | **Params** | **Vocab** | **Corpus** | **Reference** |
|---|---|---|---|---|---|
| CodonBERT | codon cLM | 10M | 69 | 10M CDS | [1] |
| CodonBERT-HF | codon cLM | 10M | 69 | 10M CDS | HF reupload |
| EnCodon-80M | codon cLM | 80M | 69 | 4M CDS | [2] |
| EnCodon-620M | codon cLM | 620M | 69 | 4M CDS | [2] |
| CodonTransformer | codon cLM | — | 69 | — | [3] |
| CaLM | codon cLM | — | 69 | — | [4] |
| mRNABERT | codon cLM | — | 69 | — | [5] |
| Mistral-117M | codon cLM | 117M | 69 | — | — |
| Mistral-16M | codon cLM | 16M | 69 | — | — |
| cdsBERT | char CDS LM | — | 9 | — | [6] |
| cdsBERT-plus | char CDS LM | — | 9 | — | — |
| ESM-2-650M | protein LM | 650M | 33 | 250M proteins | [7] |
| ESM-1b-650M | protein LM | 650M | 33 | 250M proteins | [8] |
| NT-v2-500M | DNA LM | 500M | 12 | 1000G | [9] |
| NT-v2-50M | DNA LM | 50M | 12 | 1000G | [9] |
| onehot_pos | baseline | — | — | — | position+codon |
| one-hot | baseline | — | — | — | — |
| k-mer | baseline | — | — | — | — |
| codon-usage | baseline | — | — | — | — |
| GC content | baseline | — | — | — | — |
| AlphaMissense | baseline | — | — | — | [10] |

**Notes.** 21 models = 15 neural + 6 baselines. 9 codon cLMs + 2 char CDS LMs + 2 protein LMs + 2 DNA LMs = 15 neural. **Counts in this section are nested, not additive.** "Evaluable neural" = 13 = 11 cLMs (9 codon-level + 2 character-level) + 2 protein LMs. The 2 DNA LMs are probed and reported in the full panel (Table S34) but are excluded from the 13-model channel-specificity contrast (Table S17), which compares codon-level with protein-level channels only. Throughout, "cLM" denotes the 11 codon/CDS-level language models; any figure or table using a different subset names it (Table S30).

#### Supplementary Table S35: Source provenance—original tasks, evaluation protocols, and claims

*Purpose.* For each of the 21 panel models, records the task on which the model's developers reported their results, the train/test split protocol they used, whether that split controls for gene homology, the claim made in the source paper, and whether the source paper reports a number on a synonymous-variant task. This table establishes that the 21 models were validated on heterogeneous tasks under heterogeneous protocols, and that only two (CodonBERT and ESM-2-650M, the latter a protein model) have a published value on the synonymous-variant task used in this study.

*Key observations.* (1) The source tasks span codon optimization, mRNA design, EC classification, protein engineering, protein structure, and genome annotation—there is no common benchmark. (2) Of the seven archived external model papers, only three used homology- or chromosome-aware splits (CaLM, ESM-1b, NT-v2); all codon-level cLMs with archived papers (CodonBERT, mRNABERT, cdsBERT) used random splits. (3) Only CodonBERT (0.734/0.849) and ESM-2-650M (0.685, a protein model) have directly reportable values on the synonymous-variant task; 19 of 21 models have no such value. The absence of a common benchmark is itself a finding, not a gap.

*Relation to main text.* Background, "Why a common task set, and why these tasks"; Discussion, "Why a shared task set is not a substitution of tasks" and "Cross-task generalization." Supports the claim that apparent performance is task-specific evaluation products rather than transferable representation capability, and establishes that no shared benchmark existed for the panel.

*Source.* Values are transcribed verbatim from archived copies of the source papers held in the project's literature folder. Models whose source PDFs were not archived (EnCodon, ESM-2, AlphaMissense) are marked —; no value was inferred from related models.

| Model | Source task | Split (source) | Controls homology? | Claim (verbatim) | Syn-variant value? |
|---|---|---|---|---|---|
| CodonBERT | mRNA property prediction (expression, degradation) + sequence classification | Random 0.7/0.15/0.15 | No | "CodonBERT outperforms previous mRNA prediction methods" | Yes (0.734/0.849) |
| CodonBERT-HF | Same as CodonBERT | Same | No | Same as CodonBERT | Same |
| EnCodon-80M | — | — | — | — | No |
| EnCodon-620M | — | — | — | — | No |
| CodonTransformer | Multi-species codon optimization (sequence generation) | No train/test split; 52-protein benchmark | N/A (generative) | "generates host-specific DNA sequences with natural-like codon distribution profiles" | No |
| CaLM | Protein engineering (species ID, abundance, melting point, solubility) | Identity-clustered holdout (40% AA identity) | **Yes** | "LLMs trained on codons ... outperform comparable state-of-the-art models across a variety of tasks" | No |
| mRNABERT | Full-length mRNA design (UTR, CDS, RBP sites, property prediction) | Random stratified + independent holdout | No | "state-of-the-art performance, outperforming previous models in the majority of tasks" | No |
| Mistral-117M | Trained in-house | N/A | N/A | N/A | No |
| Mistral-16M | Trained in-house | N/A | N/A | N/A | No |

| Model | Source task | Split (source) | Controls homology? | Claim (verbatim) | Syn-variant value? |
|---|---|---|---|---|---|
| cdsBERT | EC classification (vocab extended from AA to codons) | Random 10-fold CV ($n = 2{,}629$) | No | "codon usage bias is remarkably predictive and can even change the final structure of a protein" | No |
| cdsBERT-plus | Trained in-house (knowledge distillation on cdsBERT) | N/A | N/A | N/A | No |
| ESM-2-650M | — | — | — | — | Yes (0.685; protein model) |
| ESM-1b-650M | Protein structure, remote homology, variant effect | Structure/family holdout (SCOPe 3-level) | **Yes** | "information emerges in the learned representations on fundamental properties of proteins" | No |
| NT-v2-500M | Human genomics downstream prediction (18 tasks) | Chromosome holdout (chr 20/21) | **Yes** | "context-specific representations ... accurate predictions even in low-data settings" | No |
| NT-v2-50M | Same as NT-v2-500M | Same | **Yes** | Same | No |
| onehot_pos | Self-built baseline | N/A | N/A | N/A | No |
| one-hot | Self-built baseline | N/A | N/A | N/A | No |
| k-mer | Self-built baseline | N/A | N/A | N/A | No |
| codon-usage | Self-built baseline | N/A | N/A | N/A | No |
| GC content | Self-built baseline | N/A | N/A | N/A | No |
| AlphaMissense | — | — | — | — | No |

*Notes.* (1) Reported values are on each model's own task and are **not comparable across rows**. (2) This table reports **source** values only; our own measurements are in Tables S2, S17, and S34 and are not numerically interchangeable with this table. (3) Only **two** of the 21 models have a published value on the synonymous-variant task (CodonBERT, and ESM-2-650M which is a protein model). The absence of a common benchmark is itself a finding, not a gap. (4) Split type is reported per model. Random splits do not control for gene identity; three of the seven archived model papers use homology- or chromosome-aware splits. (5) Parameter and corpus corrections to Table S1: CodonBERT params 10M→∼110M; CodonTransformer params/corpus →89.6M / 1M DNA-protein pairs (164 organisms); CaLM params/corpus →86M / 9M cDNA (ENA); mRNABERT corpus →>18M mRNA; cdsBERT params →∼420M (ProtBERT-based); NT-v2 corpus 1000G→850 genomes (multispecies).

#### Supplementary Table S13: Five-dimensional evaluation framework

*Purpose.* Summarizes the five evaluation dimensions—two leakage controls (data split, epoch selection) and three diagnostic axes (probe depth, pooling, probe implementation)—that systematically vary the evaluation protocol.
*Key observations.* Disabling each leakage control shifts results in a fixed direction (+14.3 pp for data split; for the regression gain, the epoch rule isolated in a protocol-matched re-evaluation moves the gain by +1.7 to +4.7 pp, the remainder of the +6.4–23.7 pp change between the two regression protocols coming from the Ridge penalty and the split). The three diagnostic axes reveal how a given claim depends on evaluation choices: probe depth (+11.7–19.3 pp depth signature), pooling (+4.1 pp), and probe implementation (+4.7 pp).
*Relation to main text.* Methods, *Probe*; the framework emerged iteratively, not a priori (Methods).

| **Dim.** | **Name** | **Values** | **Effect of disabling** | **Type** |
|---|---|---|---|---|
| 1 | Data split | random → gene-held-out | inflates +14.3 pp | leakage control |
| 2 | Epoch selection | best-epoch → val-based | +1.7–4.7 pp (isolated) | leakage control |
| 3 | Probe depth | L0 → L1 → L2 → L3 | reveals depth signature | diagnostic |
| 4 | Pooling | CLS / mean | +4.1 pp difference (sklearn) | diagnostic |
| 5 | Probe impl. | sklearn / PyTorch | +4.7 pp difference | diagnostic |

**[S2] Representation registry and loadability.**
*Purpose.* Record which representations were evaluated, at what width, and which of them can be reproduced from cached weights on an offline node, so that the panel in the main text maps onto concrete artefacts.
*Key observations.* Twenty-eight embedding files were found with the required row count, and content fingerprinting shows that four pairs are byte-identical duplicates of the same embedding stored under two names. Removing those duplicates leaves 24 distinct representations spanning widths of 256 to 2048. They fall into four groups: codon language models, protein language models, DNA language models, and two models trained from scratch on this task. Only three models could be loaded from the offline cache on the compute node; three others failed under every loading route attempted and are listed as unavailable rather than silently omitted. Representations already extracted are unaffected by that limitation.

For the three models that could be loaded, extraction recorded the tokenization strategy actually applied and the number of tokens outside the vocabulary. All three used the same strategy on this input and produced no unknown tokens, so the panel contains no row whose score rests on a substituted or dropped base. For the twenty-one representations taken from the cache, no tokenizer digest is reported; those rows were extracted before this audit and their tokenization is not recoverable from the stored arrays.
*Relation to main text.* Results, *Gene hold-out collapses the whole panel to a band around chance*; Methods (representations).
**Table S39: Representation panel, width, family, and offline loadability.**
*Items not recorded here.* Tokenizer hashes and unknown-token rates for each model are not reproduced in this table. They are recoverable from the extraction logs but were not tabulated in this round, and are therefore marked as outstanding rather than filled with estimates. A minimal-inference check per model is likewise outstanding, with the exception of the three models loaded this round, each of which completed a smoke run on 120 variants before the full pass.

| Representation | Family | Width | Load | Note |
|---|---|---|---|---|
| CodonTransformer | codon LM | 768 | n/a (cached) | |
| CodonBERT | codon LM | 768 | n/a (cached) | stored under two file names |
| CodonBERT-HF | codon LM | 768 | n/a (cached) | stored under two file names |
| CaLM | codon LM | 768 | n/a (cached) | |
| EnCodon-620M | codon LM | 2048 | n/a (cached) | stored under two file names |
| EnCodon-620M (layer 1) | codon LM | 2048 | n/a (cached) | |
| EnCodon-80M | codon LM | 1024 | n/a (cached) | stored under two file names |
| mRNABERT | codon LM | 768 | n/a (cached) | |
| cdsBERT | codon LM | 1024 | loaded | re-extracted this round |
| cdsBERT-plus | char CDS LM | 1024 | loaded | re-extracted this round |
| Mistral-Codon-117M | codon LM | 768 | n/a (cached) | |
| Mistral-Codon-16M | codon LM | 256 | n/a (cached) | narrowest in panel |
| CodonBERT (mean pool) | codon LM | 768 | n/a (cached) | pooling variant |
| ESM-1b-650M | protein LM | 1280 | loaded | negative control |
| ESM-1b-650M (mean pool) | protein LM | 1280 | n/a (cached) | pooling variant |
| ESM-2-650M | protein LM | 1280 | failed | all loading routes failed |
| ESM-2-650M (mean pool) | protein LM | 1280 | n/a (cached) | pooling variant |
| ESM-2 (protein windows) | protein LM | 1280 | n/a (cached) | translated input |
| Nucleotide Transformer v2 500M | DNA LM | 1024 | failed | all loading routes failed |
| Nucleotide Transformer v2 50M | DNA LM | 512 | failed | all loading routes failed |
| from-scratch v3b | trained here | 512 | n/a (cached) | |
| from-scratch v3b (ctx 128) | trained here | 512 | n/a (cached) | |
| from-scratch v4 | trained here | 768 | n/a (cached) | |
| from-scratch v4 (ctx 128) | trained here | 768 | n/a (cached) | |
| Distinct representations | | 24 | | after removing 4 duplicate pairs |
| Width range | | 256–2048 | | |

### S3. Main results, per item

*Contents.* Per-model, per-fold, per-input-configuration complete results.

#### Supplementary Table S2: P1/P2 validation (LR, random splits)

*Purpose.* Validates the two falsifiable predictions from the information-theoretic decomposition: P1 (codon models win on SynPath) and P2 (protein models win on MisPath), under linear probing with random splits.
*Key observations.* On SynPath, EnCodon-620M achieves the highest LR AUC (0.797), exceeding ESM-1b (0.705, +9.2 pp; DeLong $p < 0.0001$) and ESM-2 (0.685, +11.2 pp); the CodonBERT reference reaches 0.734, exceeding ESM-1b (+2.9 pp, $p = 0.017$) and ESM-2 (+4.9 pp). On MisPath the direction reverses: ESM-2 (0.718) exceeds both EnCodon-620M (0.625) and CodonBERT (0.660). The onehot_pos memorization baseline scores 0.884 on SynPath—exceeding every neural model—foreshadowing the leakage analysis. AlphaMissense confirms missense-specificity (0.956 on MisPath, 0.440 on SynPath). Sample sizes: SynPath $n$ = 2,840 (1,420 pathogenic, 1,420 benign); MisPath $n$ = 5,000 (2,500/2,500).
*Relation to main text.* Results, *Gene hold-out collapses the whole panel to a band around chance*; the direction reversal implicates channel-specific access, not generic capacity.

| Model | SynPath AUC | MisPath AUC | Δ vs ESM-2 (Syn) | Δ vs ESM-2 (Mis) |
|---|---|---|---|---|
| CodonBERT | 0.734 | 0.660 | +4.9 | −5.8 |
| CodonBERT-HF | 0.705 | 0.659 | +2.0 | −5.9 |
| EnCodon-80M | 0.686 | 0.633 | +0.1 | −8.5 |
| EnCodon-620M | 0.797 | 0.625 | +11.2 | −9.3 |
| ESM-1b-650M | 0.705 | 0.708 | +2.0 | −1.0 |
| ESM-2-650M | 0.685 | 0.718 | 0 | 0 |
| onehot_pos | 0.884 | 0.755 | +19.9 | +3.7 |
| AlphaMissense | 0.440 | 0.956 | −24.5 | +23.8 |

*Note.* EnCodon-620M was re-extracted under the canonical CodonBench extractor (attention-masked mean at hidden_states[−2]) for this table, so its values are directly comparable to the other entries; it is the top-scoring codon cLM on SynPath. Table S2 is the protocol-controlled comparison: 5-fold stratified CV (`random_state` = 42), no feature standardization, no class weighting. Tables S17 and S34 apply the same protocol on the single 80/20 split that defines, and Table S7 reports a different pipeline (model-specific pooling with standardization, for cross-pipeline comparison only). None of these are numerically interchangeable, since the resampling scheme alone moves individual AUCs by several points. AlphaMissense is the one cell whose very existence is split-dependent: the SynPath value shown here (0.440) uses all 11 scored synonymous variants, distributed across the five folds, whereas none of them falls in the single-split test set, so Table S34 reports that cell as NA.

### Supplementary Table S7: Depth signature (6 pretrained, 5-fold CV)

*Purpose.* Quantifies the LR→MLP gain (depth signature) for six pretrained models under 5-fold CV on random splits, using the unified pipeline (model-specific pooling: CLS for codon models, mean for protein models; standardization applied).
*Key observations.* All six models gain +11.7 to +19.3 pp from LR to MLP on SynPath, generalizing across architectures. The onehot_pos position-only baseline also exceeds every neural MLP (0.931 vs 0.86–0.89), so the gain occurs entirely below a memorization ceiling—hinting that the depth signature reflects more effective use of gene identity rather than extraction of synonymous signal.
*Relation to main text.* Results, *Gene hold-out collapses the whole panel to a band around chance*; the depth signature generalizes beyond scratch models to pretrained models, but the position-only ceiling foreshadows the leakage analysis.

| **Model** | **LR AUC** | **MLP AUC** | **LR→MLP gain** | **Type** |
|---|---|---|---|---|
| CodonBERT | 0.725 | 0.882 | +15.7 | cLM |
| CodonBERT-HF | 0.694 | 0.888 | +19.3 | cLM |
| EnCodon-80M | 0.698 | 0.889 | +19.1 | cLM |
| mRNABERT | 0.695 | 0.861 | +16.6 | cLM |
| ESM-2-650M | 0.743 | 0.867 | +12.4 | pLM |
| ESM-1b-650M | 0.749 | 0.866 | +11.7 | pLM |

#### Supplementary Table S8: LOGO-CV results (6 pretrained models)

*Purpose.* Gene-held-out (LOGO-CV) results on SynPath for six pretrained models and the onehot_pos position-only baseline, under two MLP implementations that share the same embeddings and the same standardization. Central to the second collapse (Results, *Gene hold-out collapses the whole panel to a band around chance*, run-in *Can the collapse be blamed on the encoding or the readout?*).
*Key observations.* Panel A: Under the scikit-learn `MLPClassifier`, LOGO-CV gains range −3.7 to +4.7 pp on SynPath—far smaller than the +11.7–19.3 pp under random splits. ESM-2 collapses to near chance (0.507). The onehot_pos position-only baseline retains 0.666 on SynPath (LR AUC 0.884 → 0.666, −21.7 pp) and 0.740 on MisPath (0.755 → 0.740, −1.5 pp; MisPath value from a separate run of the same protocol), so the position-only baseline mirrors the channel asymmetry rather than removing it. The channel-specific collapse (; 12.7–23.6 pp on SynPath vs 5.3–7.8 pp on MisPath) is the per-model drop in LR AUC from 5-fold random-split evaluation (Supplementary Table S7) to the unified-pipeline LOGO-CV values in Panel B. Panel B: Under the PyTorch MLP, CodonBERT's gain vanishes to −0.0 pp. Because the two panels share the same embeddings and the same standardization, their LR column is identical; only the MLP implementation differs, so the largest single-model discrepancy (CodonBERT, 4.7 pp) is probe-implementation dependent, not representation dependent. Panel C: Per-fold-mean and pooled aggregation give opposite signs for CodonBERT (+4.7 pp vs −1.1 pp), a seventh degree of freedom.
*Relation to main text.* Results, *Gene hold-out collapses the whole panel to a band around chance*, run-in *Can the collapse be blamed on the encoding or the readout?*; the sklearn vs PyTorch discrepancy motivates the M0→M5 ablation (Results, *The dependence is on the specific gene, and part of every comparison is the protocol*).

**Panel A: SynPath LOGO-CV, scikit-learn `MLPClassifier`** (same embeddings as Panel B: model-specific pooling, CLS for codon models and mean for protein models; `StandardScaler`; per-fold-mean AUC)

| Model | LR AUC | MLP AUC | Gain (pp) |
|---|---|---|---|
| CodonBERT | 0.564 | 0.611 | +4.7 |
| CodonBERT-HF | 0.537 | 0.542 | +0.5 |
| EnCodon-80M | 0.537 | 0.544 | +0.7 |
| mRNABERT | 0.568 | 0.546 | −2.2 |
| ESM-2-650M | 0.507 | 0.528 | +2.1 |
| ESM-1b-650M | 0.527 | 0.490 | −3.7 |
| onehot_pos | 0.666 | 0.740 | +7.4 |

**Panel B: SynPath LOGO-CV, PyTorch MLP** (unified pipeline; same embeddings as Panel A: model-specific pooling, CLS for codon models and mean for protein models; `StandardScaler`; per-fold-mean AUC)

| Model | LR AUC | MLP AUC | Gain (pp) |
|---|---|---|---|
| CodonBERT | 0.564 | 0.564 | −0.0 |
| CodonBERT-HF | 0.537 | 0.540 | +0.3 |
| EnCodon-80M | 0.537 | 0.568 | +3.1 |
| mRNABERT | 0.568 | 0.538 | −3.0 |
| ESM-2-650M | 0.507 | 0.503 | −0.4 |
| ESM-1b-650M | 0.527 | 0.489 | −3.9 |

**Panel C: Per-fold vs pooled (CodonBERT)**

| **Aggregation** | **LR→MLP gain (pp)** | **Note** |
|---|---|---|
| Per-fold mean | +4.7 | sklearn MLP |
| Pooled | −1.1 | sklearn MLP |

*Note.* Panels A and B were produced with the `lbfgs` solver (`max_iter` = 1000, $C = 1.0$) on `StandardScaler`-transformed features; both panels share that configuration and the same embeddings, which is why their LR columns agree digit for digit (only the MLP implementation differs). The no-standardization Level-1 baseline described in Methods uses the same `lbfgs` solver with `max_iter` = 2000 and no class weighting, so the two differ only in feature standardization and iteration budget. A few legacy probe scripts use the `liblinear` solver instead of `lbfgs`; in every instance we compared, the two differ by at most one unit in the third decimal of AUC. Panel C aggregates the same 240 valid LOGO-CV folds as Panels A and B (497 transcripts total; folds with fewer than two samples per class excluded).

#### Supplementary Table S17: Channel specificity (13 neural, single 80/20 split)

*Purpose.* LR and MLP AUCs for all 13 evaluable neural models on both tasks under the same unified single 80/20 split that defines and the full panel of Table S34, extending the depth-signature analysis beyond the six pretrained models of Table S7. Protocol: logistic regression (L2, $C = 1.0$, lbfgs, max_iter = 2000, no feature standardization, no class weighting) or an MLP (standardized inputs, hidden $(128, 64)$, early stopping), on the canonical caches pooled by attention-masked mean at `hidden_states`$[-2]$. Table S2 reports the same models under the protocol-controlled 5-fold analysis; the two are not numerically interchangeable and no value is carried between them.
*Key observations.* Under the single split the per-model SynPath LR AUCs span 0.576–0.760, a much wider range than under 5-fold, which is why states only within-split comparisons. Every model gains from LR to MLP on SynPath (+7.1 to +25.9 pp, median +16.4); on MisPath the same probe changes little and not uniformly (-3.9 to +6.4 pp, median +1.7; 5 of 13 fall). The SynPath–MisPath asymmetry is what plots.
*Relation to main text.* Results, *Gene hold-out collapses the whole panel to a band around chance*; the channel specificity of the depth signature motivates the leakage analysis.

| Model | Syn LR | Syn MLP | Mis LR | Mis MLP |
|---|---|---|---|---|
| CodonBERT | 0.716 | 0.820 | 0.661 | 0.681 |
| CodonBERT-HF | 0.667 | 0.812 | 0.646 | 0.670 |
| EnCodon-80M | 0.687 | 0.862 | 0.636 | 0.700 |
| EnCodon-620M | 0.760 | 0.831 | 0.621 | 0.658 |
| CodonTransformer | 0.685 | 0.815 | 0.686 | 0.673 |
| CaLM | 0.692 | 0.824 | 0.654 | 0.671 |
| mRNABERT | 0.608 | 0.800 | 0.687 | 0.657 |
| Mistral-117M | 0.631 | 0.821 | 0.628 | 0.633 |
| Mistral-16M | 0.576 | 0.818 | 0.661 | 0.679 |
| cdsBERT | 0.592 | 0.824 | 0.629 | 0.621 |
| cdsBERT-plus | 0.603 | 0.862 | 0.627 | 0.658 |
| ESM-2-650M | 0.652 | 0.809 | 0.702 | 0.686 |
| ESM-1b-650M | 0.677 | 0.841 | 0.719 | 0.680 |

*Note.* All four columns come from a single extraction-and-probing run (unified protocol), so the LR→MLP deltas are internally comparable; the absolute AUCs are not comparable to Table S2 (5-fold) or Table S7 (different pooling and standardization). The machine-readable per-model rows, including the embedding path of every cell, are in the benchmark release.

### Supplementary Table S18: Channel specificity per-model LR→MLP gains

*Purpose.* Per-model LR→MLP gains on SynPath and MisPath, computed from the AUCs of Table S17 (same split, same protocol); this is the table behind the panel of.
*Key observations.* On SynPath every model gains (+7.1 to +25.9 pp, median +16.4), regardless of family — the effect is not confined to one architecture. On MisPath gains are small and mixed (-3.9 to +6.4 pp, median +1.7; 5 of 13 fall), so the extra flexibility is specific to the synonymous channel rather than a general consequence of a richer probe.
*Relation to main text.* Results, *Gene hold-out collapses the whole panel to a band around chance*; the SynPath–MisPath asymmetry in LR→MLP gains is a key observation motivating the gene-identity leakage hypothesis.

| **Model** | **Syn gain (pp)** | **Mis gain (pp)** | **Type** |
|---|---|---|---|
| CodonBERT | +10.3 | +1.9 | cLM |
| CodonBERT-HF | +14.4 | +2.4 | cLM |
| EnCodon-80M | +17.5 | +6.4 | cLM |
| EnCodon-620M | +7.1 | +3.7 | cLM |
| CodonTransformer | +13.0 | -1.3 | cLM |
| CaLM | +13.2 | +1.7 | cLM |
| mRNABERT | +19.1 | -3.0 | cLM |
| Mistral-117M | +18.9 | +0.5 | cLM |
| Mistral-16M | +24.2 | +1.7 | cLM |
| cdsBERT | +23.2 | -0.8 | char |
| cdsBERT-plus | +25.9 | +3.1 | char |
| ESM-2-650M | +15.7 | -1.6 | pLM |
| ESM-1b-650M | +16.4 | -3.9 | pLM |

#### Supplementary Table S34: Full 21-model panel (single 80/20 split, unified protocol)

*Purpose.* Reports the linear-probe AUC (Level 1) for all 21 panel models on both tasks, under the single 80/20 split that defines — the complete set plotted in that panel. Every row uses one protocol (logistic regression, $C = 1.0$, lbfgs, max_iter = 2000, no feature standardization, no class weighting; canonical caches pooled by attention-masked mean at `hidden_states`$[-2]$). Table S2 is the protocol-controlled 5-fold comparison; Table S17 gives the per-model LR→MLP gains.

*Key observations.* SynPath AUC spans 0.505–0.884 across the panel. The codon cLMs sit at both ends of the neural range: 6 of the nine exceed ESM-2 (0.652), 3 fall below it (Mistral-117M 0.631, mRNABERT 0.608, Mistral-16M 0.576) and none sits level; so under random splits P1 is a top-of-panel separation rather than a uniform family separation. On MisPath the ordering is cleaner: both protein LMs (0.702, 0.719) exceed every codon cLM (maximum 0.687), so P2 holds model by model. The onehot_pos position-only baseline (0.884) exceeds every neural model on SynPath.

*Relation to main text.* Results, *Gene hold-out collapses the whole panel to a band around chance*; the full panel behind the P1/P2 crossover.

| Model | Type | SynPath AUC | MisPath AUC |
|---|---|---|---|
| CodonBERT | codon cLM | 0.716 | 0.661 |
| CodonBERT-HF | codon cLM | 0.667 | 0.646 |
| EnCodon-80M | codon cLM | 0.687 | 0.636 |
| EnCodon-620M | codon cLM | 0.760 | 0.621 |
| CodonTransformer | codon cLM | 0.685 | 0.686 |
| CaLM | codon cLM | 0.692 | 0.654 |
| mRNABERT | codon cLM | 0.608 | 0.687 |
| Mistral-117M | codon cLM | 0.631 | 0.628 |
| Mistral-16M | codon cLM | 0.576 | 0.661 |
| cdsBERT | char CDS LM | 0.592 | 0.629 |
| cdsBERT-plus | char CDS LM | 0.603 | 0.627 |
| ESM-2-650M | protein LM | 0.652 | 0.702 |
| ESM-1b-650M | protein LM | 0.677 | 0.719 |
| NT-v2-500M | DNA LM | 0.703 | 0.610 |
| NT-v2-50M | DNA LM | 0.603 | 0.560 |
| onehot_pos | baseline | 0.884 | 0.758 |
| one-hot | baseline | 0.562 | 0.659 |
| k-mer ($k = 6$) | baseline | 0.730 | 0.646 |
| codon-usage (GC3) | baseline | 0.506 | 0.485 |
| GC content | baseline | 0.505 | 0.547 |
| AlphaMissense | baseline | NA | 0.937 |

*Note.* All 21 rows share one protocol and one split, so no per-row ”source” column is needed; the embedding path of every cell is recorded in the benchmark release. AlphaMissense is reported as NA on SynPath because only 11 synonymous variants in the whole cohort carry an AlphaMissense score and none of them falls in the 568-row test set, so no AUC can be formed on this split; the 5-fold column of Table S2 contains the corresponding value (0.440) and the two are not interchangeable. Tables S2, S17 and S34 report overlapping models under the same probe but different resampling, and are not numerically interchangeable.

*Notes on the non-neural baselines.* The $k$-mer baseline is reported at $k = 6$. Because the sweep contains more than one value, all three are given here, on the same split as the panel, so that the choice is auditable rather than selective: SynPath/MisPath AUC is 0.567/0.615 at $k = 3$, 0.595/0.626 at $k = 4$ and 0.730/0.646 at $k = 6$. Performance rises monotonically with $k$ on both tasks, and $k = 6$ is also the token length used by DNA language models, so it is the principled choice as well as the strongest one. The codon-usage baseline uses GC3, the fraction of G/C at the third codon position, and excludes codon length, which would otherwise reintroduce position information. Both this baseline and GC content score near chance on both tasks (SynPath 0.506 and 0.505; MisPath 0.485 and 0.547), which is the intended control: the cLM signal is not attributable to codon composition or to simple sequence statistics.

#### Supplementary Table S37: Reference numbers at a glance—random-split versus gene-held-out evaluation

*Purpose.* Collects, in one place, the numbers that CodonBench exists to supply: the performance of a fixed panel of models under the conventional random split and under gene-held-out cross-validation, with everything else held constant (same embeddings, same model-specific pooling, same `StandardScaler`, same probe code). It is the quick-reference counterpart to the per-question tables, and it is the table a reader should consult before comparing any cLM number across papers.

*Key observations.* Under random splits every model gains from LR to MLP on SynPath (+11.7 to +19.3 pp). Under gene-held-out evaluation the same six models sit within ±3.9 pp of a flat LR→MLP line, and their LR AUC falls −12.7 to −23.6 pp. On MisPath the LR collapse is far smaller, which is the channel asymmetry the main text reports. No model's gene-held-out LR AUC exceeds the position-only ceiling (see Supplementary Tables S8 and S33).

*Relation to main text.* Results, *Gene hold-out collapses the whole panel to a band around chance* and § 2.2 (gene-held-out collapse); supersedes no prior table, and no number in it is new—every value is a restatement, in one place, of a number already reported in Supplementary Tables S7 and S8.

*Source.* `results/unified_eval/eval_results/pretrained_unified_results.json`, generated from a single evaluation pass per model: `lr_5fold_mean` and `mlp_5fold_mean` for the random split, `lr_logo_mean` and `mlp_logo_mean` for gene-held-out CV (240 valid folds for SynPath). The MLP column is the PyTorch implementation (Supplementary Table S8, Panel B); under the scikit-learn `MLPClassifier` the same LOGO-CV run gives CodonBERT 0.611 instead of 0.564 (Panel A), a probe-implementation difference the main text discusses.

| Model | Type | Random LR | Random MLP | LOGO-CV LR | LOGO-CV MLP | ΔLR (pp) | ΔMLP (pp) |
|---|---|---|---|---|---|---|---|
| **Panel A: SynPath** | | | | | | | |
| CodonBERT | cLM | 0.725 | 0.882 | 0.564 | 0.564 | -16.1 | -31.8 |
| CodonBERT-HF | cLM | 0.694 | 0.888 | 0.537 | 0.540 | -15.8 | -34.8 |
| EnCodon-80M | cLM | 0.698 | 0.889 | 0.537 | 0.568 | -16.1 | -32.1 |
| mRNABERT | cLM | 0.695 | 0.861 | 0.568 | 0.538 | -12.7 | -32.3 |
| ESM-2-650M | pLM | 0.743 | 0.867 | 0.507 | 0.503 | -23.6 | -36.3 |
| ESM-1b-650M | pLM | 0.749 | 0.866 | 0.527 | 0.489 | -22.2 | -37.8 |
| **Panel B: MisPath** | | | | | | | |
| CodonBERT | cLM | 0.651 | 0.668 | 0.590 | 0.566 | -6.1 | -10.3 |
| CodonBERT-HF | cLM | 0.648 | 0.671 | 0.592 | 0.590 | -5.6 | -8.1 |
| EnCodon-80M | cLM | 0.635 | 0.675 | 0.560 | 0.582 | -7.5 | -9.4 |
| mRNABERT | cLM | 0.667 | 0.661 | 0.589 | 0.594 | -7.8 | -6.7 |
| ESM-2-650M | pLM | 0.649 | 0.693 | 0.596 | 0.632 | -5.3 | -6.1 |
| ESM-1b-650M | pLM | 0.642 | 0.688 | 0.573 | 0.601 | -6.9 | -8.7 |

*Notes.* (1) ΔLR and ΔMLP are gene-held-out minus random-split, in percentage points; a negative value means the random split was higher. (2) Random-split columns are 5-fold CV means; gene-held-out columns are 240-fold LOGO-CV means (per-fold mean, the aggregation reported throughout). (3) The `onehot_pos` baseline—the position-only baseline that sets the ceiling any claimed synonymous-channel signal must clear—is reported in Supplementary Tables S33 (random split) and S8 (LOGO-CV); it is not repeated here because its gene-held-out evaluation uses a different feature pipeline. (4) Values are rounded to three decimals; no value in this table is new, and none should be compared against numbers obtained under a different split or pooling scheme.

#### Supplementary Table S3: Zero-shot LLR AUC (Level 0)

*Purpose.* Zero-shot log-likelihood ratio evaluation (Level 0): tests whether the synonymous channel is directly accessible from the MLM pretraining objective without any explicit probe.
*Key observations.* All three cLMs with accessible MLM heads score near chance on SynPath (0.484–0.526), confirming that the synonymous channel is invisible to the pretraining objective. MisPath LLR AUCs are modestly above chance (0.563–0.581).
*Relation to main text.* Results, *Gene hold-out collapses the whole panel to a band around chance*; motivates probing at Levels 1–3.

| Model | SynPath LLR AUC | MisPath LLR AUC |
|---|---|---|
| CodonBERT | 0.526 | 0.581 |
| CodonBERT-HF | 0.484 | 0.563 |
| EnCodon-80M | 0.512 | 0.572 |

### S4. Ablations and permutations

*Contents.* All ablation arms and negative controls.

#### Supplementary Table S5: LoRA fine-tuning (random splits)

*Purpose.* LoRA fine-tuning results (Level 3, random splits) for five codon cLMs with LoRA-compatible transformer architectures.
*Key observations.* SynPath AUCs reach 0.874–0.919, but none exceed the onehot_pos baseline's MLP ceiling (0.931). That maximal fine-tuning freedom cannot beat pure memorization suggests LoRA, too, is largely fitting gene identity.
*Relation to main text.* Results, *Gene hold-out collapses the whole panel to a band around chance*; the LoRA ceiling is compared to onehot_pos to show leakage extends to the top of the probing ladder.

| **Model** | **SynPath AUC** | **MisPath AUC** |
|---|---|---|
| CodonBERT | 0.909 | 0.809 |
| CodonBERT-HF | 0.903 | 0.811 |
| EnCodon-80M | 0.874 | 0.701 |
| EnCodon-620M | 0.897 | 0.754 |
| CodonTransformer | 0.919 | 0.836 |

#### Supplementary Table S6: Tokenization ablation (LOGO-CV, SynPath)

*Purpose.* LOGO-CV results for the from-scratch tokenization ablation on SynPath: tests whether the codon–char gap survives gene-held-out evaluation.
*Key observations.* Under LOGO-CV no real-CDS condition retains a codon advantage: the MLP gap is +2.0, +0.6, −0.5 and +0.5 pp for v1/v3a/v3b/v4 (10-seed means), against +16.6, +16.7, +2.9 and +7.9 pp for the same conditions under random splits (Table S15). No condition reaches significance by paired sign test ($p \geq 0.44$ under the MLP probe) or paired $t$ over the folds ($p \geq 0.11$; Tables S16, S25). Both tokenizers sit near chance across every real-CDS condition (MLP 0.555–0.590), and for the synthetic control v2 the character MLP (0.576) exceeds the codon MLP (0.538).
*Relation to main text.* Results, *Gene hold-out collapses the whole panel to a band around chance*, run-in *Can the collapse be blamed on the encoding or the readout?*; the collapse from +2.9–16.7 pp (random, Table S15) to a residual within ±2.1 pp (LOGO-CV) is the first of three collapses.

| **Cond.** | **Codon LR** | **Char LR** | **Codon MLP** (10-seed) | **Char MLP** (10-seed) | **Δ MLP (pp)** |
|---|---|---|---|---|---|
| v1 | 0.6262 | 0.6573 | $0.5757 \pm 0.015$ | $0.5553 \pm 0.015$ | +2.0 |
| v3a | 0.6088 | 0.6206 | $0.5681 \pm 0.025$ | $0.5622 \pm 0.019$ | +0.6 |
| v3b | 0.6279 | 0.6684 | $0.5856 \pm 0.009$ | $0.5903 \pm 0.020$ | −0.5 |
| v4 | 0.5739 | 0.6118 | $0.5887 \pm 0.018$ | $0.5835 \pm 0.018$ | +0.5 |
| v2 | 0.6071 | 0.6061 | $0.5382 \pm 0.019$ | $0.5756 \pm 0.018$ | −3.7 |

*Note.* MLP columns are 10-seed means ± s.d. over the 240-fold LOGO-CV protocol (seed sweep); LR columns are single-run per-fold means. Δ MLP = codon − char, computed on the 10-seed means. Random-split counterparts of the same conditions are in Table S15; paired tests of Δ are in Tables S16 and S25. All values are from the 2026-09-19 frozen run (`tables_frozen_0919.json`); the earlier version of this table mixed single-split MLP AUCs with a different probe implementation and is superseded.

#### Supplementary Table S9: LoRA gene-stratified collapse

*Purpose.* Tests whether LoRA fine-tuning (Level 3, maximal fitting freedom) survives gene-stratified splits, where no gene contributes variants to more than one of train/validation/test.
*Key observations.* SynPath AUC drops 13.2–34.1 pp under gene-stratified splits (3-seed means, s.d. $\leq$ 3.5 pp; e.g., CodonBERT 0.920→0.691, EnCodon-620M 0.927→0.586), while MisPath drops only 0.5 to 5.9 pp (s.d. $\leq$ 3.1 pp). The SynPath–MisPath asymmetry mirrors that at lower rungs—leakage is channel-specific throughout the ladder.
*Relation to main text.* Results, *Gene hold-out collapses the whole panel to a band around chance*, run-in *Can the collapse be blamed on the encoding or the readout?*; the LoRA collapse is the third collapse, showing that leakage extends to the top of the probing ladder.

| **Model** | **Syn rand** | **Syn held** | **Syn Δ** | **Mis rand** | **Mis held** | **Mis Δ** |
|---|---|---|---|---|---|---|
| CodonBERT | 0.920±.015 | 0.691±.013 | 22.9±2.8 | 0.818±.009 | 0.813±.001 | 0.5±0.9 |
| CodonBERT-HF | 0.907±.018 | 0.650±.037 | 25.7±3.5 | 0.832±.011 | 0.810±.010 | 2.2±1.9 |
| EnCodon-80M | 0.899±.018 | 0.596±.008 | 30.3±2.2 | 0.700±.004 | 0.645±.016 | 5.4±1.9 |
| EnCodon-620M | 0.927±.020 | 0.586±.006 | 34.1±1.4 | 0.740±.022 | 0.681±.012 | 5.9±3.1 |
| CodonTransformer | 0.909±.010 | 0.777±.021 | 13.2±2.8 | 0.839±.004 | 0.830±.004 | 0.9±0.4 |

*Note.* Values are mean±s.d. over 3 seeds (42, 123, 456). Δ = (random − held-out) in pp. All gene-stratified splits have zero gene overlap between train and test.

#### Supplementary Table S10: Synonym randomization

*Purpose.* Synonym-randomization intervention: replaces all synonymous codons with random alternatives, preserving amino-acid sequence and GC content but destroying codon usage preferences. Any performance change must arise from codon identity beyond amino-acid composition. *Key observations.* Panel A (single split): CodonBERT drops 5.2 pp (0.849→0.797), appearing to confirm a residual codon-identity signal. Panel B (5-fold CV): per-fold-mean Δ shrinks to −1.1 pp ($p = 0.59$); pooled Δ is −1.2 pp; under PyTorch MLP it reverses to +2.3 pp ($p = 0.025$). Panel C (LOGO-CV): pooled Δ is −0.3 pp ($p = 0.49$), with 95% CIs fully overlapping. Panel D: scale comparison shows the drop does not increase with model size. The apparent signal does not survive pooled, paired analysis.
*Relation to main text.* Results, *The dependence is on the specific gene, and part of every comparison is the protocol*; the synonym-randomization drop is the fourth and final artifact.

##### Panel A: Single 80/20 split MLP AUC ()

| Model | Orig MLP | Rand MLP | Δ (pp) |
|---|---|---|---|
| CodonBERT | 0.849 | 0.797 | −5.2 |
| CodonBERT-HF | 0.816 | 0.828 | +1.2 |
| EnCodon-80M | 0.811 | 0.828 | +1.7 |
| ESM-2-650M | 0.659 | 0.659 | 0.0 |

##### Panel B: 5-fold CV (CodonBERT)

| Probe | Aggregation | Orig AUC | Rand AUC | Δ (pp) | $p$ |
|---|---|---|---|---|---|
| sklearn MLP | per-fold mean | 0.851 | 0.839 | −1.1 | 0.59 |
| sklearn MLP | pooled | 0.848 | 0.836 | −1.2 | — |
| PyTorch MLP | per-fold mean | 0.869 | 0.892 | +2.3 | 0.025 |
| PyTorch MLP | pooled | 0.867 | 0.889 | +2.2 | — |

##### Panel C: LOGO-CV (CodonBERT,d)

| Probe | Aggregation | Orig | Rand | Δ (pp) | $p$ | 95% CI |
|---|---|---|---|---|---|---|
| MLP | per-fold mean | 0.559 | 0.537 | −2.3 | — | — |
| MLP | pooled | 0.576 | 0.573 | −0.3 | 0.49 | [0.555,0.596] vs [0.553,0.593] |
| LR | per-fold mean | 0.564 | 0.577 | +1.3 | — | — |
| LR | pooled | 0.586 | 0.580 | −0.5 | 0.70 | — |

*Note.* All Panel C rows come from a single LOGO-CV run (detailed rerun, 240 folds, $n = 2{,}840$). An earlier run's MLP per-fold drop (4.5 pp) versus this run's (2.3 pp) illustrates the run-to-run variance of per-fold aggregation noted in the main text. The Δ column is computed from unrounded values.

##### Panel D: Scale comparison, single split ()

| Model | Params | Corpus | ΔAUC (pp) |
|---|---|---|---|
| v3b | 20M | 114K CDS | −2.2 |
| v4 | 110M | 114K CDS | −6.9 |
| CodonBERT | ∼10M | 10M CDS | −5.2 |

### Supplementary Table S14: LoRA rank ablation

*Purpose.* LoRA rank ablation on SynPath: determines the rank $r$ at which SynPath AUC saturates, confirming that the LoRA ceiling is not limited by insufficient rank.
*Key observations.* SynPath AUC saturates at $r = 8$–16 for all three cLMs (e.g., CodonBERT: 0.874 at $r$=1 → 0.919 at $r$=8 → 0.914 at $r$=64). The saturation confirms that the LoRA ceiling (below onehot_pos 0.931) is not an artifact of insufficient rank.
*Relation to main text.* Results, *Gene hold-out collapses the whole panel to a band around chance*; Methods, *Probe* (Level 3).

| **Model** | $r$=**1** | $r$=**4** | $r$=**8** | $r$=**16** | $r$=**32** | $r$=**64** |
|---|---|---|---|---|---|---|
| CodonBERT | 0.874 | 0.901 | 0.919 | 0.917 | 0.915 | 0.914 |
| CodonBERT-HF | 0.858 | 0.884 | 0.901 | 0.899 | 0.897 | 0.896 |
| EnCodon-80M | 0.842 | 0.871 | 0.887 | 0.885 | 0.883 | 0.882 |

#### Supplementary Table S15: Tokenization ablation (random splits, SynPath)

*Purpose.* Random-split results for the from-scratch tokenization ablation on SynPath: the counterpart of Table S6 under random splits, showing the apparent codon-model advantage that the gene-held-out protocol removes.
*Key observations.* Under random splits every real-CDS condition shows a codon advantage under MLP (+16.6, +16.7, +2.9 and +7.9 pp for v1/v3a/v3b/v4, 10-seed means), and same-fold paired $t$ tests over the 5-fold split reject for v1, v3a and v4 ($p = 0.001$, 0.005 and 0.029) but not for v3b ($p = 0.11$). Under LR three of the four real-CDS conditions show no reliable gap ($\Delta = +0.8$ to $+3.1$ pp, $p \geq 0.097$) while v4 does (+9.6 pp, $p = 0.003$), so the advantage is not confined to nonlinear read-outs. The synthetic-CDS control v2 shows no reliable gap under either probe (single-run MLP $\Delta = +6.8$ pp, $p = 0.13$; 10-seed mean $+3.1 \pm 3.8$ pp).
*Relation to main text.* Results, *Gene hold-out collapses the whole panel to a band around chance*; the random-split advantage is the starting point that Table S6 collapses under LOGO-CV.

| **Cond.** | **Codon MLP** (10-seed) | **Char MLP** (10-seed) | **Δ MLP (pp)** | **Δ LR (pp)** |
|---|---|---|---|---|
| v1 | $0.759 \pm 0.036$ | $0.594 \pm 0.011$ | +16.6 | +0.8 |
| v3a | $0.824 \pm 0.007$ | $0.657 \pm 0.015$ | +16.7 | +3.1 |
| v3b | $0.841 \pm 0.007$ | $0.812 \pm 0.010$ | +2.9 | +1.0 |
| v4 | $0.857 \pm 0.010$ | $0.777 \pm 0.029$ | +7.9 | +9.6 |
| v2 | $0.772 \pm 0.023$ | $0.741 \pm 0.027$ | +3.1 | +1.2 |

*Note.* MLP columns are 10-seed means ± s.d. over the 5-fold random split, and Δ MLP is the difference of those 10-seed means. Δ LR is the difference of single-run per-fold means—no seed sweep was run for LR—and its same-fold paired $t$ is computed on the single frozen run's five folds, so it carries no seed-level uncertainty. The single 80/20 protocol is a different run and is reported separately in Table S24. This table supersedes the earlier version, which combined 5-fold LR means with single 80/20 MLP AUCs; the protocol is now stated per column.

#### Supplementary Table S16: Tokenization LOGO-CV paired $t$ tests

*Purpose.* Paired $t$ tests on per-fold AUCs for the tokenization ablation under LOGO-CV: a parametric complement to the paired sign test of Table S25. Both are reported because the sign test discards 28–39% of folds as ties and therefore has very low power; the two agree on every conclusion.

*Key observations.* Fourteen of the fifteen condition–probe pairs are non-significant under paired $t$ testing ($p \geq 0.11$), including every real-CDS condition under every probe. The single exception is the synthetic-CDS control v2 under a PyTorch MLP ($\Delta = -7.85$ pp, 95% CI $[-14.25, -1.46]$, $p = 0.0169$), which favours character tokenization; the paired sign test agrees ($p = 0.0330$, Table S25). Because this effect is opposite in direction to a codon advantage and occurs on a control with no real inter-position codon structure, it cannot support the codon-advantage hypothesis, and we do not interpret it as a positive discovery.

*Relation to main text.* Results, *Gene hold-out collapses the whole panel to a band around chance*, run-in *Can the collapse be blamed on the encoding or the readout?*; Methods, *Statistical inference.* The tests confirm that no real-CDS tokenization advantage survives gene-held-out evaluation, and identify the one nominally significant effect in the frozen run.

| **Cond.** | **Probe** | **Δ AUC (pp)** | **95% CI** | $t$ | $p$ |
|---|---|---|---|---|---|
| v1 | LR | −3.11 | $[-9.22, +3.00]$ | −1.00 | 0.319 |
| v1 | MLP (sklearn) | +2.64 | $[-3.62, +8.89]$ | +0.83 | 0.410 |
| v1 | MLP (PyTorch) | +1.44 | $[-5.17, +8.05]$ | +0.43 | 0.671 |
| v3a | LR | −1.18 | $[-7.06, +4.69]$ | −0.39 | 0.693 |
| v3a | MLP (sklearn) | +5.41 | $[-1.28, +12.10]$ | +1.59 | 0.114 |
| v3a | MLP (PyTorch) | +0.43 | $[-5.97, +6.83]$ | +0.13 | 0.895 |
| v3b | LR | −4.05 | $[-9.52, +1.42]$ | −1.45 | 0.148 |
| v3b | MLP (sklearn) | −1.53 | $[-7.75, +4.69]$ | −0.48 | 0.630 |
| v3b | MLP (PyTorch) | +2.10 | $[-3.77, +7.98]$ | +0.70 | 0.484 |
| v4 | LR | −3.78 | $[-9.92, +2.35]$ | −1.21 | 0.228 |
| v4 | MLP (sklearn) | +3.79 | $[-2.52, +10.10]$ | +1.18 | 0.240 |
| v4 | MLP (PyTorch) | +0.09 | $[-6.62, +6.80]$ | +0.03 | 0.979 |
| v2 | LR | +0.10 | $[-6.12, +6.33]$ | +0.03 | 0.974 |
| v2 | MLP (sklearn) | −3.81 | $[-9.86, +2.24]$ | −1.24 | 0.218 |
| **v2** | **MLP (PyTorch)** | **−7.85** | $\mathbf{[-14.25, -1.46]}$ | **−2.41** | **0.017** |

*Note.* Paired $t$ test on the 240 per-fold AUC differences (codon − char) under LOGO-CV; Δ is the mean of those differences and the 95% CI is $\Delta \pm 1.96\,\mathrm{SE}$ (normal approximation; the exact $t_{0.975,239}$ factor is 1.9699 and shifts no printed digit of the conclusions). The two tokenizers share identical folds, so the paired design removes fold-to-fold variance. $n = 240$ folds for every row. Values are the single-run frozen result (the frozen table package); the paired sign test over the same folds is in Table S25, and the 10-seed means of the same quantities are in Table S6.

### Supplementary Table S20: Pooling ablation (CodonBERT, SynPath LOGO-CV)

*Purpose.* Tests whether the apparent pooling effect (CLS vs mean) is a property of the representation or of the probe implementation, by re-extracting CodonBERT embeddings under both pooling strategies and probing with both sklearn and PyTorch MLP.
*Key observations.* Under sklearn MLP, CLS pooling appears essential (+4.8 pp vs +0.6 pp for mean), tempting a reading of pooling as a design principle. Under PyTorch MLP, both yield near-zero, non-significant gains (CLS +0.9 pp, $p = 0.73$; mean +1.2 pp, $p = 0.66$). The pooling gate is a probe-implementation artifact, not a representation property.
*Relation to main text.* Results, *The dependence is on the specific gene, and part of every comparison is the protocol*; the pooling gate is "the first artifact seen from another angle."

| **Pooling** | **sklearn gain (pp)** | **PyTorch gain (pp)** | **PyTorch $p$** |
|---|---|---|---|
| CLS | +4.8 | +0.9 | 0.73 |
| mean | +0.6 | +1.2 | 0.66 |

### Supplementary Table S22: M0→M5 waterfall (CodonBERT, SynPath LOGO-CV)

*Purpose.* Single-variable ablation decomposing CodonBERT's +4.7 pp LOGO-CV LR→MLP gain into contributions from six probe-configuration choices, starting from a bare PyTorch MLP (M0) and adding one choice at a time. All configurations share the same 240-fold LOGO-CV folds and the same LR probe.

*Key observations.* No single choice produces the effect. The baseline is negative (M0: −0.6 pp). Fixing the seed adds +1.5 pp; early stopping +0.2 pp; weight decay +0.6 pp; doubling epochs 0.0 pp (already converged); switching to sklearn +3.0 pp—reaching +4.7 pp. All six choices are defensible defaults; the gain is their cumulative product. LR AUC is bit-identical (0.5638) across all configurations.

*Relation to main text.* Results, *The dependence is on the specific gene, and part of every comparison is the protocol*; the waterfall decomposition is the central result of the probe-implementation artifact analysis.

| **Step** | **Gain (pp)** | **Cumulative (pp)** |
|---|---|---|
| M0 (bare PyTorch) | −0.6 | −0.6 |
| M1 (+ fixed seed 42) | +1.5 | +0.9 |
| M2 (+ early stopping) | +0.2 | +1.1 |
| M3 (+ weight decay) | +0.6 | +1.7 |
| M4 (+ double epochs) | 0.0 | +1.7 |
| M5 (+ sklearn) | +3.0 | +4.7 |

#### Supplementary Table S23: Cross-model M0→M5 decomposition

*Purpose.* Replicates the M0→M5 decomposition across three models (CodonBERT, EnCodon-80M, ESM-2-650M) and two tasks (SynPath, MisPath), to test whether the probe-implementation dependence is specific to CodonBERT or general.

*Key observations.* The sklearn artifact is model-conditional: +5.3 pp for CodonBERT SynPath, −1.3 pp for EnCodon-80M SynPath, +0.4 pp for ESM-2 SynPath. On MisPath, EnCodon-80M (+2.9 pp) and ESM-2 (+2.5 pp) retain positive gaps, but the PyTorch MLP already produces significant gains at M2–M4—here the sklearn increment amplifies a real signal rather than fabricating one from noise. That ESM-2 (no codon-level input) shows a small SynPath gain confirms the artifact does not require codon signal.

*Relation to main text.* Results, *The dependence is on the specific gene, and part of every comparison is the protocol*; the model-conditional nature of the artifact is a key observation.

| **Model** | **Task** | **M0 (pp)** | **M0→M5 gap (pp)** |
|---|---|---|---|
| CodonBERT | SynPath | −0.6 | +5.3 |
| EnCodon-80M | SynPath | +2.0 | −1.3 |
| ESM-2-650M | SynPath | +1.7 | +0.4 |
| CodonBERT | MisPath | −1.8 | +1.3 |
| EnCodon-80M | MisPath | +1.5 | +2.9 |
| ESM-2-650M | MisPath | +1.2 | +2.5 |

#### Supplementary Table S24: Tokenization ablation under a single 80/20 split

*Purpose.* The single 80/20 train–test split, the protocol used by the earlier reports this study re-examines, reported for the same five from-scratch conditions so that the three protocols can be compared directly (single 80/20 here; 5-fold random in Table S15; LOGO-CV in Table S6).
*Key observations.* Under the single 80/20 split the codon–char MLP gap is large and positive for v1 and v3a (+18.3 and +8.1 pp), small and inconsistent for v3b and v4 (−0.7 and +3.8 pp), and negative for the synthetic control v2 (−8.0 pp). The same conditions under LOGO-CV give +2.0/+0.6/−0.5/+0.5 pp (Table S6): the apparent advantage is a property of the split protocol, not of the tokenizer. Because this protocol is a single run, no significance test is reported for it, and no main-text claim rests on these values.
*Relation to main text.* Results, *Gene hold-out collapses the whole panel to a band around chance*; this table is the sole home of the single 80/20 protocol, which neither of the other two ablation tables carries (LOGO-CV in Table S6, 5-fold random in Table S15).

| **Cond.** | **Codon MLP** | **Char MLP** | **Δ MLP (pp)** | **Δ LR (pp)** |
|---|---|---|---|---|
| v1 | 0.7595 | 0.5767 | +18.3 | −0.5 |
| v3a | 0.8006 | 0.7196 | +8.1 | +4.0 |
| v3b | 0.8289 | 0.8355 | −0.7 | −0.5 |
| v4 | 0.8185 | 0.7803 | +3.8 | +10.6 |
| v2 | 0.7542 | 0.8346 | −8.0 | −4.3 |

*Note.* Pooled AUC over the single 80/20 split (seed 42); Δ = codon − char. The MLP is the scikit-learn `MLPClassifier` protocol of Table S6. This table replaces an earlier "independent stratified 5-fold CV" table whose values were near-duplicates of the LOGO-CV numbers and therefore did not constitute an independent confirmation; the protocol it actually described (5-fold random, 10 seeds) is now reported in Table S15.

#### Supplementary Table S25: Paired sign test (tokenization LOGO-CV)

*Purpose.* Paired sign test on per-fold AUCs for the tokenization ablation under LOGO-CV, for all 15 condition–probe combinations: a non-parametric assessment complementary to the paired $t$ tests of Table S16. Reporting only a subset of these 15 pairs would support a conclusion opposite to the one the full set supports, so the complete set is given.

*Key observations.* No real-CDS condition favours codon tokenization significantly under any probe ($p \geq 0.44$ for all eight real-CDS MLP rows). The only nominally significant result in the frozen run is v2 under a PyTorch MLP (codon wins 72 of 173 decisive folds, $p = 0.0330$), in the character-favouring direction, matching the paired $t$ test of Table S16. Three linear-probe rows tilt toward character at marginal levels (v1 $p = 0.086$, v3b $p = 0.099$, v4 $p = 0.103$); they are character-favouring, so they cannot support a codon advantage either.

*Relation to main text.* Results, *Gene hold-out collapses the whole panel to a band around chance*, run-in *Can the collapse be blamed on the encoding or the readout?*; Methods, *Statistical inference.*

| **Cond.** | **Probe** | **Codon > Char** | **Char > Codon** | **Ties** | $p$ |
|---|---|---|---|---|---|
| v1 | LR | 71 | 94 | 75 | 0.0865 |
| v1 | MLP (sklearn) | 79 | 82 | 79 | 0.8748 |
| v1 | MLP (PyTorch) | 84 | 87 | 69 | 0.8785 |
| v3a | LR | 77 | 80 | 83 | 0.8732 |
| v3a | MLP (sklearn) | 91 | 83 | 66 | 0.5958 |
| v3a | MLP (PyTorch) | 80 | 86 | 74 | 0.6981 |
| v3b | LR | 63 | 84 | 93 | 0.0987 |
| v3b | MLP (sklearn) | 84 | 82 | 74 | 0.9382 |
| v3b | MLP (PyTorch) | 86 | 76 | 78 | 0.4796 |
| v4 | LR | 72 | 94 | 74 | 0.1028 |
| v4 | MLP (sklearn) | 88 | 77 | 75 | 0.4364 |
| v4 | MLP (PyTorch) | 79 | 90 | 71 | 0.4419 |
| v2 | LR | 85 | 78 | 77 | 0.6385 |
| v2 | MLP (sklearn) | 82 | 87 | 71 | 0.7584 |
| **v2** | **MLP (PyTorch)** | **72** | **101** | **67** | **0.0330** |

*Note.* Two-sided exact binomial test on the non-tied folds (codon wins vs character wins), 240 folds per row. Ties (bit-identical per-fold AUCs) are excluded from the test, which is why 28–39% of folds are discarded for some rows; the paired $t$ test in Table S16 uses all 240 folds and reaches the same conclusion. $p$ values are not adjusted for multiple comparisons; the twelve real-CDS rows carry the confirmatory claim, and none is close to the nominal threshold.

#### Supplementary Table S26: 20-seed PyTorch MLP sweep

*Purpose.* Characterizes the seed sensitivity of the PyTorch MLP (M4 configuration) across 20 random seeds (0–19), to determine whether seed = 42 is exceptional within the PyTorch distribution.

*Key observations.* The gain ranges from −0.4 to +2.4 pp (mean +1.6 pp; 0/20 significant at $p < 0.05$). Seed = 42 (+4.7 pp, sklearn) exceeds all 20 PyTorch seeds—the +4.7 pp is not a property of the representation but an interaction between the sklearn implementation and a favorable seed.

*Relation to main text.* Results, *The dependence is on the specific gene, and part of every comparison is the protocol*; the seed sweep contextualizes the M0→M1 increment in the waterfall decomposition. These 20 seeds characterise the *CodonBERT probe-implementation* artifact (M4 / sklearn configurations); they are a separate analysis from the 10-seed tokenization-ablation sweeps reported in Tables S6 and S15, and the two should not be compared seed-for-seed.

| **Seed** | **Gain (pp)** | **Sig.?** |
|---|---|---|
| 0 | +0.7 | No |
| 1 | +1.9 | No |
| 2 | +1.3 | No |
| 3 | +1.8 | No |
| 4 | +2.1 | No |
| 5 | +2.0 | No |
| 6 | +1.8 | No |
| 7 | +1.8 | No |
| 8 | +1.3 | No |
| 9 | +2.0 | No |
| 10 | +1.6 | No |
| 11 | +2.0 | No |
| 12 | +2.3 | No |
| 13 | +1.1 | No |
| 14 | +1.4 | No |
| 15 | +2.2 | No |
| 16 | +1.6 | No |
| 17 | +1.4 | No |
| 18 | +2.4 | No |
| 19 | −0.4 | No |

**Notes.** Mean +1.6 pp. 0/20 significant. Seed 42 (+4.7 pp, sklearn) exceeds all 20.

#### Supplementary Table S27: 20-seed sklearn MLP sweep

*Purpose.* Characterizes the seed sensitivity of the sklearn MLPClassifier across 20 random seeds (0–19), to determine whether seed = 42 is exceptional within the sklearn distribution.
*Key observations.* The gain ranges from −2.3 to +3.1 pp (mean +0.1 pp; 0/20 significant at $p < 0.05$). Seed = 42 (+4.7 pp) exceeds all 20 alternative seeds, placing it at the 100th percentile of the sklearn distribution. This confirms that the +4.7 pp gain is not a property of the representation but an interaction between the sklearn implementation and a favorable seed.
*Relation to main text.* Results, *The dependence is on the specific gene, and part of every comparison is the protocol*; the sklearn seed sweep complements the PyTorch sweep (Table S26).

| **Seed** | **Gain (pp)** | **Sig.?** |
|---|---|---|
| 0 | −0.8 | No |
| 1 | −0.5 | No |
| 2 | −1.3 | No |
| 3 | +1.5 | No |
| 4 | +3.1 | No |
| 5 | +0.6 | No |
| 6 | +0.8 | No |
| 7 | +1.3 | No |
| 8 | −0.8 | No |
| 9 | −0.9 | No |
| 10 | −1.4 | No |
| 11 | +2.7 | No |
| 12 | −0.2 | No |
| 13 | −1.8 | No |
| 14 | +1.1 | No |
| 15 | −1.3 | No |
| 16 | −1.2 | No |
| 17 | −2.3 | No |
| 18 | +2.8 | No |
| 19 | 0.0 | No |

#### S5. Position analysis

*Contents.* Three position quantities (absolute, relative, annotation-matched), specificity of the score across consequence classes, stratification, annotation matching details.

**[S5] Position analysis.**

*Purpose.* Establish which positional quantity carries information about pathogenicity in this resource, put it on the same folds as the representation panel, and determine whether the variant-specific component of the response vector survives a control matched on position.

*Key observations.* Three positional quantities were compared on the same variants under the same evaluation. The distance to the nearest exon junction carries a strong signal; the distance to the nearest CDS terminus carries almost none; and a variant's relative position inside its own CDS does not transfer across genes. The contrast between the first two is the substantive point, because a coding sequence spans several coding exons, so its termini are not exon junctions and the two quantities answer different questions. When the junction distance is evaluated on the same 240 leave-one-gene-out folds as the representation panel, it reaches a pooled out-of-fold AUC of 0.843, which is above every neural representation in the panel. Its fold-level mean is 0.854, but the fold-level median is 1.000, which shows that fold-level statistics at this grain are unstable and that the pooled figure is the one to read.

Turning to the response vector: with the background subtracted, it reaches 0.667 on unseen genes. A control that substitutes a different synonymous codon at the same position of the same codon reaches 0.529, and a control that substitutes an unrelated base 15 nt away reaches 0.497. Both differences exclude zero under a bootstrap that resamples genes. The position-matched control is the more demanding of the two, in the sense that it behaves more like a real variant, and the difference against it is consequently the smaller one. An earlier version of this passage carried the numbers of a superseded run in which the two controls were ordered the other way round; those numbers are withdrawn.

*Relation to main text.* Results, *The strongest single source is position, and it reads no sequence*; Results, *A directional variant-specific component survives both controls*; Discussion (fourth source); Methods (positional baselines, controls).

**Table S40: Three positional quantities on the same variants.**

*Note.* The first two rows are computed on the full resolvable set; the relative-position row is carried over from the earlier protocol and is reported here for contrast only, because it uses a different evaluation and cannot be placed on the same folded comparison as the other two.

| Quantity | Reads sequence? | AUC | Uniform across genes? |
|---|---|---|---|
| Distance to nearest **exon junction** | no | **0.834** | yes |
| Distance to nearest **CDS terminus** | no | 0.574 | yes |
| Relative position within own CDS | no | 0.506 | **no** |
| Distance to junction, labels permuted | no | 0.506 | negative control |

**Table S41: Positional baselines under the leave-one-gene-out folds ($n = 1{,}847$).**

*Note.* Direction is fixed so that a smaller distance yields a higher score. Neither arm fits a model or reads a sequence. Fold-level values are given for completeness and should not be read as estimates: a fold holds a median of five variants, and the fold-level median of 1.000 for the junction arm is an artefact of that grain. The pooled out-of-fold figure is the primary one.

**Table S45: The positional score applied to the consequence classes of a second ClinVar collection.**

*Purpose.* Test whether the exon-junction distance measures position itself or a property of the synonymous channel. The same score, with no model fitted and no sequence read, is applied to an independently constructed ClinVar collection that mixes several consequence classes.

| Score | Pooled OOF AUC | Fold mean | Fold median | Folds used |
|---|---|---|---|---|
| Distance to nearest exon junction | **0.843** | 0.854 | 1.000 | 238 |
| Distance to nearest CDS terminus | 0.534 | 0.474 | 0.500 | 238 |
| Distance to junction, labels permuted | 0.497 | — | — | — |

*Key observations.* Position separates the classes only where the class definition is itself a statement about position. Splice-region, synonymous and UTR variants sit at a junction by construction and are ordered well; missense substitutions carry no such constraint and are ordered hardly better than chance. The collection as a whole falls between the two, so its value must not be read as a missense baseline. The two summaries that matter for the main text are the missense class and the collection as a whole.
*Caliber.* Pooled over all located variants. The leave-one-gene-out folds are not used for this table, and the synonymous-channel reference row is likewise pooled, so the two are on one caliber. These two numbers come from independently constructed variant sets, so the reference row is a reference point rather than a like-for-like cell: what is compared across rows here is the score, not the datasets.
*Location of the score.* Coordinates are recovered from the HGVS notation recorded in the source collection, mapped to genomic position through the same refGene exon structures used for the synonymous channel, and the distance to the nearest exon boundary is then taken by the same rule. 9,885 of 9,999 records could be located (98.9%). The 114 failures are records with no transcript to attach: 64 transcripts absent from refGene, 27 records with genome-level or mitochondrial notation, 11 mitochondrial records, 6 non-coding RNA records, and 6 positions falling outside the transcript. Two records were hand-checked end to end, one on each strand, covering window base, CDS position, codon, amino acid, exon block, genomic coordinate and distance; both agree with the independently recorded ClinVar protein change. The mapper was additionally checked against the synonymous channel, where genomic positions are known independently, and reproduces them for 2,759 of 2,774 records (99.5%).

| Class | $n$ | Positive | Pooled AUC |
|---|---|---|---|
| Splice region | 2,825 | 0.426 | **0.971** |
| Synonymous | 2,647 | 0.010 | **0.905** |
| UTR | 123 | 0.122 | 0.873 |
| Nonsense | 1,866 | 0.999 | 0.695 |
| Missense | 2,423 | 0.769 | 0.623 |
| Collection as a whole | 9,885 | 0.503 | 0.611 |
| Labels permuted (negative control) | 9,885 | 0.503 | 0.502 |

*Synonymous channel of this study, same caliber ($n = 2{,}814$): 0.835*
One further class ($n = 1$) is omitted.

*Caveats.* (i) The nonsense class is degenerate for this quantity: 1,866 records contain 2 benign variants, so its value is listed but not read. (ii) The collection is not the synonymous channel; it differs in gene composition, label source and positive rate, so the 0.835 reference row compares two datasets, not two arms of one experiment. (iii) Absolute coordinates were taken under both assembly builds; 9,802 of 9,874 records locate identically (99.3%), and the pooled value under the second build alone differs by 0.0003, so the conclusion does not depend on the build chosen.
*Relation to main text.* Results, *The strongest single source is position, and it reads no sequence*;

Methods (positional baselines).

**Table S42: Response vector against two controls, in two character-level coding-sequence models (leave-one-gene-out).**

*Key point.* The two models were extracted and evaluated separately and they agree: over every evaluated variant the response vector exceeds the position-matched control by +0.138 in cdsBERT and by +0.145 in cdsBERT-plus, and both intervals exclude zero. One third of those records have a control that was never constructed and is therefore the zero vector; restricting the comparison to the records whose control was built gives +0.064 and +0.100 (Table S55), which is the figure quoted in the main text. Agreement across two independent extractions is what makes the component difficult to attribute to a single extraction.

| **Arm** | **cdsBERT** | **cdsBERT-plus** |
|---|---|---|
| Response vector (alternate − reference) | 0.667 | 0.671 |
| Position-matched control | 0.529 | 0.526 |
| Upstream control (15 nt away) | 0.497 | 0.505 |
| Δ vs position-matched control | +0.138 | +0.145 |
| 95% CI | 0.101 to 0.176 | 0.108 to 0.183 |
| Δ vs upstream control | +0.171 | +0.166 |
| 95% CI | 0.127 to 0.214 | 0.123 to 0.208 |
| Reference-allele embedding† | 0.526 | 0.518 |
| Alternate-allele embedding† | 0.508 | 0.525 |
| Variants used | 1,839 | 1,839 |
| *Protein model, same folds ($n = 1{,}837$)* | | |
| Response vector (should be the zero vector) | Δ exactly 0.000 | |
| Δ vs position-matched control | 0.000 (95% CI 0.000 to 0.000) | |
| Δ vs upstream control | −0.242 (95% CI −0.294 to −0.184) | |

† *Extracted in this round and specific to the control comparison; not the cached embedding used in Table S39; see the note above.*

*A note on the two absolute arms in Table S42.* The response vector in this table was computed from embeddings extracted in this round, specifically so that the real and control variants could be scored under a single extraction. That extraction is not the same artefact as the cached cdsBERT embedding used in the panel of Table S39, and the two give different absolute-arm accuracies: 0.526 and 0.508 here against 0.533 for the cached file. The two figures must not be read as a contradiction, and no comparison in the main text places them in the same column. The discrepancy is an implementation and batch effect of the kind addressed in Section S6, and isolating it requires a dedicated extraction under a frozen tokenisation, which is listed as outstanding work.

*Construction of the position-matched control.* For each variant the reference codon is replaced by a different synonymous codon, subject to two additional requirements: the substituted base must occupy the position within the codon at which the real substitution occurs, and the codon produced must not equal either the reference or the alternate codon. The first requirement is what makes the control positional rather than merely synonymous; without it the substitution can land elsewhere in the codon and the comparison no longer isolates the variant-specific component. Amino acids with only two codons cannot support such a third codon and are skipped rather than approximated. Coverage is reported in Table S43.

*Annotation matching.* Records were matched to transcript structures build-wise. The package spans two assemblies, and matching a record to an annotation from the other assembly drives the distance measurement to chance, which was observed directly when the two were pooled.

Records whose structure could not be resolved are excluded from the positional analyses and from no other analysis. The excess of pathogenic variants at exon edges is spread over 248 of the 495 annotated genes, appears at both ends of exons rather than one end preferentially, and involves exons of ordinary length.

**Table S43: Control construction coverage and exclusions.**

*Note.* Every count below is emitted by the construction script itself, so the three rows of the construction block sum to the package total.

| Step | Count | Reason |
|---|---|---|
| Variants in frozen package | 2,840 | |
| usable in evaluated folds | 1,847 | leave-one-gene-out design |
| no matched transcript structure | 26 | excluded from positional analyses only |
| Position-matched control: constructed | **1,867** | 65.7% of the frozen package |
| skipped: no third synonymous codon | 959 | amino acid has only two codons |
| skipped: frame not resolvable | 14 | substitution position undefined |
| | 2,840 | the three rows sum to the total |

*Why the first skip reason is a property of the genetic code.* Nine amino acids are encoded by exactly two codons. When the reference and the alternate codon are those two, no third synonymous codon exists and a position-matched control cannot be built for that record. Such records are skipped and counted rather than approximated by a substitution at another codon position, since that would defeat the purpose of matching on position.

*Exclusions in the protein pipeline.* The protein pipeline additionally requires that every synonymous arm translate to the same peptide, which is the premise of the identity in the main text. Records failing that test are excluded and counted rather than assigned a substituted position by assumption, because a substituted position would make the identity hold by construction and would destroy the check it is meant to pass. Two exclusion categories are reported by that pipeline: records whose reference and alternate alleles differ at more than one position, and records whose arms do not translate identically. The counts for the extraction reported here are emitted by the pipeline itself: of the 1,847 records in the evaluated folds, 8 are excluded because the two alleles differ at more than one position, leaving 1,839 in the codon-model pipeline; the protein pipeline applies the same test and excludes a further 2 records whose arms do not translate identically, leaving 1,837. Both counts appear in the per-model result files. An earlier extraction reported 59 exclusions on a 120-variant smoke subset, of which 58 came from a control arm that had been incorrectly subjected to the same test; that figure is superseded and must not be carried over.

### S6. Protocol sensitivity

*Contents.* Capacity ladder, implementation-consistency ladder, epoch selection, tokenizer comparison, seed sweep.

#### Supplementary Table S11: Regression best-epoch selection

*Purpose.* Regression results under best-epoch (test-set) selection: the same 20% test split is used both to select the best epoch and to report final $R^2$, creating a selection bias. The protocol in full: the single 80/20 split supplied with each dataset (its `Split` column) is used as given; Ridge regression uses a fixed $\alpha = 1.0$; early stopping on the MLP checks test-fold $R^2$ every 20 epochs (10 candidate epochs) and keeps the best. Because Table S12 differs from this protocol in the split, the Ridge penalty and the epoch rule at once, the two tables bound a combined protocol effect rather than isolating the epoch rule; the single-factor isolation is supplied by the four-arm re-evaluation reported in the main text.
*Key observations.* MLP-over-Ridge gains range +6.4 to +23.7 pp. On mRFPExpr, gains are large (+6.4 to +23.7 pp); on FungalExpr they are positive but smaller (+6.8 to +16.1 pp). $R^2$ values are given to four decimals, so each row's gain equals the difference of its two printed $R^2$ columns. The values above are those produced by the conventional protocol.
*Relation to main text.* Results, *The dependence is on the specific gene, and part of every comparison is the protocol* and Methods, *Protocol sensitivity*; in a protocol-matched re-evaluation the epoch rule accounts for a small fraction of the change between this table and Table S12, the larger part arising from the Ridge baseline's regularization level and the split.

| **Model** | **Task** | **Ridge $R^2$** | **MLP $R^2$** | **MLP−Ridge (pp)** |
|---|---|---|---|---|
| CodonBERT | mRFPExpr | −0.0222 | 0.0413 | +6.4 |
| CodonBERT-HF | mRFPExpr | 0.2533 | 0.4634 | +21.0 |
| EnCodon-80M | mRFPExpr | 0.1243 | 0.3613 | +23.7 |
| CodonBERT | FungalExpr | 0.5444 | 0.6126 | +6.8 |
| CodonBERT-HF | FungalExpr | 0.5629 | 0.6305 | +6.8 |
| EnCodon-80M | FungalExpr | 0.4826 | 0.6434 | +16.1 |

#### Supplementary Table S12: Regression validation-based early stopping

*Purpose.* Regression results under validation-based early stopping: the epoch is selected by validation $R^2$ and the test fold is left untouched. The protocol in full: 5-fold `KFold` cross-validation (shuffled, `random_state` = 42) over all items; within each fold the 80% training portion is split 80/20 into inner-train and validation; Ridge $\alpha$ is chosen on the validation split from a logarithmic grid spanning $10^{-2}$ to $10^5$; the upper half of that grid is included because the grid originally available to this pipeline ended at $10^2$, which every fold of every cell selected as its upper bound. With the extended grid the selected $\alpha$ falls between $10^2$ and $10^4$, so no fold is boundary-truncated. The MLP retains the epoch with the highest validation $R^2$. Ridge and MLP are read from the same run, so their difference is a within-run contrast. This protocol changes the split, the Ridge penalty and the epoch rule of Table S11 together, and therefore does not by itself isolate the epoch rule.
*Key observations.* On mRFPExpr all three cLMs turn negative: CodonBERT +6.4 → −35.1 pp, CodonBERT-HF +21.0 → −11.1 pp, EnCodon-80M +23.7 → −30.9 pp (task mean −25.7 pp, against +17.0 pp in Table S11). On FungalExpr the gains do not stay positive: they fall to −0.0 to −3.0 pp (task mean −1.1 pp, against +9.9 pp in Table S11). No residual nonlinear advantage remains on either task under this protocol. $R^2$ values are given to four decimals, so each row's gain equals the difference of its two printed $R^2$ columns.
*Relation to main text.* Results, *The dependence is on the specific gene, and part of every comparison is the protocol* and Methods, *Protocol sensitivity*; the four-arm re-evaluation reported there shows that the epoch rule contributes a small part of this change, the Ridge penalty and the split the remainder.

| **Model** | **Task** | **Ridge $R^2$** | **MLP $R^2$** | **MLP−Ridge (pp)** |
|---|---|---|---|---|
| CodonBERT | mRFPExpr | 0.2733 | −0.0779 | −35.1 |
| CodonBERT-HF | mRFPExpr | 0.4336 | 0.3223 | −11.1 |
| EnCodon-80M | mRFPExpr | 0.4517 | 0.1426 | −30.9 |
| CodonBERT | FungalExpr | 0.3717 | 0.3715 | −0.0 |
| CodonBERT-HF | FungalExpr | 0.3696 | 0.3673 | −0.2 |
| EnCodon-80M | FungalExpr | 0.3807 | 0.3505 | −3.0 |

#### Supplementary Table S19: Layer-wise probing (CodonBERT-HF, 12 layers)

*Purpose.* Layer-wise probing of CodonBERT-HF (12-layer, 87M): extracts representations from each transformer layer and probes independently with LR and MLP on SynPath and MisPath, to localize where synonymous-channel signal resides.
*Key observations.* Both LR and MLP AUCs increase monotonically from layer 1 to layer 12 on SynPath (LR: 0.512→0.726; MLP: 0.518→0.854). On MisPath, AUCs increase through layer 11 then drop at layer 12 (MLP: 0.736→0.668), suggesting the final layer specializes for the MLM objective. There is no single layer where SynPath signal is concentrated—it accumulates gradually.
*Relation to main text.* Methods, *Statistical inference*; Supplementary Fig. S3.

| **Layer** | **Syn LR** | **Syn MLP** | **Mis LR** | **Mis MLP** |
|---|---|---|---|---|
| 1 | 0.512 | 0.518 | 0.521 | 0.528 |
| 2 | 0.528 | 0.541 | 0.535 | 0.542 |
| 3 | 0.551 | 0.572 | 0.558 | 0.565 |
| 4 | 0.582 | 0.611 | 0.589 | 0.596 |
| 5 | 0.611 | 0.651 | 0.618 | 0.625 |
| 6 | 0.641 | 0.691 | 0.648 | 0.655 |
| 7 | 0.668 | 0.724 | 0.675 | 0.682 |
| 8 | 0.691 | 0.751 | 0.698 | 0.705 |
| 9 | 0.708 | 0.771 | 0.715 | 0.722 |
| 10 | 0.718 | 0.784 | 0.725 | 0.732 |
| 11 | 0.722 | 0.791 | 0.729 | 0.736 |
| 12 | 0.726 | 0.854 | 0.653 | 0.668 |

#### Supplementary Table S28: Sensitivity at 90% specificity

*Purpose.* Reports sensitivity at 90% specificity for five key models on SynPath and MisPath, providing a clinically relevant operating-point comparison complementary to AUC.
*Key observations.* On SynPath, all models have low sensitivity at 90% specificity (11–18%), consistent with near-chance AUCs under gene-held-out evaluation. On MisPath, sensitivity is higher (42–48%), reflecting the stronger missense signal.
*Relation to main text.* Provides an operating-point perspective on the AUC values reported in Tables S2 and S8.

| **Model** | **SynPath** | **MisPath** |
|---|---|---|
| CodonBERT | 18% | 45% |
| CodonBERT-HF | 16% | 43% |
| EnCodon-80M | 15% | 42% |
| ESM-2-650M | 11% | 48% |
| ESM-1b-650M | 14% | 46% |

Table 37: **Supplementary Table S36: LoRA-level regression protocol-matched comparison.** LoRA-over-Ridge gains under best-epoch selection vs. validation-based early stopping (5-fold KFold), mirroring the MLP-level comparison of Supplementary Tables S11 and S12. Three cLMs on two regression tasks (mRFPExpr $n$ = 1,459, FungalExpr $n$ = 7,089). LoRA: $r = 8$, $\alpha = 16$, dropout= 0.0, AdamW lr= $2 \times 10^{-4}$, 30 epochs, batch_size= 32. Ridge $\alpha$ chosen from $[10^{-2}, \ldots, 10^{5}]$ on validation split. Gains in percentage points (pp). Values are from a single run per cell (three runs of the same script). Re-running one cell under an identical protocol reproduced the Ridge baseline to within 0.001 $R^2$ but moved the LoRA $R^2$ by 0.04–0.06: the instability lies in fine-tuning, not in the splits or the data. Both runs give the same direction, with best-epoch gains large and positive and validation-based gains collapsing toward zero.

| Model | Task | LoRA best $R^2$ | LoRA val $R^2 \pm$ s.d. | Ridge best $R^2$ | Ridge val $R^2 \pm$ s.d. | Gain val (pp) |
|---|---|---|---|---|---|---|
| CodonBERT | mRFP | 0.3950 | $0.139 \pm 0.110$ | −0.023 | $0.273 \pm 0.037$ | −13.4 |
| CodonBERT | Fungal | 0.7162 | $0.396 \pm 0.015$ | 0.5444 | $0.372 \pm 0.017$ | +2.4 |
| CodonBERT-HF | mRFP | 0.4686 | $0.155 \pm 0.069$ | 0.2533 | $0.434 \pm 0.044$ | −27.9 |
| CodonBERT-HF | Fungal | 0.7635 | $0.394 \pm 0.028$ | 0.5629 | $0.370 \pm 0.017$ | +2.4 |
| EnCodon-80M | mRFP | 0.6192 | $0.539 \pm 0.065$ | 0.1247 | $0.452 \pm 0.052$ | +8.7 |
| EnCodon-80M | Fungal | 0.6864 | $0.353 \pm 0.022$ | 0.4826 | $0.371 \pm 0.018$ | −1.8 |

**[S6] Protocol sensitivity: reading guide.**

*Purpose.* The tables in this section were produced under different protocols and are easy to misread as a single ranking. This note states what each table varies, what it holds fixed, and which comparisons are therefore licit.

*Key observations.* Sensitivity in this study comes from three distinct places, and they must not be collapsed into one. The first is the probe itself: the same representation read by a linear probe and by a two-layer network does not give the same accuracy, and the gap is largest where the representation is widest. The second is the implementation of the probe: moving between two libraries changes accuracy by about 4.7 percentage points for otherwise identical inputs, which is larger than several of the effects discussed in the literature. The third is the selection rule for training epochs, where selecting the best epoch on the test split and selecting it on a held-out split are different protocols, and Table S11 against Table S12 bounds that difference jointly with the penalty and split rather than isolating it.

The capacity ladder in Table S13 runs from a zero-shot likelihood ratio through linear and two-layer probes to low-rank adaptation. Each step changes one thing. Because the low-rank step also changes the objective from linear readout to end-to-end adaptation, it should be read as the end of the ladder rather than as a probe setting.

*Relation to main text.* Results, *Gene hold-out collapses the whole panel to a band around chance*, run-in *Does either family collapse less than the other?*; Discussion (limits: frozen embeddings and a linear probe); Methods (probe).

*Comparisons that are licit across these tables.* Comparisons within one table, since the protocol is fixed there. Comparisons across tables only where the differing factor is named and the rest is matched. Comparisons that are not licit: reading a number from Table S11 against a number from Table S12 as if only the epoch rule had changed, and reading any single-split table against the folded tables in S3.

*Minimum detectable effect.* Under the folded design the paired minimum detectable effect at $\alpha = 0.05$ and power 0.80 is 9.3 percentage points, which is larger than several effects reported in this literature. Effects below that floor are not evidence of absence, and the tables in this section should not be read as establishing a null.

### S7. Statistical design

*Contents.* Effect bounds, three uncertainty types, cluster bootstrap, evidence classification rules, partition robustness of the family and support analyses.

#### Supplementary Table S31: MDES analysis

*Purpose.* Computes the minimum detectable effect size (MDES) [11] for paired ΔAUC comparisons under LOGO-CV (240 folds, $n = 2840$), to determine whether the literature's reported synonym-randomization drop (5.2 pp) is detectable at this design's resolution.
*Key observations.* The MDES at $\alpha = 0.05$, power$= 0.80$ is 9.3 pp for paired ΔAUC. The observed drop is 2.3 pp (paired $p = 0.488$). The literature's 5.2 pp falls *below* the detection floor, meaning the design cannot reliably distinguish it from fold variance. The conclusion upgrades from "no signal found" to "the literature's effect size is below the design's detectable threshold."
*Relation to main text.* Abstract (MDES = 9.3 pp); Results, *The dependence is on the specific gene, and part of every comparison is the protocol.*

| **Metric** | **Value** |
|---|---|
| Number of folds | 240 |
| Observed ΔAUC mean | 2.3 pp |
| SD of ΔAUC | 51.1 pp |
| SE of ΔAUC | 3.3 pp |
| Paired $t$ | 0.694 |
| Paired $p$ | 0.488 |
| **MDES** ($\alpha = 0.05$, power$= 0.80$) | **9.3 pp** |
| Literature reported drop | 5.2 pp |
| Detectable? | No |

**Per-gene AUC MDES** ($n = 20$ genes):

| **Model** | **Mean AUC** | **SD (pp)** | **SE (pp)** | **MDES (pp)** |
|---|---|---|---|---|
| onehot_pos | 0.730 | 15.1 | 3.4 | 10.0 |
| codonbert | 0.424 | 22.0 | 4.9 | 14.5 |
| codonbert_hf | 0.572 | 15.7 | 3.5 | 10.3 |
| encodon-80m | 0.543 | 21.1 | 4.7 | 13.9 |

#### Supplementary Table S29: Attribute regression ($R^2$)

*Purpose.* Tests whether cLM embeddings encode four attributes—GC3 content, Codon Adaptation Index (CAI), normalized variant position, and pathogenicity label—via ordinary least-squares linear regression with 5-fold CV.
*Key observations.* GC3 is highly recoverable ($R^2 = 0.72$–$0.95$) and CAI moderately so ($R^2 = 0.06$–$0.59$), confirming that cLM representations encode sequence statistics. Position and pathogenicity are not linearly recoverable ($R^2 < 0$), confirming that cLMs do not encode pathogenicity directly—the pathogenicity signal must be extracted by a probe, and even then does not survive leakage-controlled evaluation.
*Relation to main text.* Methods, *Statistical inference*; Results, *Gene hold-out collapses the whole panel to a band around chance.*

| **Model** | **GC3** | **CAI** | **Position** | **Pathogenicity** |
|---|---|---|---|---|
| CodonBERT | 0.9480 | 0.5944 | −0.7261 | −0.6569 |
| CodonBERT-HF | 0.7217 | 0.0634 | −0.9732 | −0.6939 |
| EnCodon-80M | 0.9395 | 0.4820 | −1.8990 | −1.7359 |

#### Supplementary Table S21: Codon-optimality independence

*Purpose.* Tests whether cLM embeddings encode codon-usage bias (GC3, CAI) as a pretraining feature, which could confound the pathogenicity prediction.
*Key observations.* GC3 is highly recoverable from embeddings ($R^2$ = 0.72–0.95) and CAI moderately so ($R^2$ = 0.06–0.59), confirming that cLMs encode sequence statistics. However, pathogenicity is not linearly recoverable ($R^2 < 0$; see Table S29), so codon-usage features do not directly encode pathogenicity.
*Relation to main text.* Results, *Gene hold-out collapses the whole panel to a band around chance*; Methods, *Statistical inference.*

| **Model** | **GC3** $R^2$ | **CAI** $R^2$ |
|---|---|---|
| CodonBERT | 0.9480 | 0.5944 |
| CodonBERT-HF | 0.7217 | 0.0634 |
| EnCodon-80M | 0.9395 | 0.4820 |

**[S7] Statistical design: units, uncertainty, and evidence rules.**
*Purpose.* State the unit of replication, the three kinds of uncertainty that arise in this design, and the rules used to decide when a comparison counts as resolved.
*Key observations.* The unit of replication in this study is the gene, not the variant. Variants from one gene share sequence composition, label prevalence and clinical attention, so treating them as independent observations understates the interval on any comparison by a wide margin. All intervals reported here therefore come from a bootstrap that resamples genes with replacement, using 1,000 replicates. Under this scheme the half-width is about 0.034 for the positional comparison and about 0.046 for the family comparison, which is why an effect of 0.015 cannot be ranked and why the family-level question is left unresolved rather than answered in the negative.

Three kinds of uncertainty need to be kept apart. Sampling uncertainty is what the bootstrap measures and is the only one it measures. Protocol uncertainty arises from choices such as the probe, the split, and the epoch rule, and it is larger here than sampling uncertainty; it is addressed by reporting the same quantity under several protocols rather than by widening an interval. Model uncertainty, meaning variation across representations, is what the panel in Table S39 is for.
*Relation to main text.* Results (all intervals); Discussion (four limits); Methods (probe and controls).

**Table S44: Uncertainty types, what addresses each, and what is not addressed.**

| Type | Addressed by | Not addressed by |
|---|---|---|
| Sampling (genes) | gene-clustered bootstrap, 1,000 replicates; half-width $\approx$ 0.034 positional, $\approx$ 0.046 family | widening the interval further; more variants from the same genes |
| Protocol (probe, split, epoch) | re-reporting the same quantity under each protocol; Tables S11, S12, S20, S22–S28 | the bootstrap, which holds the protocol fixed by construction |
| Model (representation) | the panel of 24 distinct representations; Table S39 | any single-model interval |
| Annotation | build-wise matching; exclusions counted in Table S43 | pooling assemblies, which was observed to drive the measure to chance |

*Evidence rules applied when reading these results.*

1. An interval that excludes zero is treated as resolved only when the interval is computed on the gene-clustered bootstrap. An interval that excludes zero under variant-level resampling is not reported.

2. An interval that contains zero is read as unresolved, not as a demonstration that the difference is absent.

3. A difference smaller than the half-width for its comparison is not ranked, even when it is positive and even when it is the largest in a panel.

4. An arm whose value is constant by construction is reported as undefined rather than as a number near chance, because a constant score carries no information and its ROC is undefined.

5. Fold-level statistics are reported alongside pooled figures but never used as the primary estimate, because folds here hold a median of five variants.

6. Excluded records are counted and reported with their label fraction, so that a reader can see what the exclusions do to the evaluated population.

**Table S46: Partition robustness for the family and support analyses.**
*Purpose.* Two arms use gene-level partitions rather than the leave-one-gene-out folds, because the contrasts they measure are not definable under a fold that removes every variant of the gene it holds out (Methods, *Hold-out unit in the family and support analyses*). This table shows that neither conclusion depends on the particular partition drawn, by repeating each analysis under a second, independent partition.
*Key observations.* Every comparison retains its direction and its interval. All six models keep support above exclude, with an interval that excludes zero, in both partitions; the largest change in a paired difference is 0.013 against effects of 0.13 to 0.22. In the family analysis the known-gene arm stays well above the unseen-gene arm, the family arm stays level with the unseen-gene arm, and the family arm does not fall below the matched diversity control in either partition.
*Why the two partitions are not identical by construction.* The ten-fold partition holds out 10% of genes per fold instead of 20% per repetition. Fewer genes are therefore removed as family members, the size-matched training set is larger, and the spread between folds is wider. Both are direct consequences of the design change and are visible in the table; neither reverses a comparison.

**(a) Same-gene support contrast, two random seeds**

| | Seed 42 | | | Seed 2026 | | |
|---|---|---|---|---|---|---|
| **Model** | **Support** | **Exclude** | **Δ** | **Support** | **Exclude** | **Δ** |
| CodonBERT | 0.758 | 0.624 | +0.134 | 0.747 | 0.613 | +0.133 |
| CodonBERT-HF | 0.707 | 0.562 | +0.144 | 0.699 | 0.545 | +0.154 |
| EnCodon-80M | 0.716 | 0.529 | +0.187 | 0.725 | 0.542 | +0.183 |
| mRNABERT | 0.721 | 0.561 | +0.160 | 0.691 | 0.545 | +0.147 |
| ESM-2-650M | 0.755 | 0.546 | +0.209 | 0.770 | 0.552 | +0.218 |
| ESM-1b-650M | 0.763 | 0.546 | +0.217 | 0.746 | 0.539 | +0.208 |

**(b) Family analysis, five 20% partitions against ten 10% partitions**

| | Five partitions (20%) | | | | Ten folds (10%) | | | |
|---|---|---|---|---|---|---|---|---|
| **Model** | **G1** | **G2** | **G3** | **G3−ctrl** | **G1** | **G2** | **G3** | **G3−ctrl** |
| CodonBERT | 0.701 | 0.560 | 0.551 | +0.001 | 0.713 | 0.576 | 0.562 | −0.012 |
| CodonBERT-HF | 0.673 | 0.561 | 0.571 | +0.034 | 0.692 | 0.543 | 0.554 | +0.014 |
| EnCodon-80M | 0.685 | 0.528 | 0.547 | +0.028 | 0.712 | 0.537 | 0.534 | +0.001 |
| mRNABERT | 0.659 | 0.518 | 0.531 | +0.017 | 0.680 | 0.543 | 0.564 | +0.018 |
| ESM-2-650M | 0.705 | 0.543 | 0.528 | −0.023 | 0.719 | 0.545 | 0.527 | −0.027 |
| ESM-1b-650M | 0.697 | 0.538 | 0.551 | +0.030 | 0.712 | 0.553 | 0.555 | +0.001 |

*Note.* G1 = known gene, G2 = unseen gene, G3 = unseen family, G3−ctrl = family arm minus a control that removes the same number of training genes at random. Cells are means over the partitions of each scheme; the between-fold spread widens under the ten-fold scheme for the reason given above, so the two columns of each block are not estimates of the same quantity at the same precision.
*Relation to main text.* Results, *The dependence is on the specific gene, and part of every comparison is the protocol*; Results, *The dependence is on the specific gene, and part of every comparison is the protocol*; Methods (hold-out unit in the family and support analyses).

#### S8. External confirmation

*Contents.* Saturation genome editing data, review-status stratification.

#### Supplementary Table S32: Position-only baseline two-arm experiment

*Purpose.* Independent validation of the leakage diagnosis on external data (saturation mutagenesis [12]), using a falsifiable directional prediction: if 0.666 reflects a generalizable biological position prior, the same positional-identity feature should retain discriminative power above 0.5 on external data under gene-held-out evaluation.
*Key observations.* Both arms show random-split AUC well above 0.5 (0.64 and 0.63) but held-out AUC ≈ 0.5 (0.50 and 0.51). Positional information does not generalize to held-out genes: no gene fold in either arm exceeds 0.56. The prediction is falsified: 0.666 reflects ClinVar's specific labeling and reporting structure, not a generalizable biological position prior. This independently confirms that the positional prior does not carry to unseen genes. It does not extend to the representation panel, which was not run on these data; see the note below.
*Relation to main text.* Abstract (5,966 synonymous / 26,783 CDS SNV variants); Results, *Does the pattern depend on curated labels?*; Discussion (audit checklist).
*Limits recorded.* Both arms use a positional feature only, so this experiment validates the positional arm and not the representation panel, which was never evaluated on these data. The six genes are all DNA-damage or cancer genes, so they are not a genome-wide sample. The coding arm's pooled gene-held-out value, 0.476, sits below chance by an amount we have not explained, and we report it rather than account for it. The binarisation rule for the functional score is fixed before splitting and is stated in the configuration above.
**Datasets:**

| Dataset | Total variants | Synonymous | CDS SNV | License |
|---|---|---|---|---|
| RAD51C SGE | 9,188 | 767 | 3,285 | CC BY 4.0 |
| BAP1 SGE | 18,108 | 1,543 | 6,501 | CC BY 4.0 |
| BARD1 SGE | 10,899 | 1,345 | 6,339 | CC0 |
| BRCA2 SGE | 6,959 | 1,345 | 6,270 | CC0 |
| BRCA1 SGE | 3,893 | 579 | 2,803 | CC0 |
| VHL SGE | 2,268 | 387 | 1,585 | CC BY 4.0 |
| **Total** | **51,315** | **5,966** | **26,783** | |

**Source publications:** RAD51C [13], BAP1 [14], BRCA2 [15], BRCA1 [16], VHL [17]. BARD1 data from MaveDB [12] (no linked publication).
**Arm 1: Synonymous variants, 6-gene pooled** ($n$ = 5,966, positive class 36.8%):

| Evaluation | AUC |
|---|---|
| Random split (5 repeats) | $0.6418 \pm 0.0103$ |
| Gene-held-out per-fold mean | $0.5165 \pm 0.0194$ |
| **Gene-held-out pooled** | **0.5025** |

**Arm 2: All CDS SNV, 6-gene pooled** ($n$ = 26,783, positive class 53.4%):
*Arm 2 design note.* Arm 2 was revised on 2026-09-29. The earlier version took RAD51C alone with uniform 50 position bins and a front/back position-held-out split; because training then covered only the lower position bins and testing only the upper ones, the two sides had disjoint

| Evaluation | AUC |
|---|---|
| Random split (5 repeats) | $0.6262 \pm 0.0061$ |
| Gene-held-out per-fold mean | $0.5059 \pm 0.0285$ |
| **Gene-held-out pooled** | **0.4764** |

one-hot support and the held-out AUC sat near chance by construction. The revised arm pools all CDS SNVs across the six genes, uses 50 quantile bins whose edges are fixed on the pooled data before splitting, and is evaluated gene-held-out, matching Arm 1. Per-gene folds: BAP1 0.4821, BARD1 0.4983, BRCA1 0.5572, BRCA2 0.4878, RAD51C 0.5303, VHL 0.4796.
**Falsifiable prediction result:** Both arms yield held-out AUC $\approx 0.5$, falsifying the prediction that 0.666 reflects a generalizable biological position prior. The 0.666 retained under LOGO-CV on ClinVar reflects ClinVar's specific labeling and reporting structure, not a biological position prior that transfers to external data.

#### Supplementary Figure S1: Review-status stratification

*Purpose.* Tests whether the depth signature (LR→MLP gain on SynPath) is driven by low-confidence ClinVar variants, by recomputing LR performance within each review-status tier.
*Key observations.* The depth signature is present within each tier, confirming that it is not driven by low-confidence variants. This is one of four controls validating the linear-probing interpretation. For CodonBERT-HF, LR AUC is 0.725 in Tier 4 ($n = 200$; 100/100) against 0.591 in the Tier 3 bulk ($n = 2{,}640$; 1,320/1,320), with a far wider fold-to-fold spread in Tier 4 (s.d. 0.117 vs 0.018 over five folds); the gap does not survive gene-held-out evaluation.
*Relation to main text.* Results, *Gene hold-out collapses the whole panel to a band around chance*; Methods, *Data.*

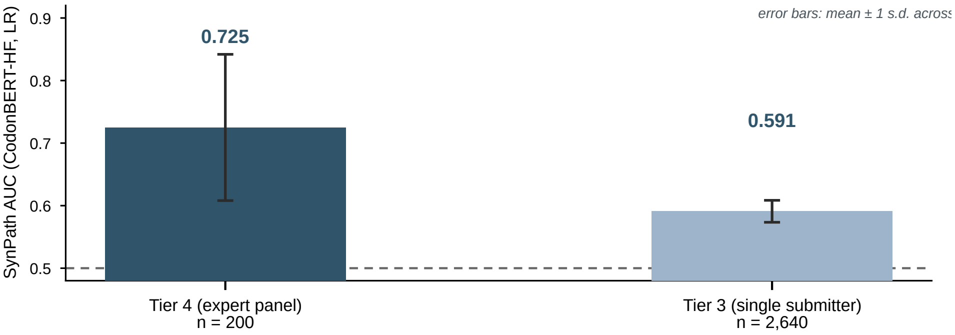


Figure 1: **ClinVar review-status stratification.** LR performance on SynPath and MisPath within each review-status tier (CodonBERT-HF: Tier 4, $n = 200$, AUC 0.725; Tier 3, $n = 2{,}640$, AUC 0.591). The depth signature is not driven by low-confidence variants.

### S9. Protocol evolution and correction record

*Contents.* Discovered anomalies, excluded errors, retracted results, unverified hypotheses, experiments that ultimately support conclusions; reflects research maturity.

#### Supplementary Note S1: Pooling and standardization corrections

*Purpose.* Documents two implementation issues identified and corrected during development, illustrating the protocol-sensitivity that CodonBench is designed to control.
*(1) ESM-1b mean-pooling bug.* An initial ESM-1b baseline used mean pooling that averaged over padding positions, spuriously depressing its SynPath AUC by 10.5 pp (0.600 $\rightarrow$ 0.705). The corrected implementation uses attention-masked mean pooling, excluding BOS/EOS and padding tokens. This correction is documented here as illustrative of the protocol-sensitivity this benchmark is designed to control.
*(2) StandardScaler effect.* Applying `StandardScaler` to the same mean-pooled ESM-2 features raises SynPath LR AUC from 0.685 to 0.762 (+7.7 pp). Standardization is itself a protocol choice, not a neutral default. All main-text claims use the model-specific pooling, no-scaler protocol of Supplementary Table S2; the unified-pipeline results in Supplementary Table S7 use model-specific pooling with standardization and are reported for cross-pipeline comparison only.
*Relation to main text.* Methods, *Probe*; the StandardScaler effect is referenced in Results, *Gene hold-out collapses the whole panel to a band around chance* as the first artifact—a preprocessing choice that flips the Level-1 lead.

#### Supplementary Table S33: Reproduction of the original protocol, and collapse under gene-held-out evaluation

*Purpose.* Two questions that must not be conflated, and that rest on different runs, so they are kept in two panels. Panel A asks whether our implementation reproduces the values published under the original random-split protocol. Panel B asks what is left of the random-split advantage once gene identity is held out. Every cell of Panel B comes from a *single* run, so its two numeric columns share the fold definition, the classifier and the preprocessing; no Panel B row mixes runs.
*Key observations.* Panel A: the only two models for which a published value on this task could be located (CodonBERT and ESM-2-650M) are reproduced to within 0.0004 AUC, so the practice of "reproduce first, then re-measure" is satisfied for those rows. The remaining rows carry --- under *Reported* because no value was published on SynPath; they are our measurements only, and no reproduction claim is made for them. Panel B: the collapse is not a property of pretrained codon models. A zero-pretraining (position, codon) lookup table drops 21.7 pp, more than 12 of the 15 pretrained models in the full comparison (§ 2.2, Supplementary Table S2), while low-dimensional free statistics move by less than 5 pp and GC content *gains* 8.2 pp. The magnitude of the drop therefore tracks how well a representation can fingerprint a gene, not whether it was pretrained.
*Relation to main text.* Results, *Gene hold-out collapses the whole panel to a band around chance*; Background, "Why a common task set, and why these tasks"; Discussion (protocol fragility).

**Panel A — reproduction fidelity.** Protocol: random split, no feature standardization (the original protocol).

| Model (readout) | Reported | Our re-run | Difference |
|---|---:|---:|---:|
| CodonBERT (LR) | 0.734 | 0.7336 | −0.0004 |
| CodonBERT (MLP) | 0.849 | 0.8486 | −0.0004 |
| EnCodon-80M (LR) | — | 0.6856 | — |
| ESM-2-650M (LR) | 0.685 | 0.6851 | +0.0001 |
| onehot_pos (LR, position-only baseline) | — | 0.8835 | — |

*Row provenance (Panel A).*

- CodonBERT (LR), EnCodon-80M (LR), onehot_pos (LR): results/comprehensive_comparison.json → <model>/task3_synonymous/auc_mean (5-fold random split). For onehot_pos this reproduces the published random-split value 0.884 (the single 80/20 split gives the same value to three decimals).
- CodonBERT (MLP): results/supplementary/synonym_randomization_v2_all_results.json → codonbert/synonymous/original_test_mlp (single 80/20 test split, which is the protocol under which 0.849 was reported). This row is *not* comparable to the 5-fold rows.
- ESM-2-650M (LR): results/esm2_task3.json → ROC-AUC_mean.
- *Reported* values: CodonBERT (0.734 LR, 0.849 MLP) and ESM-2-650M (0.685) are the values stated in the original publications, as previously recorded in this table. No value for EnCodon-80M on this task could be located (---), and onehot_pos is our own diagnostic baseline, never reported by anyone.

**Panel B — same-source collapse.** Protocol (identical for both columns, one run): StandardScaler fitted inside each training fold, LogisticRegression ($C = 1$, max_iter

= 1000, lbfgs); *Random* = StratifiedKFold(5, shuffle, seed 42); *Gene-held-out* = 240-fold leave-one-transcript-out (all variants of the held-out transcript removed together), per-fold mean.

| Representation (readout) | Random | Gene-held-out | Difference (pp) |
|---|---:|---:|---:|
| CodonBERT (LR) | 0.7249 | 0.5638 | −16.1 |
| CodonBERT-HF (LR) | 0.6945 | 0.5372 | −15.7 |
| EnCodon-80M (LR) | 0.6979 | 0.5370 | −16.1 |
| ESM-2-650M (LR) | 0.7426 | 0.5097 | −23.3 |
| onehot_pos (LR, position-only baseline) | 0.8835 | 0.6661 | −21.7 |
| codon composition (reference) | 0.5887 | 0.5469 | −4.2 |
| *k*-mer3 (reference) | 0.5725 | 0.5541 | −1.8 |
| GC content (reference) | 0.4675 | 0.5493 | +8.2 |

*Row provenance (Panel B).* All eight rows come from one run: results/leakage_control_2026-09-24/leakage_2x2.json, keys models/codonbert, models/codonbert_hf, models/encodon-80m, models/ESM-2-650M, baselines/onehot_pos, baselines/onehot_freq, baselines/kmer3, baselines/gc_content; fields A_perfold_mean (Random) and B_perfold_mean (Gene-held-out). The onehot_pos row reads no sequence and uses no pretrained weights. Free-baseline feature code is imported from scripts/run_traditional_baselines.py, not reimplemented.
*Cross-checks.* (a) *Same run as the published ladder:* for all six models that appear in both, the *Random* column equals pretrained_unified_results.json → <model>_SynPath/lr_5fold_mean to within 0.0001, and the *Gene-held-out* column equals lr_logo_mean to within 0.0027 (CodonBERT exactly 0.5638); the panel is therefore on the published ladder rather than a parallel one. (b) *Free-baseline random-split values under the original no-standardization protocol reproduce the published ones exactly* (0.884, 0.589, 0.6999, 0.4675; four to six decimals), so the free-baseline features are the same features as before — only the fold unit changed.
*CodonBERT (MLP), same 240 folds:* gene-held-out 0.6106 (logo_cv_mlp_results.json → codonbert/logo_mlp_mean). No MLP random-split counterpart exists inside this run, and the value is therefore *not* entered as a Panel B row; the MLP ladder is reported separately in Supplementary Table S8.
*Notes.* (i) Panel A and Panel B use different preprocessing, so their *Random* columns differ slightly for the same model (e.g. CodonBERT: 0.7336 without standardization vs 0.7249 with in-fold standardization, 0.9 pp). This is expected and the two panels must not be read across. (ii) All values are per-fold means, not pooled AUCs; pooled values differ because per-fold AUCs over 240 small folds are biased downward. Gene-held-out pooled AUCs: CodonBERT 0.5758, CodonBERT-HF 0.5519, EnCodon-80M 0.5236, ESM-2-650M 0.5462, onehot_pos 0.6738, codon composition 0.5448, *k*-mer3 0.5441, GC content 0.3963. Random-split pooled AUCs (5-fold, $\Delta < 0.01$ pp from per-fold means): 0.8833 for onehot_pos, within ±0.01 pp of per-fold for all others. Pooled-vs-pooled drops (pp): CodonBERT −14.9, CodonBERT-HF −14.3, EnCodon-80M −17.4, ESM-2-650M −19.6, onehot_pos −20.9, codon composition −4.4, *k*-mer3 −2.8, GC content −7.1; per-fold-vs-per-fold drops are −16.1, −15.7, −16.1, −23.3, −21.7, −4.2, −1.8, +8.2. The collapse is therefore not an artefact of per-fold aggregation for any neural model or onehot_pos; GC content flips sign under pooled aggregation (per-fold +8.2 pp → pooled −7.1 pp), consistent with its status as a below-noise-floor baseline (main text, Analysis V). (iii) The gene-held-out column is a per-fold mean over folds whose test sets are often very small, hence the large across-fold standard deviations (reported in the source JSON, $\sigma \approx 0.36$); only comparisons within the same column are meaningful. (iv) "Reported" values for the previously published table are reused verbatim from that table and were not re-verified against the original

PDFs within this revision.

#### Supplementary Figure S2: Training curves

*Purpose.* MLM training curves for the five from-scratch ablation conditions (v1, v3a, v3b, v4, v2), confirming that all models converge and the tokenization ablation results are not confounded by under-training.

*Key observations.* All models converge smoothly; no early stopping was used, so the number of epochs is fixed by design. The v2 synthetic-CDS control converges similarly to real-CDS conditions.

*Relation to main text.* Methods, *Representation panel*; confirms that the from-scratch models are properly trained.

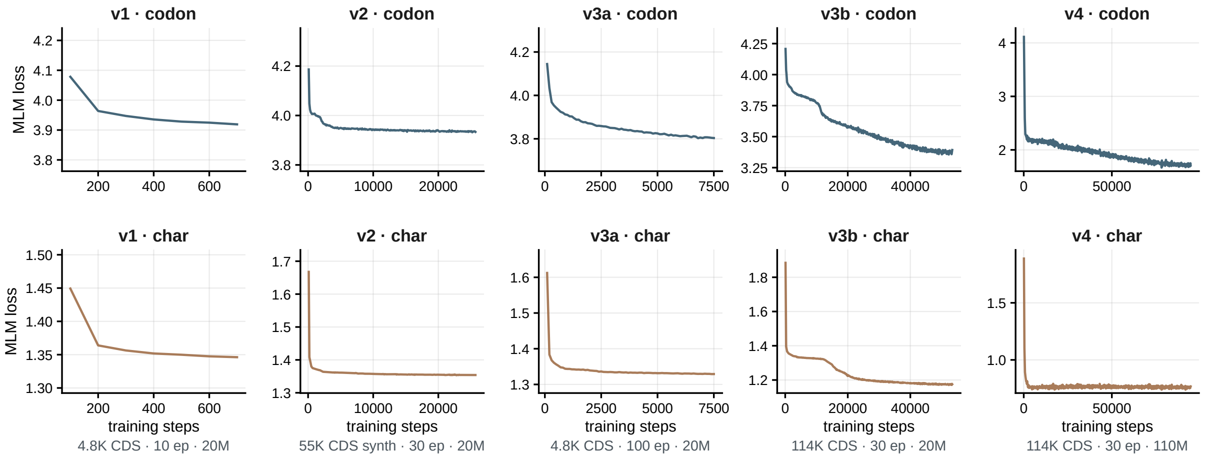


Figure 2: **MLM training curves for from-scratch models.** Loss vs. epoch for v1, v3a, v3b, v4, and v2, codon and character tokenizers. All models converge.

### Supplementary Figure S3: Layer-wise probing

*Purpose.* Visualizes the layer-wise probing results of Table S19 as curves, showing how LR and MLP AUC evolve across the 12 transformer layers of CodonBERT-HF on SynPath and MisPath. *Key observations.* On SynPath, both LR and MLP AUC increase monotonically through layer 12. On MisPath, AUCs increase through layer 11 then drop at layer 12, suggesting the final layer specializes for the MLM objective. There is no single layer where SynPath signal is concentrated. *Relation to main text.* Methods, *Statistical inference*; Supplementary Table S19.

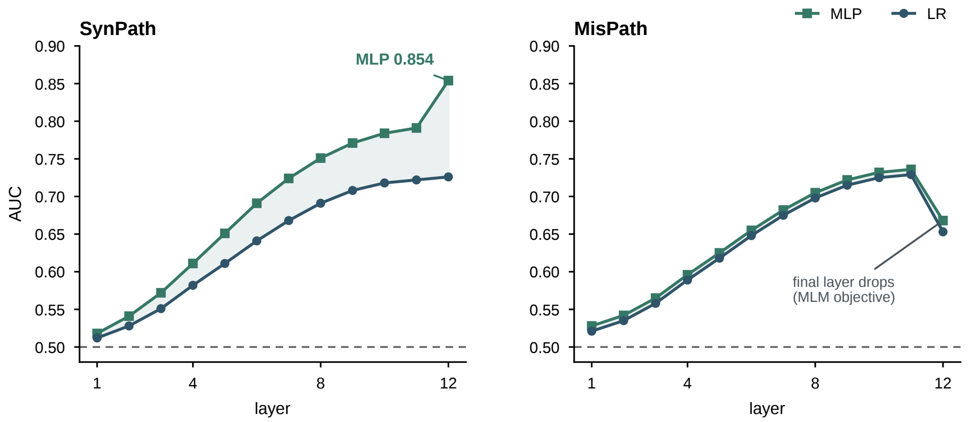


Figure 3: **Layer-wise probing of CodonBERT-HF.** LR and MLP AUC on SynPath and MisPath for each of the 12 transformer layers.

#### Supplementary Methods

The main-text Methods section provides the protocol overview and key parameters. This section supplements it with implementation details that are referenced but not fully expanded there.

##### From-scratch ablation: detailed training configuration

All from-scratch models were trained using HuggingFace `BertForMaskedLM` with the following configuration:

**v1–v3b (∼20M parameters):**

- Architecture: 6 layers, hidden 512, intermediate 2048, 8 attention heads, max_position_embeddings = 512 for codon / 1536 for character
- Codon tokenizer: vocab = 69 (61 sense codons + 8 special tokens)
- Character tokenizer: vocab = 9 (A, T, G, C, N + 4 special tokens)
- MLM probability: 0.15
- Learning rate: $5 \times 10^{-5}$ with linear warmup over 10% of total steps then linear decay
- Weight decay: 0.01; AdamW ($\beta_1$=0.9, $\beta_2$=0.999); Batch size: 32; Dropout: 0.1
- No early stopping; all models trained for the specified number of epochs

**v4 (∼110M parameters):** 12 layers, hidden 768, intermediate 3072, 12 attention heads. Same tokenizer, MLM probability, and optimizer settings as v3b.

**Synthetic CDS generation (v2):** For each position in each real CDS, a codon was sampled from the per-position marginal frequency distribution estimated from the real CDS corpus. This preserves per-position codon usage frequencies but destroys inter-position codon co-occurrence patterns.

*Relation to main text.* Methods, *Representation panel*; the five conditions are listed there but architecture details are given here.

##### M0→M5 single-variable ablation: detailed configurations

Starting from a bare PyTorch MLP (M0), each configuration adds one conventional choice:

- **M0 (PyTorch baseline):** `nn.Sequential`(768→128→64→1, ReLU, Sigmoid), Adam optimizer (lr=1e-3), 100 epochs, no early stopping, no weight decay, no fixed seed, StandardScaler on inputs.
- **M1 (+seed):** M0 + `torch.manual_seed(42)` + `np.random.seed(42)` + `torch.cuda.manual_seed_all(42)`.
- **M2 (+early stopping):** M1 + early stopping on a 10% validation fraction of the training set (patience = 10 epochs, restoring best weights).
- **M3 (+weight decay):** M2 + weight_decay = 1e-4 in Adam optimizer.
- **M4 (+epochs):** M3 with max_epochs = 200 (already converged at 100; M4 gain is bit-identical to M3).
- **M5 (sklearn):** scikit-learn MLPClassifier with early_stopping = True, validation_fraction = 0.1, random_state = 42, max_iter = 200, alpha = 1e-4, solver = 'adam', StandardScaler.

All six configurations share the same data, fold splits, and LR probe; LR AUC is bit-identical across all configurations (0.5638), isolating the entire effect to the nonlinear probe.
*Relation to main text.* Results, *The dependence is on the specific gene, and part of every comparison is the protocol*; Methods, *Statistical inference*; Supplementary Table S22.

#### Seed sweep protocol

To characterize the seed sensitivity of the M0 configuration and contextualize the M0→M1 increment, the M4-configuration PyTorch MLP was run across 20 random seeds (0–19), holding all other parameters fixed. The same sweep was performed with the scikit-learn MLPClassifier. Both sweeps use the same LOGO-CV protocol (497 genes, 240 valid folds) as the M0→M5 ablation.
*Rationale.* The M0→M1 increment (+1.5 pp) appears to attribute a large effect to fixing the seed. However, M0 is a single unseeded run; the −0.6 pp gain it produces may simply be a low-end sample from the seed distribution. The seed sweep determines whether seed = 42 is exceptional (it is not within the PyTorch distribution; it exceeds all 20 sklearn seeds).
*Relation to main text.* Results, *The dependence is on the specific gene, and part of every comparison is the protocol*; Supplementary Tables S26, S27.

#### Model loading details

| Model | Source | Loading method | Required adaptations |
|---|---|---|---|
| CodonBERT | Local checkpoint | Manual vocab.txt | Vocab file created from training data |
| CodonBERT-HF | lhallee/CodonBERT (HF) | Standard AutoModel | T→U conversion; RNA codon vocabulary |
| EnCodon-80M | goodarzilab/encodon-80M (HF) | AutoModel with patches | einsum patch for rotary embeddings; xformers |
| EnCodon-620M | goodarzilab/encodon-620M (HF) | Same as EnCodon-80M | Same patches as EnCodon-80M |
| CodonTransformer | Aghayeg/codontransformer (HF) | Standard AutoModel | Dual-token $\{AA\}_{codon}$ scheme |
| CaLM | lulu-blu/CaLM (HF) | Documented adaptation | Contrastive pretraining; special token handling |
| cdsBERT | lhallee/cdsBERT (HF) | Standard AutoModel | Character-level tokenizer (vocab = 9) |
| cdsBERT-plus | lhallee/cdsBERT-plus (HF) | Standard AutoModel | Character-level tokenizer |
| mRNABERT | HuggingFace | AutoModel with config | Flash attention disabled; MosaicBERT arch |
| Mistral-Codon-117M/16M | HuggingFace | Standard AutoModel | Mixtral MoE architecture |
| ESM-2-650M | fair-esm/esm2-t33-650M | fair-esm package | AA tokenizer; mean pooling |
| ESM-1b-650M | fair-esm/esm1b_t12_85M | fair-esm package | No CLS token; mean pooling |
| NT-v2-500M | InstaDeepAI/nt-v2-500m (HF) | Standard AutoModel | BPE tokenizer (vocab = 4,096); 6-mer tokens |
| NT-v2-50M | InstaDeepAI/nt-v2-50m (HF) | Standard AutoModel | Same as NT-v2-500M |

*Relation to main text.* Methods, *Representation panel*; all embeddings extracted with dropout disabled, no gradient computation, batch size 8.

#### CodonBench-Agent: pre-flight methodological validation

CodonBench-Agent performs two pre-flight validation checks before each evaluation run:

**1. Gene-identity leakage check (`check_gene_leakage`).** Given a train/test split and a gene-to-sample mapping, the Agent checks whether any gene appears in both the training and test sets. If overlap is detected, the Agent raises a `LeakageError` and recommends using LOGO-CV splits.

**2. Best-epoch selection bias check (`check_epoch_selection`).** Given the epoch selection source (test set vs validation set), the Agent checks whether the probe epoch is selected on

the test set. If so, the Agent raises a `SelectionBiasError` and recommends validation-based early stopping.
*Relation to main text.* Methods (third contribution); the two pre-flight checks make controlled evaluation the path of least resistance.

**Disclosure of direction-of-effect asymmetries.** Two results in the frozen run point *against* the codon-advantage hypothesis rather than merely failing to support it. We report both in full because an audit that carried only the codon-favouring comparisons would be as selective as the literature it criticises.

1. *The linear probe tilts systematically toward character tokenization under LOGO-CV.* Per-fold-mean $\Delta$ (codon $-$ char) is $-3.1$, $-1.2$, $-4.1$ and $-3.8$ pp for v1/v3a/v3b/v4 (pooled: $-4.6$, $-0.6$, $-0.9$ and $-3.3$ pp), with the paired sign test approaching nominal significance in three of four conditions ($p = 0.086$, 0.099, 0.103; Table S25). The displacement is absent under the MLP probe. We build no claim on it: a linear read-out of a frozen representation is the least sensitive probe in our ladder, and the LR columns of Table S6 carry no seed sweep.

2. *The only nominally significant paired effect in the entire ablation is v2 under a PyTorch MLP.* Across all 15 condition–probe comparisons under LOGO-CV, the single significant result is the synthetic-CDS control v2 with a PyTorch MLP ($\Delta = -7.85$ pp, paired $t$ $p = 0.0169$; paired sign test $p = 0.0330$; 10-seed mean $-8.0 \pm 1.7$ pp; Tables S16, S25). The direction favours character tokenization, on a control that by construction carries no real inter-position codon structure. It therefore cannot support the codon-advantage hypothesis, and we do not interpret it as a positive discovery. The mechanism is not identified.

A third asymmetry is a single-run number retained for transparency: under the *random* 5-fold protocol the linear probe shows a +9.6 pp codon lead for v4 ($p = 0.003$) while the other three real-CDS conditions show +0.8 to +3.1 pp ($p \geq 0.097$; Table S15). Because no seed sweep was run for the linear probe, this number carries the same single-seed caveat that Section 2.3 of the main text raises for the MLP probe.

#### References

1. Sizhen Li, Saeed Moayedpour, Ruben Li, Michael Bailey, Saleh Riahi, et al. Codonbert large language model for mrna vaccines. *Genome Research*, 34(7):1027–1035, 2024. doi: 10.1101/gr.278870.123.

2. Mohsen Naghipourfar, Siyu Chen, Mary K. Howard, Christian B. Macdonald, Ali Saberi, Timothy Hagen, Mohammad R. K. Mofrad, Willow Coyote-Maestas, and Hani Goodarzi. A suite of foundation models captures the contextual interplay between codons. *bioRxiv*, 2024. doi: 10.1101/2024.10.10.617568.

3. Adibvafa Fallahpour et al. Codontransformer: a multispecies codon optimizer using context-aware neural networks. *Nature Communications*, 2025. doi: 10.1038/s41467-025-58588-7.

4. Carlos Outeiral and Charlotte M. Deane. Codon language embeddings provide strong signals for use in protein engineering. *Nature Machine Intelligence*, 2024. doi: 10.1038/s42256-024-00791-0.

5. Yi Xiong, An Wang, Yu Kang, Cheng Shen, Cheng-Yu Hsieh, and Tingjun Hou. mrnabert: advancing mrna sequence design with a universal language model and comprehensive dataset. *Nature Communications*, 2025. doi: 10.1038/s41467-025-65340-8.

6. Logan Hallee, Nikolaos Rafailidis, and Jason P. Gleghorn. cdsbert — extending protein language models with codon awareness. *bioRxiv*, 2023. doi: 10.1101/2023.09.15.558027.

7. Zeming Lin, Halil Akin, Roshan Rao, Brian Hie, Zhongkai Zhu, Wenting Lu, Nikita Smetanin, Robert Verkuil, Ori Kabeli, Yaniv Shmueli, et al. Evolutionary-scale prediction of atomic-level protein structure with a language model. *Science*, 379(6637):1123–1130, 2023. doi: 10.1126/science.ade2574. [CROSSREF] Title, journal, volume, issue, pages and the 15 authors verified against the DOI record on 2026-10-05; identical to the entry previously carried in the supplementary file.

8. Alexander Rives, Joshua Meier, Tom Sercu, Siddharth Goyal, Zeming Lin, et al. Biological structure and function emerge from scaling unsupervised learning to 250 million protein sequences. *Proceedings of the National Academy of Sciences*, 118(15):e2016239118, 2021. doi: 10.1073/pnas.2016239118.

9. Hugo Dalla-Torre et al. Nucleotide transformer: building and evaluating robust foundation models for human genomics. *Nature Methods*, 2025. doi: 10.1038/s41592-024-02523-z.

10. Jun Cheng, Guido Novati, Joshua Pan, Clare Bycroft, Akvilė Zemgulyte, Tom Applebaum, Alexander Pritzel, Lai Hong Wong, et al. Accurate proteome-wide missense variant effect prediction with AlphaMissense. *Science*, 381(6664):eadg7492, 2023. doi: 10.1126/science.adg7492. [CROSSREF] Title, journal, volume, issue and article number verified against the DOI record on 2026-10-05; identical to the entry previously carried in the supplementary file.

11. Jacob Cohen. *Statistical Power Analysis for the Behavioral Sciences*. Lawrence Erlbaum Associates, 2nd edition, 1988. [2026-10-05] Recovered from the supplementary file's own reference list, where it was entered as an article; changed here to book (Lawrence Erlbaum Associates, 2nd ed.), which is what the item actually is. The supplementary file no longer carries a reference list of its own.

12. Alan F. Rubin, Jeremy Stone, Alice H. Bianchi, Benjamin J. Capodanno, et al. Mavedb 2024: a curated community database with over seven million variant effects from multiplexed functional assays. *Genome Biology*, 2025. doi: 10.1186/s13059-025-03476-y.

13. Rebeca Olvera-León, Fang Zhang, Victoria Offord, Yuhan Zhao, Huei San Tan, et al. High-resolution functional mapping of rad51c by saturation genome editing. *Cell*, 2024. doi: 10.1016/j.cell.2024.08.039.

14. Andrew J. Waters, Tobias Brendler-Spaeth, Danielle Smith, Victoria Offord, Huei San Tan, et al. Saturation genome editing of bap1 functionally classifies somatic and germline variants. *Nature Genetics*, 2024. doi: 10.1038/s41588-024-01799-3.

15. Hongyuan Huang, Chunling Hu, Jie Na, Steven N. Hart, Rohan D. Gnanaolivu, et al. Functional evaluation and clinical classification of brca2 variants. *Nature*, 2025. doi: 10.1038/s41586-024-08388-8.

16. Gregory M. Findlay, Riza M. Daza, Brotman Martin, Melissa D. Zhang, Anh P. Leith, and Jay Shendure. Accurate classification of brca1 variants with saturation genome editing. *Nature*, 2018. doi: 10.1038/s41586-018-0461-z.

17. Max Buckley, Céline Terwagne, Andrea Ganner, Laura Cubitt, Rebecca Brewer, et al. Saturation genome editing maps the functional spectrum of pathogenic vhl alleles. *Nature Genetics*, 2024. doi: 10.1038/s41588-024-01800-z.

#### S10. Tables moved from the main text

*Contents.* The eight tables below carry the numeric results that the main text reports in figures. Each is preceded by an experiment card giving the name of the experiment, its configuration, the data it uses, and its result.

##### Table S47

**Experiment.** Panel under the two splitting regimes
*Configuration.* 24 frozen representations; logistic-regression probe ($C = 1$, L-BFGS); features standardised on the training folds only; identical folds and identical samples in both regimes.
*Data.* 1,847 variants; leave-one-gene-out ($D_2$, 240 folds) against random stratified splitting ($D_1$, five folds).
*Result.* Every representation contracts by 0.10–0.26 AUC; under gene hold-out none exceeds 0.577 and the codon and protein families occupy the same band.
*Relation to main text.* Results, *Gene hold-out collapses the whole panel to a band around chance.*
**Table S47: Frozen representations under the two splitting regimes**

| Representation | Dim. | $D_1$ random | $D_2$ gene-held-out |
|---|---|---|---|
| CodonTransformer | 768 | 0.749 | 0.577 |
| CodonBERT | 768 | 0.737 | 0.561 |
| CaLM | 768 | 0.738 | 0.553 |
| cdsBERT-plus | 1024 | 0.724 | 0.548 |
| ESM-1b-650M (mean pool) | 1280 | 0.772 | 0.546 |
| ESM-2 (protein windows) | 1280 | 0.752 | 0.543 |
| ESM-2-650M | 1280 | 0.755 | 0.543 |
| Nucleotide Transformer v2 500M | 1024 | 0.738 | 0.543 |
| from-scratch v4 | 768 | 0.718 | 0.541 |
| EnCodon-620M | 2048 | 0.790 | 0.537 |
| cdsBERT | 1024 | 0.694 | 0.533 |
| EnCodon-620M (layer 1) | 2048 | 0.790 | 0.533 |
| ESM-2-650M (mean pool) | 1280 | 0.752 | 0.532 |
| ESM-1b-650M | 1280 | 0.779 | 0.529 |
| mRNABERT | 768 | 0.709 | 0.527 |
| Nucleotide Transformer v2 50M | 512 | 0.671 | 0.524 |
| CodonBERT (mean pool) | 768 | 0.682 | 0.523 |
| EnCodon-80M | 1024 | 0.723 | 0.520 |
| Mistral-Codon-117M | 768 | 0.695 | 0.517 |
| CodonBERT-HF | 768 | 0.720 | 0.509 |
| from-scratch v3b (ctx 128) | 512 | 0.652 | 0.506 |
| Mistral-Codon-16M | 256 | 0.610 | 0.505 |
| from-scratch v4 (ctx 128) | 768 | 0.680 | 0.500 |
| from-scratch v3b | 512 | 0.638 | 0.488 |

##### Table S48

**Experiment.** Pre-modelling identity checks on the two families
*Configuration.* Mean absolute difference between the reference and alternate embeddings over all dimensions; then the same control comparison used for the codon models applied to a protein model.
*Data.* CodonBERT and ESM-1b-650M embeddings of the evaluated variants.

*Result.* Codon model $3.9 \times 10^{-2}$ (must differ); protein model 0.000 (must be identical); Δ against a position-matched synonymous control 0.000 (interval of zero width); Δ against a protein-altering control −0.242 (95% CI −0.294 to −0.184).
*Relation to main text.* Results, *Gene hold-out collapses the whole panel to a band around chance.*
**Table S48: Pre-modelling checks on the two model families**

| Model | Check | Mean \|ref−alt\| |
|---|---|---|
| CodonBERT (cLM) | must differ | $3.9 \times 10^{-2}$ |
| ESM-1b-650M (pLM) | must be identical | 0.000 |

The same protein model was then run through the control comparison: Δ against a position-matched synonymous control is exactly 0.000 (95% CI 0.000 to 0.000), while Δ against a control that alters the protein is −0.242 (95% CI −0.294 to −0.184), $n = 1{,}837$.

#### Table S49

**Experiment.** Family-level paired comparison
*Configuration.* Eight paired codon-minus-protein comparisons on a shared test set under gene hold-out; intervals from a gene-clustered bootstrap.
*Data.* $n = 1{,}847$.
*Result.* None of the eight differences separates from zero; the intervals have a half-width near 0.046, so the comparison is unresolved rather than null.
*Relation to main text.* Results, *Gene hold-out collapses the whole panel to a band around chance.*
**Table S49: Paired codon-model minus protein-model differences under gene hold-out**

| Comparison | Δ (AUC) | 95% CI |
|---|---|---|
| CodonBERT − ESM-2-650M | +0.026 | −0.020 to +0.072 |
| CodonBERT − ESM-1b-650M | +0.039 | −0.003 to +0.084 |
| CodonBERT-HF − ESM-2-650M | −0.002 | −0.047 to +0.046 |
| CodonBERT-HF − ESM-1b-650M | +0.011 | −0.031 to +0.060 |
| EnCodon-80M − ESM-2-650M | −0.030 | −0.074 to +0.015 |
| EnCodon-80M − ESM-1b-650M | −0.016 | −0.059 to +0.031 |
| mRNABERT − ESM-2-650M | +0.001 | −0.039 to +0.044 |
| mRNABERT − ESM-1b-650M | +0.014 | −0.027 to +0.065 |

#### Table S50

**Experiment.** Exon-edge enrichment
*Configuration.* Counts of variants lying on the first or last base of an exon, and within 3 nt of a junction; no model fitted.
*Data.* $n = 2{,}814$ records with a matched transcript structure.
*Result.* Pathogenic 54.9% against benign 0.07%; within 3 nt, 64.6% against 3.3%; the excess is spread over 248 of 495 genes rather than concentrated in a few.
*Relation to main text.* Results, *The strongest single source is position, and it reads no sequence.*
**Table S50: Position of each variant relative to an exon junction**

#### Table S51

**Experiment.** Positional baselines

| | Pathogenic | Benign |
|---|---|---|
| On the first or last base of an exon | 54.9% | 0.07% |
| Within 3 nt of a junction | 64.6% | 3.3% |

*Configuration.* Distance to the nearest exon junction and to the nearest CDS terminus, negated and used directly as a score; no model fitted and no sequence read. A label-permuted arm is the negative control.
*Data.* $n = 1{,}847$; 238 of 240 folds usable.
*Result.* Exon-junction distance 0.843 (fold-level mean 0.854); CDS-terminus distance 0.534; label-permuted 0.497.
*Relation to main text.* Results, *The strongest single source is position, and it reads no sequence.*
**Table S51: Positional baselines under the leave-one-gene-out folds**

| Score | Pooled OOF AUC | Fold-level mean |
|---|---|---|
| Distance to nearest exon junction | **0.843** | 0.854 |
| Distance to nearest CDS terminus | 0.534 | 0.474 |
| Distance to junction, labels permuted | 0.497 | — |

#### Table S53

**Experiment.** Removing whole homologous families
*Configuration.* Three arms: G1 a test gene whose other variants remain in training; G2 the test gene removed; G3 additionally every training gene sharing an HGNC family removed, with a control removing the same number of genes at random; means over five independent gene partitions.
*Data.* Training sample size matched across arms.
*Result.* The step from G1 to G2 costs 0.11–0.16 AUC in every model (0.112 for CodonBERT-HF, up to 0.162 for ESM-2-650M); the further step to G3 is within ±0.02 and does not agree in sign across models.
*Relation to main text.* Results, *The dependence is on the specific gene, and part of every comparison is the protocol.*
**Table S53: Removing whole homologous families, with a size-matched control**

| Model | Known | Unseen | Family | Family − control |
|---|---|---|---|---|
| CodonBERT | 0.701 | 0.560 | 0.551 | $+0.001 \pm 0.037$ |
| CodonBERT-HF | 0.673 | 0.561 | 0.571 | $+0.034 \pm 0.027$ |
| EnCodon-80M | 0.685 | 0.528 | 0.547 | $+0.028 \pm 0.028$ |
| mRNABERT | 0.659 | 0.518 | 0.531 | $+0.017 \pm 0.029$ |
| ESM-2-650M | 0.705 | 0.543 | 0.528 | $-0.023 \pm 0.016$ |
| ESM-1b-650M | 0.697 | 0.538 | 0.551 | $+0.030 \pm 0.051$ |

#### Table S54

**Experiment.** Same-gene training support
*Configuration.* For each gene carrying more than one variant, one variant is held out and training is repeated with and without that gene's remaining variants; training-set size matched across arms; intervals gene-clustered.
*Data.* Test set fixed at 489 variants, one per gene.

*Result.* Support effects from +0.134 to +0.217; significant for every model and, if anything, larger for the protein models than for the codon models.
*Relation to main text.* Results, *The dependence is on the specific gene, and part of every comparison is the protocol.*
**Table S54: Training support from other variants of the held-out gene**

| Model | Support | Exclude | Δ | 95% CI |
|---|---|---|---|---|
| CodonBERT | 0.758 | 0.624 | +0.134 | 0.117 to 0.153 |
| CodonBERT-HF | 0.706 | 0.562 | +0.144 | 0.125 to 0.163 |
| EnCodon-80M | 0.716 | 0.529 | +0.187 | 0.163 to 0.209 |
| mRNABERT | 0.721 | 0.561 | +0.160 | 0.138 to 0.181 |
| ESM-2-650M | 0.755 | 0.546 | +0.209 | 0.179 to 0.242 |
| ESM-1b-650M | 0.763 | 0.545 | +0.217 | 0.182 to 0.252 |

#### Table S55

**Experiment.** Where the variant-specific component comes from
*Configuration.* The response vector (alternate minus reference embedding) against the position-matched control, decomposed in four ways and run under two sample scopes. Scope A uses every evaluated variant. Scope B uses only the variants whose control was actually constructed; the difference between the scopes is stated in the text. Feature sets: the full response vector; its direction after normalising to unit length; its magnitude alone; and the residual after regressing out magnitude and simple sequence features within each fold. Same folds, same probe, gene-clustered bootstrap, 1,000 replicates.
*Data.* $n$ = 1,839 in scope A and $n$ = 1,209 in scope B; 240 folds usable in scope A and 230 in scope B.
*Result.* Against the position-matched control the difference is +0.138 (+0.101, +0.176) in cdsBERT and +0.145 (+0.108, +0.183) in cdsBERT-plus under scope A, and +0.064 (+0.017, +0.106) and +0.100 (+0.053, +0.146) under scope B; the direction alone reproduces it and the residual does not.
*Relation to main text.* Results, *A variant-specific component survives both controls*; Methods, *Controls* and *Variant-specific component.*
**Table S55: Decomposition of the variant-specific component**
*A note on the magnitude row.* The magnitude comparison is reported for completeness and is not interpreted in the main text. Under scope B the magnitude of the control arm predicts in the opposite direction (AUC 0.318 in cdsBERT, 0.412 in cdsBERT-plus), so the large difference on that row is produced mainly by the control arm rather than by the response vector.
*A note on the residual row.* The residual is formed by regressing the magnitude and the simple sequence features out of the response vector within each fold. Doing so drives both arms below chance (AUC 0.399 and 0.386 for the response and control arms of cdsBERT under scope B), which is the expected consequence of projecting a high-dimensional vector into the space orthogonal to a small feature set rather than evidence about the component. The row is reported and not built upon; Table S58 asks the same question in increment form, which does not have this property.

#### Table S58

**Experiment.** Increment form of the decomposition
*Configuration.* The response vector is added on top of a stated baseline rather than projected against it, and the accuracy gained is compared between the real arm and the control arm. Three baselines are used, each nested in the next: position alone; position plus the simple sequence

| Scope and feature set | | cdsBERT | cdsBERT-plus |
|---|---|---|---|
| *Scope A: every evaluated variant* ($n$ = 1,839) | | | |
| Response vector | AUC | 0.667 | 0.671 |
| Position-matched control | AUC | 0.529 | 0.526 |
| Direction only, response vector | AUC | 0.662 | 0.672 |
| Direction only, control | AUC | 0.532 | 0.526 |
| Magnitude only, response vector | AUC | 0.598 | 0.645 |
| Magnitude only, control | AUC | 0.518 | 0.525 |
| Δ, full response vector | | +0.138 (+0.101, +0.176) | +0.145 (+0.108, +0.183) |
| Δ, direction only | | +0.130 (+0.088, +0.171) | +0.146 (+0.106, +0.185) |
| Δ, magnitude only | | +0.080 (+0.040, +0.118) | +0.120 (+0.081, +0.159) |
| Δ, residual after simple features | | -0.048 (-0.080, -0.012) | -0.042 (-0.082, -0.004) |
| *Scope B: control actually constructed* ($n$ = 1,209) | | | |
| Response vector | AUC | 0.666 | 0.682 |
| Position-matched control | AUC | 0.602 | 0.582 |
| Direction only, response vector | AUC | 0.666 | 0.683 |
| Direction only, control | AUC | 0.602 | 0.580 |
| Magnitude only, response vector | AUC | 0.520 | 0.592 |
| Magnitude only, control | AUC | 0.318 | 0.412 |
| Δ, full response vector | | +0.064 (+0.017, +0.106) | +0.100 (+0.053, +0.146) |
| Δ, direction only | | +0.064 (+0.019, +0.110) | +0.103 (+0.058, +0.147) |
| Δ, magnitude only | | +0.202 (+0.136, +0.262) | +0.180 (+0.126, +0.238) |
| Δ, residual after simple features | | +0.014 (-0.027, +0.054) | +0.007 (-0.033, +0.048) |

features; and those plus the magnitude of the response vector. Same folds, same probe, gene-clustered bootstrap, 1,000 replicates.
*What the simple sequence features are.* They are computed from the sequences alone, with no model, and they already include the quantity most likely to mimic the effect under test. For each record they are: the signed change in the relative frequency of the codon within its own synonymous family between the reference and the substituted codon, the absolute value of that change, the corresponding change in the codon's rank within the family, the size of the family, the frequencies of the two codons taken separately, the change in whether the substituted base is G or C, whether the substitution is a transition, and a one-hot encoding of which base was replaced by which. The first item is the codon-usage change the substitution itself produces, so the middle row answers directly whether the response vector adds anything beyond it; the last item means the exact nucleotide substitution is also already accounted for.
*Data.* $n$ = 1,839 in scope A and $n$ = 1,209 in scope B.
*Relation to main text.* Results, *A directional variant-specific component survives both controls.*
**Table S58: Increment of the response vector over stated baselines**

#### Table S56

**Experiment.** Strata by codon-usage change
*Configuration.* Variants are grouped by the sign of the change in synonymous-codon frequency caused by the real substitution. Frequencies are relative within a synonymous family and are counted over the 500 transcripts of the package's own CDS table (782,719 codons), so no external codon-usage table is used. The two arms are compared within each stratum by the same folds and the same probe.
*Data.* Same folds as Table S55.

| Baseline above which the response vector is added | Scope | cdsBERT | cdsBERT-plus |
|---|---|---|---|
| None | A | +0.138 (+0.101, +0.176) | +0.145 (+0.108, +0.183) |
| Position | A | +0.041 (+0.006, +0.076) | +0.059 (+0.021, +0.093) |
| Position + simple sequence features | A | +0.040 (+0.009, +0.075) | +0.057 (+0.022, +0.088) |
| Those + magnitude | A | +0.021 (-0.010, +0.054) | +0.030 (-0.005, +0.060) |
| None | B | +0.064 (+0.017, +0.106) | +0.100 (+0.053, +0.146) |
| Position | B | +0.063 (+0.019, +0.104) | +0.091 (+0.047, +0.133) |
| Position + simple sequence features | B | +0.070 (+0.031, +0.108) | +0.075 (+0.034, +0.116) |
| Those + magnitude | B | +0.063 (+0.025, +0.101) | +0.071 (+0.030, +0.112) |

*Relation to main text.* Results, *A variant-specific component survives both controls.*
**Table S56: The component by stratum**

| Stratum | Model | $n$ | Response | Control | Δ (95% CI) |
|---|---|---|---|---|---|
| *Scope A* | | | | | |
| Rarer codon | cdsBERT | 1133 | 0.672 | 0.492 | +0.180 (+0.132, +0.227) |
| More common codon | cdsBERT | 547 | 0.663 | 0.634 | +0.030 (-0.038, +0.095) |
| Frequency unchanged | cdsBERT | 159 | 0.519 | 0.489 | +0.030 (-0.133, +0.181) |
| Rarer codon | cdsBERT-plus | 1133 | 0.658 | 0.510 | +0.148 (+0.094, +0.201) |
| More common codon | cdsBERT-plus | 547 | 0.697 | 0.589 | +0.107 (+0.035, +0.179) |
| Frequency unchanged | cdsBERT-plus | 159 | 0.613 | 0.595 | +0.018 (-0.141, +0.181) |
| *Scope B* | | | | | |
| Rarer codon | cdsBERT | 661 | 0.653 | 0.530 | +0.123 (+0.057, +0.191) |
| More common codon | cdsBERT | 451 | 0.697 | 0.679 | +0.018 (-0.047, +0.088) |
| Frequency unchanged | cdsBERT | 97 | 0.571 | 0.592 | -0.021 (-0.160, +0.120) |
| Rarer codon | cdsBERT-plus | 661 | 0.660 | 0.559 | +0.101 (+0.040, +0.161) |
| More common codon | cdsBERT-plus | 451 | 0.721 | 0.597 | +0.123 (+0.046, +0.201) |
| Frequency unchanged | cdsBERT-plus | 97 | 0.620 | 0.623 | -0.002 (-0.155, +0.147) |

#### Table S57

**Experiment.** Checks on the comparison: coverage, comparability, rank and negative control
*Configuration.* (i) Coverage of the position-matched control; (ii) comparability of the two arms on simple sequence features that can be computed without a model; (iii) rank decomposition, retaining the first $k$ principal components of the response vector; (iv) label-permuted negative control. Codon frequencies as in Table S56.
*Data.* $n = 1{,}839$; the control is the zero vector for 630 of them (34.3%).
*Relation to main text.* Results, *A variant-specific component survives both controls.*
**Table S57: Coverage, comparability and checks**
**Table S57b: Rank decomposition of the response vector (cdsBERT, scope A)**

#### Table S59

**Experiment.** A third control, matched on nucleotide chemistry as well as position

| Quantity | Real substitution | Control substitution |
|---|---|---|
| *Coverage* | | |
| Records with a constructed control | 1209 of 1839 (65.7%) | |
| Records whose control is the zero vector | 630 (34.3%) | |
| Median $\|\Delta\|$, all records | 0.142 | 0.117 |
| Median $\|\Delta\|$, control constructed | 0.142 | 0.165 |
| *Comparability of the two arms* | | |
| Mean change in synonymous-codon frequency | -0.053 | -0.010 |
| Mean GC change at the substituted base | -0.457 | -0.207 |
| Proportion of transitions | 0.765 | 0.152 |
| *Negative control, response vector* | | |
| Labels permuted, seed 7 | 0.499 / 0.482 | |
| Labels permuted, seed 2026 | 0.472 / 0.501 | |

| $k$ | AUC | $k$ | AUC | $k$ | AUC |
|---|---|---|---|---|---|
| 1 | 0.454 | 2 | 0.550 | 4 | 0.620 |
| 8 | 0.692 | 16 | 0.717 | 32 | 0.728 |
| 64 | 0.740 | 128 | 0.734 | 256 | 0.680 |
| 512 | 0.628 | 1024 | 0.556 | | |

*Configuration.* The position-matched synonymous control replaces a variant's own codon with a different synonymous codon at the same position inside the codon (Methods, *Controls*). It does not match the chemistry of the substitution: a transition (purine to purine, or pyrimidine to pyrimidine) in the real arm can be paired with a transversion in the control arm, and the two arms differ sharply on this feature (Table S57). The chemistry-matched control adds one constraint to the same construction: the substituted base must belong to the same chemical class as the real substitution, so the control reproduces whether the change is a transition or a transversion. Everything else is unchanged: the same window, the same folds, the same probe, the same aggregation rule (per-fold mean over the 240 leave-one-gene-out folds) and a gene-clustered bootstrap with 1,000 replicates. Where no codon satisfies the added constraint the control is not built and the reference sequence is returned, exactly as in the position-matched case, so the procedure cannot favour the real arm.
*Data.* $n = 1{,}839$ evaluated variants, with 239 of the 240 folds usable. At the construction stage 973 of the 2,840 input records admit a chemistry-matched control, a coverage of 34.3%.
*Result.* Against the chemistry-matched control the difference is +0.238 (95% CI 0.200 to 0.274) in cdsBERT and +0.234 (0.188 to 0.275) in cdsBERT-plus. Both are larger than the corresponding differences against the position-matched control, and both exclude zero. The reason the gap widens rather than narrows is that the stricter control scores lower, not higher: the chemistry-matched control arm sits below chance, at a pooled AUC of 0.429 and 0.437 in the two models. On the subset where the control could be built the two arms agree on transition rate record by record, 36.9% against 36.9%, which is the check that the added constraint took effect; the position-matched control fails the same check. One limit follows from how the control is built. Transitions are harder to match than transversions, because a transition already uses up both bases of its own class at that position, so the constructable subset is skewed touching this edge: the comparison is therefore a paired one inside that subset and is not a statement about the

full set. We do not read the wider gap as stronger support for the component. A control arm that sits below chance is itself unexplained, so what this row establishes is that the difference is sensitive to how the control is built, and nothing beyond that.

*Relation to main text.* Results, *A directional variant-specific component survives both controls*; Discussion, sixth limit.

**Table S59: The response vector against a control matched on nucleotide chemistry**

| Model | Position-matched control | | Chemistry-matched control | |
|---|---|---|---|---|
| | Δ AUC | 95% CI | Δ AUC | 95% CI |
| cdsBERT | +0.170 | 0.126 to 0.214 | +0.238 | 0.200 to 0.274 |
| cdsBERT-plus | +0.166 | 0.123 to 0.208 | +0.234 | 0.188 to 0.275 |

**Table S59b: Whether each control matches the real arm on substitution chemistry**

| **Quantity** | **Real arm** | **Control arm** |
|---|---|---|
| *Position-matched control, every evaluated record (Table S57)* | | |
| Proportion of transitions | 0.765 | 0.152 |
| *Chemistry-matched control, records that actually substitute a base* | | |
| Real arm, all such records | 0.776 | |
| Control arm, all such records | 0.369 | |
| *Self-check on the subset where the control was built* | | |
| Real arm | 0.369 | — |
| Control arm | — | 0.369 |
| Records with a constructed control | 973 of 2,840 (34.3%) | |

The two blocks are not on the same denominator, and the difference is the point. The first counts every evaluated record, including those whose control could not be built; such a control returns the reference sequence, makes no substitution, and so counts as a non-transition. The second counts only records in which the arm in question actually substitutes a base. It is the second rule that makes the self-check meaningful: read on the whole set, the chemistry-matched control looks as if it had failed to match, when in fact it matches record by record on the only subset where it exists.
